# Adaptation to complex environments is associated with higher fitness costs under benign conditions

**DOI:** 10.64898/2025.12.22.695988

**Authors:** Giacomo Zilio, Emilie Aubin, Stéphanie Bedhomme, Laurence Bolick, Ignacio G. Bravo, Claude Bruand, Delphine Capela, Marie Challe, Guillaume M. Charrière, Gaelle Courtay, Marie-Ange Devillez, Fiona Elmaleh, Stanislas Fereol, Rémy Froissart, Justina Givens, Claire Gougat-Barbera, Lynn Govaert, Alice Guidot, Jeanne Hamet, Michèle Huet, Philipp Hummer, Staffan Jacob, Oliver Kaltz, Marc Krasovec, Delphine Legrand, Guillaume Martin, Thibault Nidelet, Denis Orcel, Hervé Philippe, Gwenael Piganeau, M. Stefania Przybylska, Philippe Remigi, Laurent Sauviac, Fabian Schneider-Nettsträter, Diego Segond, Celine Serre, Delphine Sicard, João Gabriel Colares Silveira, Jeanne Tonnabel, François Vasseur, Apolline Vedrenne, Edith Vidal, Cyrille Violle, Marielle Kathrin Wenzel, Emanuel A. Fronhofer, Luis-Miguel Chevin

## Abstract

Complex environments with simultaneous change in multiple stressors is the new norm in ecosystems worldwide, and adaptive evolution will be critical for population persistence under these combined environmental challenges. Yet, how the selective complexity imposed by such stressor combinations affects the pace of evolution remains poorly understood. Using a coordinated experiment, we exposed 14 species, from bacteria to unicellular eukaryotes and plants, to single stressors versus their pairwise combinations over multiple generations, while keeping the overall stress level comparable. We find that populations evolving under combined stressors tended to have lower fitness increase, and a shallower relation between initial maladaptation and fitness gain, than populations evolving under single stressors, although this direct response depended strongly on stressor type. When assayed in the control, benign conditions, lines evolved under combined stressors had higher fitness reductions than those evolved under single stressors, with higher consistency between species than for direct evolutionary responses. Our unique coordinated experimental evolution approach provides new insights into how environmental complexity can impact the pace of evolution, together with revealing the diversity of these responses. It further highlights the importance of incorporating evolutionary dynamics into the combined-stressor research agenda.

**Significance Statement:** Understanding how organisms respond to environmental stressors is a long-standing problem in biology, made more urgent as global change increasingly exposes species to combined stressors. We hypothesised that adaptation would be slower under combined stressors, because evolution then needs to solve multiple biological problems at once. We tested this prediction by experimentally evolving 14 organisms under different combinations of stressors. We found a weak tendency for combined stressors to constrain adaptation relative to single stressors, but a much stronger and consistent additional cost for combined stressors through reduced fitness under benign conditions. Accounting for such constraints should prove crucial for the management of biodiversity.

## Introduction

Natural populations live in multifaceted and complex environments, often characterised by the concurrent variation of multiple stressors, which reduce fitness by causing higher mortality or reduced reproduction (1). Anthropogenic activity is one important source of stress, imposing changes in temperatures (2) and salinity (3), but also drought or pollution. Hence, environmental change combining several stressors is the new norm in ecosystems worldwide (4–7). Understanding the impacts of such combined stressors on organismal performance and ecological outcomes is therefore a central goal for both basic and applied research. Most evidence for responses to combined stressors comes from short-term ecological studies. Ecotoxicology in particular provides an important framework for investigating how combinations of stressors interact on a short timescale (8–10). For instance, when co-occurring stressors mobilize the same physiological process with limited energetic budget, synergistic (negative) effects on an organism performance might be predicted (11), but are not always found (12, 13). However, these ecological studies tend to focus on within- or trans-generational plasticity (14, 15), and ignore genetic evolution. As the speed of adaptive evolution can determine how effectively organisms will be able to persist under changing environments (16), it is crucial to investigate how rapidly species can evolve in response to complex environments combining several stressors, as has been investigated in specific contexts such as multidrug treatments in bacteria (17).

Evolutionary theory has long recognized that rates of adaptation can be reduced by a cost of complexity. The main reason for this cost is that mutations that simultaneously solve a combination of biological challenges are expected to be less common than those favoured under simpler selective pressures (18, 19). Consistent with this expectation, pioneering work on the application of multidrug treatments has indeed shown that combinations of antibiotics can limit the evolution of resistance in bacterial pathogens (20–22). Similarly, exposure of the chlorophyte *Chlamydomonas reinhardtii* to a mixture of herbicides constrained the adaptive evolution of resistance (23). Abundant empirical evidence also supports that trade-offs limit an organism’s performance and fitness across environments, through mechanisms such as antagonistic pleiotropy, genetic correlations, epistasis, and resource limitation (24, 25). For example, the pleiotropy of mutations fixed under selection increased with increasing environmental complexity in *Saccharomyces cerevisiae* (26). On the other hand, selection for more biological functions under combined stressors may also mean that beneficial mutations overcoming these environmental challenges may arise at more places along the genome. This broader genomic target may accelerate adaptation by increasing the overall supply of advantageous mutations, though this possibility was not considered in classic theory on the cost of complexity (18, 19).

Adapting to environmental challenges may also incur an additional cost: a fitness reduction in environments where selection did not occur (27, 28), as exemplified by costs of resistance to antibiotics or pesticides (29). Although such fitness costs in non-selective environments (*i.e.*, indirect responses to selection) are not always quantified and considered (30), they may constrain future evolutionary responses, if the selective environment later changes (31, 32). While some empirical evidence supports costs of adaptation in non-selective environments (25), other experiments show no relationships with adaptation in the selective environment, or even positive correlated responses (33–36). More importantly, whether or not such costs differ between populations that evolved in response to simple *vs.* combined stressors remains little understood.

Although a few notable evolutionary experiments have investigated how adaptation proceeds in response to single *vs*. combined stressors (26, 37–41), we still lack consistent and quantitative evidence on this question. As each of these studies used somewhat different methodologies and metrics, especially regarding whether and how the intensity of stress and strength of selection were controlled, drawing general conclusions is difficult. A powerful alternative is to use a coordinated experiment, where an analogous experimental protocol is shared across systems, thereby overcoming the idiosyncrasies of specific experimental setups or laboratory conditions, enabling broader and more robust inference to be drawn. Such coordinated experiments have been successfully applied in ecology, to study notably dispersal and plant productivity (42–44). Here, we used this approach in an evolutionary context, to experimentally contrast adaptive evolution in response to simple *vs.* combined stressors.

## Results

Our coordinated experiment involved 12 research groups applying an analogous protocol across 14 species (**Table 1**), ranging from aquatic and terrestrial bacteria to unicellular eukaryotes (ciliates, Euglenozoa, chlorophytes, fungi) and plants, with one or more strains each (in total 23 strains, *i.e.* lineages characterized by distinct ancestral genetic backgrounds; Supplementary Text for details), and at least three replicated evolutionary lines per strain and treatment. For most species, the evolutionary lines started from an isogenic or monoclonal genetic background, such that the genetic responses to selection mainly relied on *de novo* mutations. The only exceptions were the two plant species *Arabidopsis thaliana* and *Brassica rapa*, and one experimental group of *Dunaliella salina* (details in SI), which were instead designed to maximize the contribution from standing variation. For *A. thaliana*, we used recombinant F2s from random crosses between 358 wild accessions spanning a large range of genetic, geographic, and phenotypic variation. For *B. rapa*, the starting population was similarly derived from a broad sampling of natural populations that subsequently underwent artificial selection for one generation. Lastly, one experimental group of the microalga *D. salina* was created by initially mixing in equal proportions the four strains used in the monoclonal lines. We expected no sexual reproduction to take place between these strains, such that the mixed-strain group evolved under clonal competition.

**Table 1.** List of species used in the coordinated experiment, with biological and experimental detail. The indirect response (IR) column indicates whether the species was tested in control conditions after evolution. In the stressor identity column, T indicates Temperature. In the column reporting the final replicates, Ctrl, S, and C stand for Control, Simple and Complex stressor, respectively. Unless specified, the different strains within species experienced the same experimental conditions and replication. The dilution rate provides information on population growth within each cycle; assuming demographic replacement on average, the population is multiplied by the inverse of the dilution rate in each cycle. Dilution rate is not applicable for plants (NA), and (*) denotes a dilution rate inferred from population density before transfer (see also SI). Number of generations was calculated as the number of transfers * log2 of dilution rate, which assumes a binary fission with negligible death (applicable to all the species in the study apart from the two plants), and represents a conservative estimate assuming that the population at least replaces itself over the transfers.

| Species | Group | Strains | N° of transfers | Dilution rate | Time (months) | N° of generations | Fitness estimate | IR | Stressor identity | Final replicates per evolutionary treatment |
|---|---|---|---|---|---|---|---|---|---|---|
| *Arabidopsis thaliana* | Plants | 1 | 5 | NA | 60 | 5 | Visual counting | Yes | Resources, Herbivory | Ctrl x7, S x14, C x7 |
| *Brassica rapa* | Plants | 1 | 3 | NA | 12 | 3 | Visual counting | No | Resources, Shade | Ctrl x5, S x10, C x5 |
| *Colpidium striatum* | Ciliates | 1 | 17 | 1/5 | 12 | 39 | Computer vision | Yes | T, Salt, Copper | Ctrl x3, S x8, C x5 |
| *Dunaliella salina*<br>(all strains) | Chlorophytes | 4 | 36 | 2/15 | 8 | 105 | Flow cytometry | Yes | T, Salt | Ctrl x3, S x6, C x3 |
| *Euglena gracilis* | Euglenozoan | 1 | 20 | 1/10 | 5 | 66 | Computer vision | Yes | T, Salt, Copper | Ctrl x10, S x14, C x14 |
| *Ostreococcus tauri* | Chlorophytes | 1 | 17 | ~ 1/100<sup>*</sup> | 4 | 113 | Flow cytometry | No | T, Salt | Ctrl x6, S x12, C x6 |
| *Paramecium caudatum* | Ciliates | 1 | 48 | 1/5 | 12 | 111 | Visual counting | Yes | T, Salt | Ctrl x5, S x7, C x3 |
| *Picochlorum costavermella* | Chlorophytes | 1 | 12 | ~ 1/20<sup>*</sup> | 3 | 52 | Flow cytometry | No | T, Salt | Ctrl x6, S x12, C x5 |
| *Pseudomonas aeruginosa* | Bacteria | 1 | 26 | 1/100 | ~1 | 173 | Optical density | Yes | Salt, Copper | Ctrl x12, S x20, C x9 |
| *Ralstonia solanacearum* | Bacteria | 1 | 15 | ~ 1/1000<sup>*</sup> | ~0.5 | 149 | Optical density | Yes | T, Salt | Ctrl x5, S x10, C x5 |
| *Saccharomyces cerevisiae*<br>(all strains) | Fungi | 2 | 26 | 1/5 | 3 | 60 | Optical density | Yes | T, Copper | Ctrl x3, S x6, C x3 |
| *Sinorhizobium meliloti* | Bacteria | 1 | 20 | ~ 1/1000<sup>*</sup> | ~0.66 | 200 | Optical density | Yes | T, Salt | Ctrl x5, S x10, C x5 |
| *Tetrahymena thermophila*<br>Strain T1<br>Strain D16<br>Strain D19<br>Strain D21<br>Strain D3 | Ciliates | 5 | 20 | 1/50 | ~5 | 113 | Computer vision and Optical density | Yes | T, Salt, Copper,<br>Salt, Antibiotic<br>Salt, Antibiotic<br>Salt, Antibiotic<br>Salt, Antibiotic | Ctrl x10, S x15, C x15<br>Ctrl x6, S x9, C x4<br>Ctrl x6, S x12, C x5<br>Ctrl x6, S x12, C x6<br>Ctrl x6, S x12, C x5 |
| *Vibrio aestuarianus*<br>Strain 12/016<br>Strain U29 | Bacteria | 2 | 22<br>16 | 1/100 | ~1.5<br>~0.25 | 146<br>106 | Optical density | Yes | Salt, Copper | Ctrl x3, S x6, C x3 |

For each species, we used two sources of stress, in four treatments: a control (absence of stressors); two simple stress treatments, with each single stressor applied separately; and a complex stress treatment combining the two single stressors. Stressor levels were chosen to be biologically relevant to species-specific stress responses (**Table 1**), and where feasible (in *Paramecium caudatum*, *Vibrio aestuarianus*, *Ralstonia solanacearum*, *Sinorhizobium meliloti*), were measured after a few generations or cycles of acclimation, to limit the immediate impact of the transition among environments (45). Importantly, we strived to adjust stressor levels in each system so that the initial fitness reduction under stress was comparable between simple and complex stressor treatments, to distinguish the role of the cost of complexity from that of the total magnitude of stress. Stressors were chosen among major threats to ecosystems worldwide, thought to exert strong selective pressures *in natura* on the respective focal taxa (**Table 1**). They comprised temperature, salt, copper, antibiotics, and resource deprivation (2, 46–48).

We assessed evolutionary responses by comparing demographic performance at the beginning and end of experimental evolution. For micro-organisms, we estimated the exponential growth rate *r*_0_ from growth curves (as in **Fig. 1**) obtained *via* optical density, flow cytometry, visual counting, or computer vision. For plants we instead used the logarithm of the expected number of seeds or fruits, which analogously quantifies changes in log-abundance, but per generation rather than per unit time (see Supplementary Text for more detail about estimation protocols). Log-abundance (or log-density) and time were standardised by centering and reducing (within each strain of each species, and across treatments), thereby enabling comparison among species with different growth rates. In all analyses we used linear mixed models (LMM), propagating the error associated with the fitness estimates. This leads to conservative results, and allows effectively weighting fitness effect sizes by study precision.

**Fig. 1.**
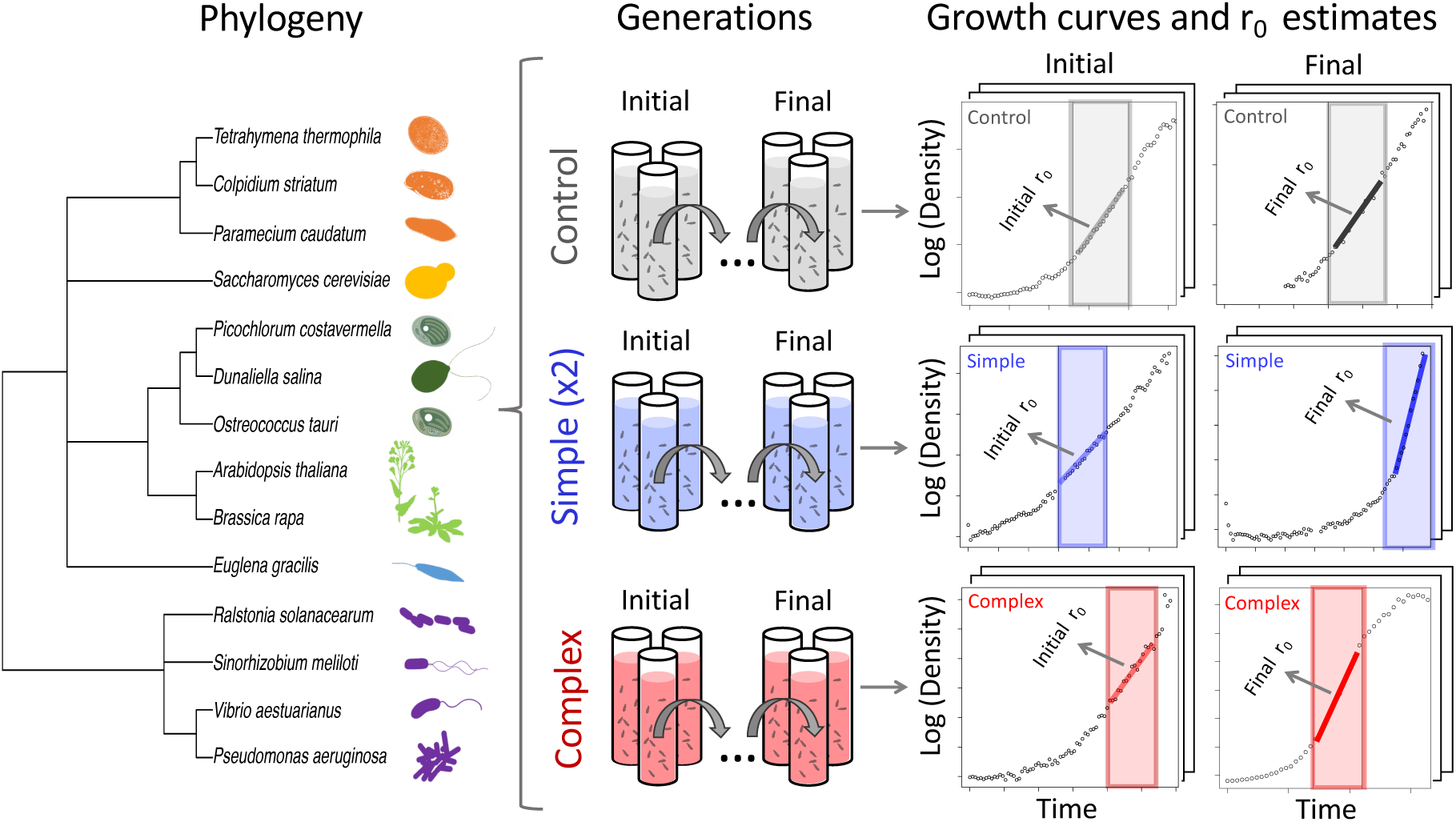
Experimental design. Schematic representation of the main procedures of the study, showing three evolutionary replicates per treatment for illustrative purpose. The model organisms (silhouettes in orange: protists; yellow: fungus; dark green: chlorophytes; light green: plants; light blue: Euglenozoa; purple: bacteria) were propagated by serial transfer (apart from plants, see SI) under 4 treatments: control (grey), simple stress (single stressors, blue), or complex stress (combined stressors, red). For all unicellular species (12/14), the steepest slope in the growth curve relating the logarithm of population density to time estimates the intrinsic growth rates *r*_0_, which was used as proxy for fitness in the main analyses. For plants (2/14 species), fitness estimates were instead obtained using life-history traits (e.g., fruits and seeds characteristics, see section SI). The difference between the final and initial fitness assayed in the evolutionary treatments yielded the direct evolutionary response. Final fitness was also estimated in the absence of stressors (control conditions) for 11 species, yielding an indirect evolutionary response, or cost of adaptation. Experimental protocols and additional details for each species are provided in the SI.

### The initial fitness drop was comparable between simple and complex stress

We initially asked whether the initial fitness drop caused by maladaptation was affected by the environmental conditions: control, single, or combined stressors. The organisms were assayed under the same conditions as those experienced during evolution. Whenever feasible, we allowed for some generations of acclimation before the first fitness measurement.

Initial exposure to environmental stressors reduced fitness by ∼55 % on average, compared to the control treatment (**Fig. 2A**, Linear Mixed-effect Model (LMM) in a Bayesian framework with “Species” and “Strain” as random effects, fitness median and 95% Compatibility Interval (CI): 1.140 [0.622, 1.555]; (49)). As intended by our protocol, single and combined stressors imposed similar fitness reductions relative to the control (LMM, Simple stress treatment: -0.533 [-0.715, - 0.345]; Complex: -0.742 [-0.951, -0.529]), as visualized by the largely overlapping distribution in the **Fig. 2A** inset. Separate analyses on each strain (linear models, **Table S1**) confirmed that the initial fitness reduction was not statistically different between single and combined stressors in 18 cases out of 23 (posteriors overlapping with zero, **Fig. S1**). Further, one of the five cases that did not overlap zero (*Vibrio aestuarianus* U29, **Fig. S1**) exhibited a larger initial drop in fitness under single stressor than under combined stressors, contrary to the expectation if we had not controlled the stressor levels in the treatment with combined stress. Thus, overall, our experimental protocol initially imposed comparable fitness reductions in treatments with simple *vs.* complex stress.

**Fig. 2.**
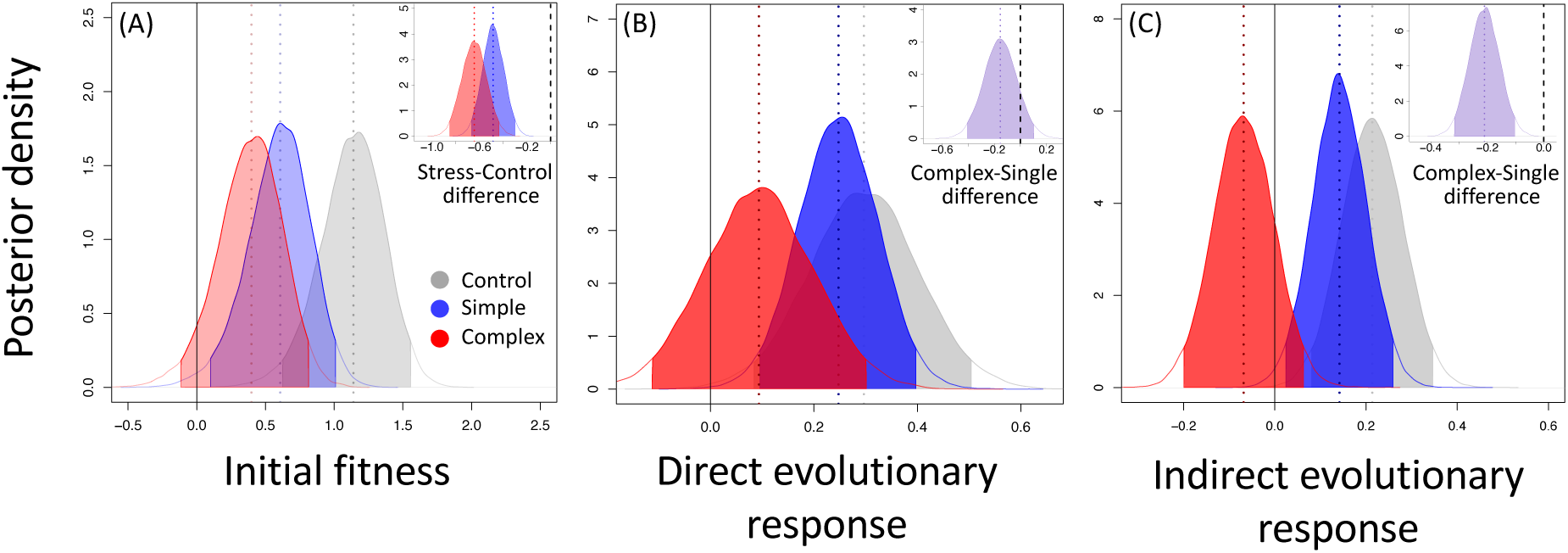
Initial fitness, direct and indirect responses. All fitness measurements were standardized by centering and reducing within strain. The dashed lines visualize the median posterior values, and the shaded areas show the posterior distributions within their 95% compatibility intervals. Grey represents the control in the absence of stressors, while blue and red show the single and combined stressors (simple vs complex stress), respectively. (A) Fitness following the initial transfer to the experimental conditions. (B) Direct response to selection, assayed in the conditions of experimental evolution, by calculating the difference final minus initial fitness for each treatment. (C) Indirect response to selection, obtained by calculating the difference between the final fitness of populations from the three evolutionary treatments when assayed under control conditions, and the initial fitness under control conditions. In all panels, the inset illustrates the posterior difference between combined and single stressor. The complete dataset (14 species and 23 strains) was used in (A) and (B), and a reduced dataset (11 species and 20 strains) in (C).

### Direct evolutionary response tended to be weaker under combined stressors, but variable among stressor types

After experimental evolution, fitness had increased in all treatments (**Fig. 2B**; see also **Fig. S2** with raw data on initial and final fitness for each strain), indicating that adaptive evolution had occurred. We quantified the direct evolutionary response (**Fig. 2B**) as the difference between final and initial fitness, for each treatment. Populations evolving under single stressors showed similar increases in fitness as those in the control treatment (LMM, Single and Control difference: 0.247 [0.095, 0.397], 0.296 [0.083, 0.504]; **Fig. 2B**), whereas the combined stressor treatment exhibited a less marked fitness gain than the other two (LMM, 0.093 [-0.113, 0.301]). Specifically, the combined stressor treatment tended to be associated with a smaller fitness increase than the single stressor treatment, although the difference was not strongly supported (LMM, Complex minus Single difference - 0.153 [-0.406, 0.103], **Fig. 2B** inset).

The effect of the evolutionary treatment alone explained 17% [0.124, 0.217] of the variance in direct evolutionary response (fixed effect only; marginal R^2^), while the full model including fixed and random effects (species and strains) explained 57% [0.532, 0.610] (conditional R^2^). This indicates a good model fit but substantial between-taxon variability, with a strong structuring of the data by strain and species (inter-specific variation explained∼ 2/3 of the total heterogeneity; I^2^ species = 57.42 % [9.83 %, 85.35 %], see also **Table S2**). When running separate analyses (linear models, **Table S1**) on each strain, the pattern of smaller fitness increase under combined *vs.* single stressors was found in 13 cases out of 23 (lower median values, **Fig. S3** and contrast in **Fig. S4**). Since the plants differed from microorganisms in the way we measured fitness, and (together with one experimental group of the microalga *D. salina*) in having started from inflated standing genetic variation, we also carried out two complementary analyses removing these organisms, which did not change our results qualitatively (**Fig. S5A, B**). Our results are also robust with regards to differences in duration of the individual experiments: Including the number of generations (Table 1) as a modulator in this and the other statistical models had no qualitative impact (**Fig. S6**).

### Complex environments were associated with higher indirect costs of adaptation

To investigate costs of adaptation, we then assayed the fitness of evolved populations under control environmental conditions (**Fig. S7**), for 11 out of 14 species for which these tests were logistically feasible (see “Indirect response” column in **Table 1**). We found that populations that evolved under simple stress showed an increase in fitness (difference between final and initial fitness) in control environmental conditions (LMM, Single difference: 0.141 [0.023,0.258]; **Fig. 2C**), comparable to that of control populations (LMM, Control: 0.213 [0.080,0.345]). This indicates that evolution in single stressors did not come at a cost for fitness in the absence of stress. In contrast, populations evolved under complex stress showed a tendency for decreased fitness under control environmental conditions (LMM, Complex: -0.068 [-0.200, 0.062], **Fig. 2C**). More importantly, they clearly differed from control-evolved populations (**Fig. 2C**, red *vs*. grey distribution: -0.282 [-0.404, - 0.159]), and from populations evolved under simple stress (LMM, Complex minus Single difference: -0.209 [-0.316, -0.102], **Fig. 2C** inset). The treatment alone explained a smaller fraction of the total variance (marginal R^2^: 7% [0.03, 0.126]) of the full model including also the random terms (conditional R^2^: 73% [0.686, 0.772], with substantial between-taxon variability and strong structuring of the data by strain and species. The inter-specific variation resembled that for direct response, explaining the majority of the biological differences across study systems (I^2^ = 76.92 % [31.83 %, 95.35 %]; **Table S2**). Nevertheless, separate linear models run on each strain (**Table S3**) consistently suggested the same general pattern of greater fitness decreases in the control treatment for populations from complex than simple stress (lower median values, **Fig. S8,** contrast in **Fig. S9**) in 16 cases out of 20.

### The relationship between direct and indirect responses varied among stressor types

Costs of adaptation can reflect a trade-off, such that fitness gain in one environment is accompanied by fitness loss in another (48). This is illustrated in **Fig. 3A**, which shows the overall relationship between the direct and indirect responses to selection in our experiment, together with the contrast between single and complex stress for each. To further explore how the nature of the stressors influences these responses, we conducted post-hoc analyses on data subsets associated with each stressor (**Fig. 3B-G**). For the direct response (*x*-axis in **Fig. 3B-G**), most stressors markedly differed from the overall pattern, with the notable exception of antibiotics (**Fig. 3B**, LMM Simple: 3.427 [2.855, 4.002]; Complex: 1.632 [1.138, 2.119]). As this stressor type also had the strongest signal, it may be largely driving the overall pattern for direct responses, despite having only been used in one system with four strains (*Tetrahymena spp*). Complementary analysis removing these four strains from the dataset show that the direct evolutionary response became comparable between simple and complex stresses, and the weak tendency for combined stressors to constrain adaptation relative to single stressors disappeared (**Fig. S10**).

**Fig. 3.**
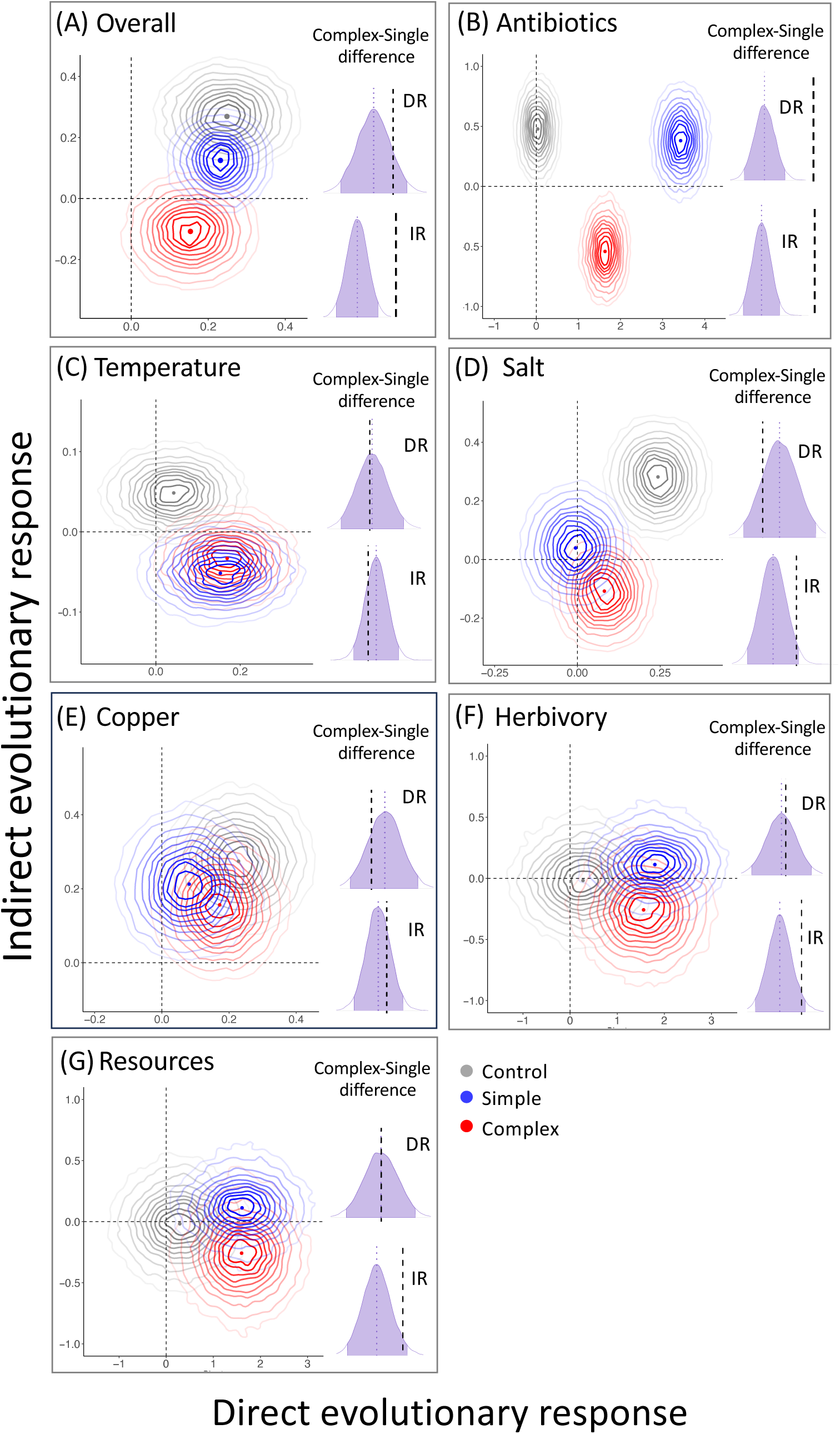
Joint posterior distributions of the direct and indirect responses to selection, for the control, simple, and complex stress evolutionary treatments. Closed lines are the probability isoclines, spanning from 10% to 95% from darker to lighter. The coloured dots are the median values of the posterior distributions. The overall distribution across stressors (11 species and 20 strains, reduced dataset) is shown in (A), and the marginal distributions for antibiotics (1 species and 4 strains) in (B), temperature (8 species and 12 strains) in (C), salt (9 species and 17 strains) in (D), copper (6 species and 8 strains) in (E), herbivory (1 species and 1 strain) in (F), and resources (1 species and 1 strain) in (G). The purple distributions are the contrasts between treatments, showing the difference between complex and single stressor for direct (DR) and indirect (IR) responses. Distributions with negative values that do not overlap with zero (black dashed line) indicate lower change in fitness in complex *vs.* single stressor.

While the signal for direct response to single or combined stressor depended on stressor identity, for indirect response there was a strong and largely consistent signal for complex environments leading to higher indirect costs of adaptation than simple environments (**Fig. 3** and **Table S4**), in line with the main findings. However, temperature, one of our most commonly used stressors, led to similar evolutionary change when applied as single *vs.* combined stressor, both for direct (**Fig. 3C**, LMM Simple: 0.152 [-0.003, 0.305]; Complex: 0.168 [0.031, 0.302]) and indirect responses (**Fig. 3C**; Simple: -0.051 [-0.102, -0.0001]; Complex: -0.033 [-0.082, 0.018]). Further detailed results for each stressor are reported in **Table S4** (see also **Fig. S11**).

### The relationship between initial fitness and fitness change was shallower in complex environments

We found that greater direct evolutionary responses (fitness gain in selective environment) occurred for populations that initially experienced more maladaptation (*i.e.*, started from lower fitness relative to the control environment), after controlling for potential spurious correlations due to the dependency of both variables on initial fitness by bootstrapping the data 10k times (see Material and Methods) (**Fig. 4**). This is consistent with a common observation in experimental evolution, described as the rule of declining adaptability (50, 51). Specifically, the negative relationship between initial maladaptation and evolutionary fitness gain was well-supported for simple environments (median and 95% CI slope posterior for simple: -0.218 [-0.490, -0.027]), but less steep and less well-supported for complex environments (Complex: -0.069 [-0.268, 0.062]), with clear evidence for a difference in slope between these treatments (posterior distribution in the inset of **Figure 4**). This indicates that more complex environments impose more limits on adaptation, even when controlling for the initial maladaptation. While the points with the largest maladaptation were associated with a small direct response, this did not affect the relationship between maladaptation and direct response (**Fig. S12**), and our data does not suggest a non-linear relationship overall (see **Table S5** with model comparison).

**Fig. 4.**
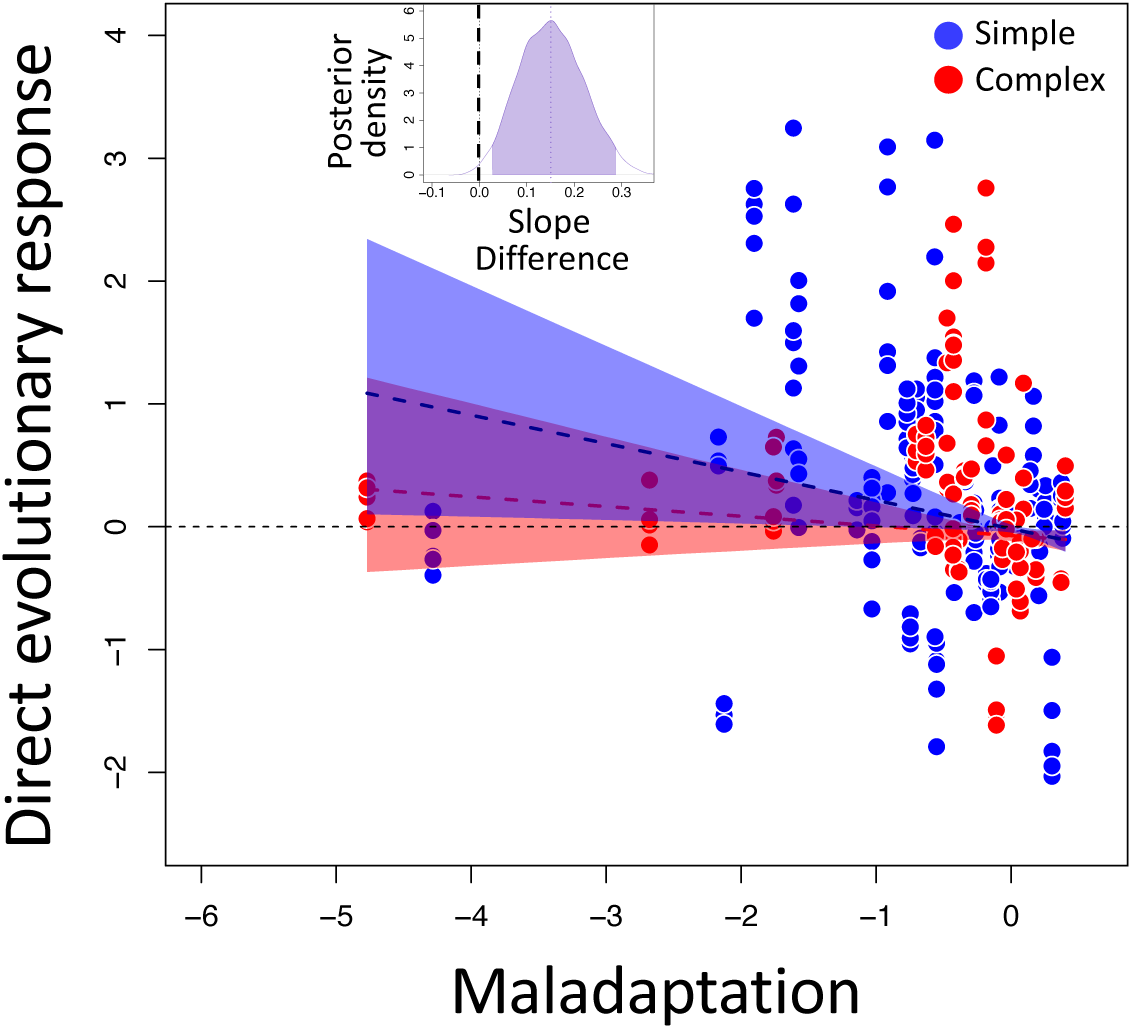
Declining adaptability in simple vs. complex environments. Relationship between direct response to selection and initial maladaptation (fitness in treatment minus fitness in control), for single (blue) and combined (red) stressors. Each point represents an independent evolutionary replicate, standardized as the difference to the average initial fitness of its corresponding control (14 species and 23 strains, further details in the Material and Methods). The data were bootstrapped 10k times and randomly assigning to the response and predictor variable. The dashed lines are the medians, and the shaded areas the 95% CI of the 10k model predictions. The inset represents the posterior distribution of the difference complex minus simple, for the slope of the relationship between direct response to selection and initial maladaptation.

## Discussion

Using a coordinated experiment combining a diversity of organisms and stressor types, we uncovered differences in patterns of adaptation in response to single *vs.* combined stressors, showing clear effects of environmental complexity on evolutionary adaptation. Earlier studies across different fields and biological systems have found that combinations of herbicides (23), antibiotic molecules (17, 22), or antibiotics and phages (52) (or predators (53)), led to lower evolved levels of resistance. By contrast, the green alga *Chlamydomonas reinhardtii* evolved for ∼450 generations in 96 treatments that included up to eight combinations of simultaneous stressors, showed higher fitness (growth rates) when evolved under combined stressors (37). However, this result was explained by the fact that more stressors led to larger initial fitness drop (stronger maladaptation) in this experiment. Here in contrast, we aimed at keeping maladaptation comparable when going from simple to combined stressors.

One of our most striking results was that populations that evolved under combined stressors consistently exhibited a loss in fitness when tested in control environmental conditions, while those evolved under single stressors showed a fitness increase. Testing for correlated responses in other environments than where selection took place should therefore be crucial to fully understand eco-evolutionary trajectories in response to multiple stressors. This approach is often done in studies on evolution under single stressors. For instance, the evolution of the marine diatom *Chaetoceros simplex* under nitrogen limitation precluded thermal adaptation, questioning the adaptive potential to higher temperature of these organisms under future scenarios of reduced nitrogen availability (54). Thermal selection of the marine diatom *Thalassiosira pseudonana* and four freshwater protist species showed correlated evolutionary responses in functional and life-history traits (55, 56). The fitness changes of *S. cerevisiae* populations selected in one environment, and assayed in another, tended to follow consistent trajectories (57). For instance, populations evolved in a rich, non-selective growth medium (YPD) *plus* acetic acid, tend to lose fitness when tested back in YPD alone, resembling our findings for indirect responses. Extending this effort to complex stressors should prove especially useful (see also (58)), and could have direct implications for the management of resistance in pests and pathogens exposed to combinations of pesticide and antibiotics (59).

Despite the strong signal for indirect response and the general trend for direct responses under combined stressors, there was also substantial variation among species and strains in their responses to evolutionary treatments, as indicated by the variance partitioning and the per-species/strain analyses. Such heterogeneity of responses is not surprising considering the diverse life-histories and ecological niches of species in this study, as well as the variable stressor combinations, highlighting the usefulness of coordinated experiments that combine a broad diversity of taxa. This also reveals a fundamental limit of combined-stressor studies: a single genetic background may be insufficient to obtain reliable and general results within species (intraspecific), and similarly for species within broader taxonomic groups. Thus, to understand both short- or long-term responses to complex environments, it is crucial to consider both intra and interspecific variation, as also suggested by recent findings on the evolution of thermal performance curve (single stressor) in different species and strains of yeast (60).

The variability of responses among stressors likely reflects their different modes of action and breadth of impact on the organism’s physiology. Temperature is of particular interest, as heat-waves and rising mean temperatures are expected to affect all ecosystems worldwide (61), often in combination with other stressors (*e.g.*, pollutants, pH, salinity). In our study, evolutionary responses to temperature were very similar when applied alone or combined with another stressor, and for direct *vs.* indirect responses (**Fig. 3A**), suggesting that the mechanisms for facing thermal stress are consistent and overrule the influence of other stressors. These findings are in line with genomic analyses of the copepod *Acartia tonsa* evolved under warming (simple), acidification (simple), or concurrent warming and acidification stress (complex) (62). That study found that 37% of responses to selection were shared between warming and complex stress, but only 1% between the latter and acidification (62), thereby suggesting that warming was the dominant selective force. Regarding the involved mechanisms, another study showed that heat-adapted lines of a marine diatom (*Chaetoceros simplex*) prioritized protein protection over synthesizing new proteins or photosynthetic structures (63). Although this helped them survive high-temperatures, it also reduced their growth and limited adaptation when the levels of nitrate were low, suggesting that thermal adaptation might be constrained in nutrient-poor conditions (see also (64)). Another important combination of stressors for which some short-term response mechanisms are known is salt and copper, as they share the same Na⁺-transport machinery, which copper can also disrupt (65). However, the resulting toxicity is non-monotonic across the salinity range, and a high level of salinity (similar to seawater level) can be partially protective against copper (65). In light of their different modes of action, single stressors might themselves be considered to have different degrees of complexity, in terms of the biological challenges that they impose on organisms, and the evolutionary responses that they elicit. For instance, temperature can affect many biological processes (*e.g.*, protein stability, enzymatic activity, membrane permeability, metabolic rate (66, 67)); antibiotics may only target a few specific mechanisms (*e.g.*, chloramphenicol in this study is an inhibitor of mitochondrial protein synthesis (68)); and salt can induce both osmotic and oxidative stress (69, 70). Nevertheless, this should not strongly impact our results, as the random structure of our statistical analyses was such that treatment comparisons were made within system, and we expect combined stressors to represent a more complex environment than single stressors, regardless of how complex each of these single stressors is.

The negative relationship that we found between fitness gain and initial fitness is consistent with the “rule of declining adaptability” described in evolutionary theory (71, 72) and a number of experimental studies (50, 51, 57). This broad pattern is generally explained by diminishing-returns epistasis, wherein low-fitness genotypes have access to mutations with larger fitness gains than high-fitness genotypes (73, 74). This is also described as “global” epistasis, to emphasize that fitness effects depend on a macroscopic aspect of the organism (its fitness). For instance, recent work with *S. cerevisiae* found that the fitness effects of ∼100 mutations exhibited a consistent negative relationship with the fitness of their background genotype, with minimal variation in slope across the 6 environmental conditions tested (75). This pattern is notably predicted to arise in Fisher’s geometrical model of adaptation (71, 76), from which the cost of complexity was originally defined (18, 77). The shallower relationship that we observed under combined stressors is consistent with a cost of complexity (*sensu* Fisher-Orr (18, 77)), provided that complex environments exert selection on more traits. This hypothesis could be investigated directly by testing whether the loci under selection (or even better, the specific mutations that rose in frequency at these loci) were more pleiotropic in complex than in simple environments, for instance they affected more GO terms, connections in regulatory networks, or tissues for multicellular organisms (78). As the cost of complexity has been previously defined with respect to the complexity of the organism rather than of the environment, we might have expected metrics of organismal complexity to also influence the rate of adaptation, or how it changes with the complexity of the environment. However, we found no signal for species with more protein-coding genes or protein families (proxies for the number of functions/traits, and therefore for organismal complexity (79)) to be less constrained in their evolutionary response to combined stressors (Supplementary Text, **Fig. S13A-B**). Beyond the cost of complexity, another non-exclusive possibility for the difference between evolutionary responses to single *vs.* combined stressors could be that they lead to different fitness maxima. Obtaining multiple fitness measurements along the experiments could have revealed to which extent different treatments had reached different plateaus of fitness, but this was not feasible across systems.

The impacts of stress complexity on adaptation that we detected should be mediated mostly by probabilities of occurrence and fixation of beneficial mutations in systems initiated by isogenic lines (which form the majority of our dataset), and by the strength and efficiency of selection in the presence of linkage in systems initiated with standing variation (two plant species and one experimental group of the chlorophyte *D. salina*). While a formal comparison of evolutionary responses from *de novo* mutations *vs.* standing variation is beyond reach in our study (as only 3 out of 23 strains relied on standing genetic variation), separate analyses on these 3 strains revealed a weak signal for the direct response to selection (**Fig. S14**) that seemed to conform to the “*de novo*” pattern (**Fig. S5A-B**). Hence, we have no indication that the responses to single *vs.* combined stressors was mostly explained by differences in origination and fixation of new mutations in our experiment.

### Perspectives and conclusion

Understanding and predicting the ecological and evolutionary consequences of combined stressors is a pressing challenge, relevant to applied fields (80–83). Our work reveals that transversal patterns exist to some extent within and among species, suggesting general rules about the evolutionary consequences of environmental complexity. However, intra- and interspecific variability in the effects of single and combined stressors also highlight the need to further integrate the different levels of biological diversity into experimental studies and predictive attempts. While future extensions could include more stressors, fluctuating environments, and a broader taxonomic sampling, our large-scale coordinated experiment does highlight the critical need for the combined-stressor research to consider evolutionary changes.

## Materials and Methods

### Experimental treatments and data collection

The evolutionary experiments followed the same rationale and analogous protocols across species, and had four treatments: a control, two single stressors applied separately (simple stress), and the two stressors combined (complex stress). For three strains (**Table 1**), three single stressors and three combined stressors (all pairs of the single stressors) were used. For the combined-stressor treatments, the levels of each individual stressor were adjusted so that their combined impact on fitness was comparable to that of the single stressors. Each organism was then propagated for several generations in all treatments (3 to 12 independent evolutionary replicates per strain per treatment, precise description in the Supporting Text and **Table 1**), and at the end we measured the fitness of all 14 species in their own evolutionary treatment, and for 11 species also in the control conditions.

### Data inspection and statistical analyses

We analysed the data using Bayesian Linear Mixed Models in R (version 4.2.0) with the “rstan” (v. 2.32.7; Stan Development Team, 2025 (84)), “brms” (v. 2.22, (85)), and “rethinking” (v. 2.21; (49)) packages.

For unicellular organisms, we considered their exponential growth rate as fitness proxy (86). Population density was log-transformed and the time series for each growth curve was visually inspected to identify the exponential growth phase (shading, **Fig. 1**), and remove potential outliers and no-growth cases. Methods that automatically screen growth curves for the steepest slope (exponential growth) or fit a parametric model such as the logistic function (*e.g.*, growthcurver (87), rstan model fitting (88), LoLinR (89)) were also tested, but discarded as more prone to spurious results (such as estimating locally high slopes that do not correspond to the true exponential phase). Both log-density and time were then standardised by centering and reducing across treatments, within each strain of each species. This enabled comparison of the influence of treatments among species with different growth rates and generation times (see Supporting Text for plants).

We fitted Bayesian linear models to the identified growth phase, and estimated the exponential growth rate (*r*_0_) as the slope of log abundance over time (**Table S6,** and **S7**). Models were fitted to the most relevant biological level, namely the experimental evolution line. When the data included technical replicates, these were treated as a random effect. We used vaguely informative priors for the intercepts and slope parameters (normal and half-normal distribution, respectively), with mean 0 and standard deviation 1. The slope posteriors provided standardized estimates of the intrinsic growth rates *r*_0_ (**Fig. S2**, and **S7**). In the follow-up analyses, we included the associated error (standard deviation) of the slope posteriors to effectively weight fitness effect sizes by study precision, which lead to conservative results (49). Convergence (R-hat=1), the absence of divergent transitions, and effective sample size was confirmed for these and the models described below, using 3 independent chains. Additional details on chain length and warmup phase are provided in the SI Tables. Prior and posterior checks were performed for each analysis as illustrated in **Fig. S15-16**.

We conducted five analyses on these fitness estimates, always including “Species” and “Strain” as random effects, but not phylogeny. Following Cinar *et al.* (90), under weak phylogenetic signal as in our study (mean phylogenetic matrix correlation of 0.27), the phylogenetic and non-phylogenetic species variability would collapse into one, reducing the model reliability (91). Complementary analyses incorporating the phylogeny led to an almost unidentifiable species random term variance (Strain = 11.4%, Species = 0.001%, phylo = 79.4%) compared to the simpler model (Strain = 20.4%, Species = 64.3%). This is because the redundancy between the two terms results in an unstable variance split (87). Further, the inclusion of the phylogeny did not significantly improve the model fit (**Table S8**), nor did it change the main effects. We therefore retained the simpler model with strain and species as independent random effects, as the inclusion of the phylogeny would in fact call into question the estimates’ trustworthiness (90).

The stressor identity was also not included as a random effect because some stressors were used in one species only (**Table 1**), leading to divergent transitions. We evaluated marginal and conditional R^2^, and the heterogeneity (I^2^) of direct and indirect responses using posteriors of the variances (σ^2^) (91–93). The posteriors were extracted from the models, and we calculated I^2^ with 95% compatibility intervals for the random effects, following Viechtbauer (94).

For the main analyses, we first investigated the direct responses to selection on the complete dataset (14 species, 23 strains) using a linear multilevel model. We asked whether the change in fitness between the initial and final time point, assayed under the same conditions as those under which experimental evolution took place, was different between the control, single, and combined stressors (3 treatments x 2 time points).

Second, we analysed indirect responses, defined as the change in fitness under experimental evolution (with stressors) assayed under control environmental conditions (without stressors). For each evolved population, we compared its fitness in control conditions at the end of the experiment against the initial fitness in control conditions (4 explanatory factors). For the three species *B. rapa*, *O. tauri*, and *P. costavermella* the indirect response assays were not logistically feasible. Thus, this analysis was conducted on a reduced dataset (11 species, 20 strains, Table 1), using the same model structure as for direct responses.

Third, we merged direct and indirect response data, focusing on the reduced dataset where both are available. After centering and reducing, we refitted Bayesian linear models to have all data on the same scale. The fitness estimates for the control treatment were the same for direct and indirect response, apart from *A. thaliana*, *D. salina*, and *V. aestuarianus* because their control treatments varied between responses due to specific protocol handling. We investigated direct and indirect response using the posteriors obtained from the single- and combined-stressor treatments only (change in fitness) and their contrasts.

Fourth, in post-hoc investigations we repeated the analysis above with respect to each to specific stressors (**Table 1**): temperature (8 species and 12 strains), salt (9 and 17), antibiotics (1 and 4), herbivory (1 and 1), resources (1 and 1), and copper (6 and 8). A complete model overview is provided in **Table S9**.

Lastly, using a linear multilevel model we explored whether the final change in fitness (direct response) was related to the initial cost of stress, and whether the slope of this relation depends on the stressor treatment. For the initial cost of stress, we averaged the initial fitness by strain (separately within single and combined stressor treatments), from which we subtracted the average initial fitness of the control populations. For the evolutionary change in fitness, we took the difference between the final and initial fitness estimate of each evolutionary replicate. The associated error was always propagated at each step, and the weighting procedure was applied to both responses and explanatory variables. We refitted the model 10k times after bootstrapping the data (initial fitness) and randomly assigning them to the response and predictor, thus making the axes fully independent, to avoid potential spurious correlations that may arise through shared error terms between the x- and y-axis (since both depend on initial fitness).

We additionally tested direct, indirect response and their relationship for each strain running the statistical analyses described above, without random effects (**Fig. S3, S4, S8, S9, and S11**).

## Acknowledgments

We thank Sylvain Gandon, Marine Combe, and members of the ExpEvolOcc network for discussion at the early stages of this project. We thank David Alvarez Ponce for providing the data on the protein family. We thank the editor and two anonymous reviewers for their insightful suggestions. This work was funded primarily by Occitanie Regional Council’s program “Key challenge BiodivOc”, grant ComplexAdapt (All participants), and received additional funding by Agence Nationale de la Recherche (ANR) project PHYTOMICS, ANR-21-CE02-0026 (GP), Agence Nationale de la Recherche, project POLLUCLIM, ANR-19-CE02-0021-01 (DL), Agence Nationale de la Recherche, project AraBreed, ANR-17-CE02-0018-01, (CV), Agence Nationale de la Recherche, TULIP, Laboratory of Excellence, ANR-10-LABX-41 (EA, CB, DC, GMC, GC, FE, AG, DL, HP, SJ, PR, LS).

## Data availability

All data and code used in the analysis will be provide on public repository such as Zenodo.

## Supporting Information Text

### Individual species protocol

#### Arabidopsis thaliana

##### Authors

François Vasseur, M. Stefania Przybylska, Cyrille Violle

##### Study organism

*Arabidopsis thaliana* is a small plant from the mustard family (Brassicaceae) widely distributed throughout Europe, Asia, and North America (1). At maturity, *A. thaliana* can reach 10 to 50 cm in height, and can produce a variable number of seeds, from a few dozen up to more than 5000 seeds. Due to its short life cycle (∼ 6 weeks), ease of manipulation, and well-characterised genome, *A. thaliana* is an established model species in several research fields from genetics and molecular investigations, biochemistry, physiology, to ecology and evolution (2, 3). *A. thaliana* can rapidly respond to selection (4), and its evolutionary history, as well as its response to environmental variables, are well studied (5).

##### Experimental setup and treatments

Before the beginning of the evolution experiment, we created 350 recombinant lines (F2 lines) from random crosses between 358 wild accessions spanning a large range of genetic, geographic, and phenotypic variation (6). Around 2500 seeds per cross were pooled to be used

In November 2017, we set up an experimental evolution project (AraBreed project) in the experimental field of Centre d’Ecologie Fonctionnelle et Evolutive (CEFE), Montpellier, France (43° 38’ 18“ N, 3° 51’ 44” E). 28 closed mesocosms with 1.2 m length x 1.2 m width x 1 m height were built in seven rows. Mesocosms were 1.5 m apart and contained entirely, both belowground and aboveground, in an insect-proof veil. During their installation, we removed the top 30 cm of soil (calcareous clay soil), steam sterilized it to prevent seed bank sprouting, mixed it with blond peat, and placed it back within the mesocosms. Mesocosms were assigned to four evolutionary treatments: resources single stressor (low Resources/ low Herbivory; R-/H-), herbivory single stressor (R+/H+), combined resources and herbivory stressors (R-/H+), and a control without stressors (R+/H-). Treatment assignment was random but ensuring that each row of mesocosms contained one replicate of all four treatments (4 treatments × 7 replicates). At the end of November 2017, we placed approximately 17500 randomly selected F2 seeds in each mesocosm in a density of around 1.2 seeds/cm^2^.

##### Rearing protocol

From 2018 to 2020, we let plant generations spontaneously grow in the mesocosms, with spontaneous seed establishment and germination. In this period, from January to June (*i.e.*, the season at which *A. thaliana* generally grows and sets seeds), we applied the treatments to the mesocosms and we harvested fruits for seed extraction. We watered and fertilized mesocosms assigned to high resources (R+) with commercial organic fertilizer twice per week. Low resource (R-) mesocosms received no applications of water or fertilizer. In the same period, we introduced intermittently snails (*Helix aspersa*, a generalist herbivore) into the mesocosms assigned to high herbivory (H+). Meanwhile, we sprayed mesocosms assigned to low herbivory (H-) with large-spectrum insecticide every three weeks. Regarding fruit harvest, every three weeks, we harvested one fruit per individual located in different parts of the mesocosm, for a maximum of 100 fruits per mesocosm. We pooled fruits of a given mesocosm in a given year and extracted seeds, which were then stored in air dry condition at 4°C.

##### Population size as direct responses to selective environment

Using pictures of each mesocosm taken in March 2018-2022, the month that roughly corresponded to the peak of vegetative growth, we estimated population size in all 28 mesocosms. For that, we counted the number of plants in the images using the cell counter plug-in in ImageJ. To obtain the fitness estimates for each mesocosm, we fitted Bayesian linear models to the log-transformed population size data. We used vaguely informative prior (normal distribution), with mean 0 and standard deviation 1. Since populations evolved with natural recruitment, comparing the size of the first generation (G1) to the number of seeds sown initially (G0) was not relevant to estimate initial fitness. We thus calculated the difference in fitness between G2 and G1, and G5 to G4, and considered these values as initial and final fitness, respectively. These fitness estimates were used in the main analyses (direct responses) together with their corresponding standard deviation, which was included as measurement error in the follow-up statistics to effectively weight fitness effect sizes by precision. The description of the main analyses with references of the statistical software and packages are provided in the main text.

##### Common garden and indirect responses in non-selective environment

We performed a completely randomized common garden experiment between February and July 2021 under shade cloth in the experimental field of Centre d’Ecologie Fonctionnelle et Evolutive (CEFE), Montpellier, France (7). Pots of 0.08 L were filled with a sterilized soil mixture composed of 50 % river sand, 37.5 % calcareous clay soil from the experimental field at CEFE, and 12.5 % blond peat moss. Such conditions represented a non-selective environment for the evolutionary lines, further allowing control for the confounding effect of phenotypic plasticity (8). As detailed below, the data obtained during this common garden were used as indirect fitness responses in the main analyses. 630 individuals were planted in separate pots organized in three blocks: 198 individuals from the same pool of F2 seeds that were used to found populations in the mesocosms (G0), and 432 individuals from the third generation of evolution (G3) in the four treatments (108 individuals × 4 treatments). At the end of the reproduction phase (fruit ripening), inflorescences were cut and photographed. On inflorescence pictures, all fruits were counted and the length of three fruits per plant were measured using ImageJ. To obtain fitness estimates, we then considered the number of fruits produced per plant weighted by their length (number of fruits * mean fruit length). We fitted Bayesian linear models to the log-transformed fruits’ metric using vaguely informative prior (normal distribution), with mean zero and standard deviation one. Estimates for G0 and G3, corresponding to the initial and final fitness, were used in the main analyses (indirect responses) together with their standard deviation (measurement error, as described in the population size paragraph above). The main analyses, and references on statistical software and package, are detailed in the main text.

#### Brassica rapa

##### Authors

Jeanne Tonnabel, Denis Orcel, Marie Challe, François Vasseur, Cyrille Violle

##### Study organism

Our experimental evolution lines were established using the *Brassica rapa* Wisconsin Fast Plant (WFP) line (Carolina Biological Supply Company, Burlington, North Carolina, USA). This line is particularly well suited for experimental evolution studies, as it was derived from a broad sampling of natural *B. rapa* populations that subsequently underwent artificial selection for a short generation time (∼8 weeks), while retaining considerable genetic and phenotypic variation (9, 10). *B. rapa* is a hermaphroditic, insect-pollinated annual herb in the Brassicaceae family. Like other members of the genus, it possesses a genetic self-incompatibility system (11, 12), although this mechanism is leaky, allowing variable levels of self-fertilization. Such incomplete self-incompatibility has been observed in the WFP line as well, where selfing has evolved under experimental evolution conditions (13). Pollination in our experiment was performed by the fly *Phormia terraenovae*, which did not evolve during the course of the study. This species had been validated as an effective pollinator of *B. rapa* in a preliminary experiment, and was reared from larvae obtained from a commercial supplier of fishing bait (https://appats-michel.fr/).

##### Experimental setup, treatments, and growing protocol

To initiate the experimental evolution populations, plants first underwent three to four generations of selection under strong polygamous mating in five replicate populations. In each generation, seeds were collected from 85 parental plants, each of which had three flowers pollinated with mixed pollen derived from 12 anthers per plant pooled across all 85 individuals. Seeds from the three pollinated flowers of each plant were combined to produce the next generation. The present study used seeds from the third and fourth generations of these lines, hereafter referred to as the “source population”.

Our protocol followed standard experimental evolution procedures, in which natural selection acts on standing genetic variation present in the source population (14). We subjected this initial standing genetic variation to four treatments: light deprivation (single stressor), nutrient deprivation (single stressor), the combination of both stressors (combined stressor) and a control (no stressor). Experimental populations were grown in a greenhouse arranged in a 4 × 5 grid. Because of a potential light gradient in the greenhouse, population positions were randomized following specific rules: (i) each row (corresponding to a given distance from the external glass wall) contained one population from each treatment; (ii) treatment assignments were rotated across positions at each generation; and (ii) each population was relocated to a different position between generations.

Using well-mixed seeds from the source population, we established 20 experimental populations, randomly assigning five populations to each of the four treatments in the first generation. All experimental populations were cultivated in a single greenhouse at the CNRS Experimental Plateform in Montpellier. Each population was grown in an individual 80 x 80cm tray containing 100 plants arranged in a 10x10 grid, with individuals spaced 7 cm apart. Two seeds were sown per position, to allow estimation of germination rate based on the absence of plants at a given position, which was later incorporated into individual fitness estimates. After two weeks, when both seeds germinated at a given position and none at other positions, one seedling was transplanted elsewhere within the same population to maximize the total number of plants while retaining a single seedling per position. Germination rate was calculated as the total number of seedlings divided by the total number of seeds sown.

All populations were grown in sterilized, nutrient-poor soil composed of 2/3 sand and 1/3 compost. During the first three weeks of growth, liquid nutrients were supplied at watering to temporarily restore nutrient-rich conditions and allow plants to establish before flowering. We used a liquid fertilizer containing nitrogen, phosphorus and potassium (Universal Liquid Fertilizer, StarJardin). Each population received weekly 10L watering with a concentration of 5mL/L of liquid fertilizer. After this initial period, nutrient availability was manipulated by varying the concentration of liquid fertilizer: 5mL/L in the control and light-deprivation treatments to maintain nutrient-rich conditions, and 2.5mL/L in the double-stress treatment.

At flowering onset, each population tray was enclosed within a mesh cage. A paper cup containing approximately 100 *P. terraenovae* larvae and a glass of sugary water covered with fine mesh (allowing access to the solution while preventing drowning) were placed inside each cage. The high abundance of pollinators ensured that none of the populations experienced pollen limitation. Because pollinators were confined to their respective populations, reproduction occurred independently in each population, preventing any gene flow among them. Pollinator densities were visually monitored and adjusted as needed by adding larvae to maintain similar numbers across populations until all plants had senesced.

##### Seed mass as direct responses to selective environment

At the end of each generation, when plants had died, we harvested every individual, counted its total number of fruits, and stored the fruits separately by plant. Seeds were then manually extracted, and total seed mass was measured using a precision balance (Sartorius, Entris II 220g/0.1mg).

We measured the total seed mass for all plants that germinated in each experimental population. To obtain fitness estimates at the population level (20 independent evolutionary replicates), we fitted Bayesian linear models to the log-transformed seed mass using vaguely informative prior (normal distribution), with mean 0 and standard deviation 1. The individual plant identity, for which the seed mass was calculated, was included as random term. We obtained estimates of averaged seed mass for first and third generation, corresponding to the initial and final fitness, were used in the main analyses (direct responses) together with their standard deviation (error used in the follow-up statistics to fitness effect sizes by precision). The main analyses, and references on statistical software and package, are detailed in the main text. To verify that total seed mass provided a reliable proxy for seed number, we randomly chose 30 plants per population and counted all produced seeds - or a maximum of 30 fruits for highly fecund individuals - before weighing them at the end of the first generation. A linear regression of total seed number on total seed mass confirmed that seed mass is a reliable predictor of seed production (linear model estimate = 601.4 ± 15.7 SD; t = 38.4, p < 0.0001; R² = 0.73).

Seeds from all plants within a population were bulked to establish the next generation, ensuring that the contribution of each individual to the next generation reflected its reproductive success. Each experimental population thus evolved independently under its assigned treatment. In total, three generations of evolution were completed for the 20 experimental populations, corresponding to approximately 6000 plants in total. The fitness assessments took place between October 2023 and January 2024, February and May 2024, and October 2024 and January 2025, respectively.

#### Colpidium striatum

Lynn Govaert, Laurence Bolick, Fabian Schneider-Nettsträter and Marielle Kathrin Wenzel

##### Study organism

Freshwater ciliates are well suited organisms for addressing ecological and evolutionary questions, they have high population densities, short generation times, are easy to maintain, and the microcosms they live in are easy to manipulat (15). *Colpidium striatum* is commonly found in freshwater and frequently used in microcosm experiments (15). An original culture of *Colpidium striatum* was purchased at Carolina Biocompany and maintained for several years in the laboratory kept in 250-mL autoclaved erlenmeyers containing two wheat seeds and filled with 100 mL of autoclaved protozoan medium maintained under standard laboratory conditions (20°C and 24h light condition). Since November 2021, protozoan medium consisted of Volvic water Naturelle with 0.40 g/L alfalfa powder (A-4363-BIO Piowald.com) and fed with the bacteria *Serratia marcescens*. From this culture, a single cell was taken and washed to start a monoclonal culture. Throughout the experiment, *Colpidium striatum* reproduces asexually.

##### Experimental setup and treatments

At the end of June 2023, an experimental evolution experiment was started using a clonal culture of *C. striatum*. Initially, a single cell was isolated from a genetically diverse stock culture maintained under standard laboratory conditions (*i.e.,* 20°C and 24h light condition) and grown to a dense population. Then, triplicates of this population were exposed to an increased salinity, copper concentration and temperature as well as all pairwise combinations of these stressors. Specifically, triplicates were kept as a control (0 g/L NaCl, 0 µg/L CuSO_4_, 20°C), exposed to increased temperature (0 g/L NaCl, 0 µg/L CuSO_4_, 24°C), increased salinity (3 g/L NaCl, 0 µg/L CuSO4, 20°C), increased copper concentration (0 g/L NaCl, 200 μg/L CuSO4), or exposed to an elevated level of each two-stressor combination (**Table S10**). In total, these were 21 cultures that were maintained for one year. The experimental evolution populations were transferred into new medium at a 3-week interval using autoclaved 250-mL Schott bottles with two wheat seeds, which were maintained in a 24h-light environment and were fed with the bacteria *Serratia marcescens*.

The concentrations of salt, copper and temperature were chosen based on previous pilot experiments where we observed extinction of *C. striatum* when exposed to 4 g/L NaCl (when not prior acclimated) and showed a large fitness reduction at 26°C (*95*). In addition, the combination of 3 g/L NaCl and 26°C previously indicated a high risk for population extinction (16). To track the population growth of the ciliates, the cell density was measured by taking videos. For this, the bottles were sampled (250 μL) daily for 8 days, then every 2nd day for another five timepoints. The videos were taken with an Olympus SZX16 stereo-stereomicroscope (1.6-fold magnification) and Hamamatsu Orca Flash 4 video camera. 10 second videos were recorded of an effective sample volume 36.125 μL (height 0.5 mm). The videos were processed with the software ImageJ (version 1.46a) and an adapted version of the BEMOVI R package (17) to extract information on density.

##### Rearing and transferring protocol

Every three weeks, 20% (*i.e.*, 20 mL) of the experimental evolution populations were transferred into new medium using autoclaved 250-mL Schott bottles with two wheat seeds and filled with 80 mL autoclaved protozoan medium (i.e., autoclaved Volvic water and 0.4 g/L alfalfa powder) and were placed in incubators at the respective treatment temperatures with full 24h-light environment. Cultures were fed with the bacteria *Serratia marcescens*. Given that *C. striatum* has an average generation time of 16h, this resulted on average in 32 generations between transfers.

##### Final phenotypic tests and data collection

Mid-July 2024, a common garden experiment was started to test for evolutionary changes in the evolved *Colpidium* populations from the different stressor environments. Specifically, each of the populations evolved in a stressor environment was measured in the control environment (*i.e.*, 0 g/L NaCl, 0 µg/L CuSO_4_ and 20°C) and their original stressor environment. Additionally, the populations evolved in the control environment were measured in all the stressor environments and in the control environment (**Table S11**).

For the common garden experiment, we used evolved populations of the following stressor combinations (**Table S11**): three replicates of the control, two replicates of the high salt environment (single salinity), three replicates of the high copper environment (single copper), three replicates of the high temperature environment (single temperature), three replicates of the combined high salt and high copper environment (combined salt x copper), and two replicates of the high temperature and high copper environment (combined temperature x copper). Unfortunately, all populations from the high temperature and high salinity environment became extinct by the end of the experimental evolution phase, thus the combined temperature and salinity was not available. In total, we had 16 uniquely evolved populations.

From each of these populations 10 individuals were washed by passing each individual with minimum of liquid through 5 drops of autoclaved protozoan medium to reduce bacterial load coming from the selection cultures. The washing of the individual cells was done using the protozoan medium of their respective stressor environment. Each of the 16 evolved populations was triplicated and exposed to their respective stressor environment and to the control (as described earlier), resulting in a total of 132 experimental units. Experimental units consisted of autoclaved 250-mL Schott bottles with two wheat seeds and were filled with 3 mL of the corresponding selection culture (cultured for 1.5 weeks in their respective stressor treatment) and with 97 mL of the corresponding protozoan medium with salinity and copper concentrations modified such that adding the selection culture resulted in salinity and copper concentrations of 3 g/L NaCl and 200 µg/L CuSO_4_, respectively (**Table S12**). Because transferring the selection cultures into their common garden environments brings additional medium with it, the control condition was set to a concentration of 0.093 g/L NaCl and 6.186 µg/L CuSO_4_, which is known to not have any effects on the ciliate densities from prior experience. For temperature treatments, autoclaved protozoan medium of the respective salinity and copper concentrations was placed in a 24°C incubator to obtain the correct temperature. All autoclaved protozoan medium was enriched with a 5% 36-h bacterial culture of *Serratia marcescens* reflecting an addition of 24,338,256 CFU/mL. Experimental units were given a random ID and placed in either 20 or 24°C incubators depending on the temperature treatment and maintained in a full 24h light environment. To track the population growth of the ciliate populations during the common garden experiment, the cell density was measured by taking videos. For this, we took 150 μL daily for 9 days, then 2 days no sample followed and then by every 2 days for three timepoints. The last samples were taken another three days later (in total three weeks). The videos were taken and processed as previously described.

#### Dunaliella salina (all strains)

##### Authors

Philipp Hummer, Stanislas Fereol and Luis-Miguel Chevin

##### Study organism

*Dunaliella salina* is a halophile unicellular green alga that is well-studied because of to its extreme physiology. It is one of the most salt-tolerant eukaryotes, and is the main primary producer in hypersaline environments (18, 19). It is commonly found in salt lakes, evaporation ponds of salterns, hypersaline coastal waters and salt marshes worldwide (20). *D. salina* can also withstand wide ranges of environmental stresses such as temperature (from -35°C to +40°C) and nitrogen content (*99, 100*). In standardised laboratory conditions at 24°C and 12/12 dark/light cycle (200 μmol m^-2^ s^-1^ light intensity), it reproduces asexually with a growth rate of ∼ 1 generation per day (21).

##### Experimental setup and treatments

To start our experiment, we used 4 strains isolated from salterns in southern France (clonal/isogenic strains; for details see **Table S13**) and their combination (mixed strain; standing genetic variation). The mixed strain was established by mixing equal densities of the four different strains. We call these 5 genetic backgrounds “strains” (including the mixed strain) for simplicity.

*D. salina* strains were cultivated in flasks, using artificial seawater complemented with NaCl and nutritive F/2 medium. The required salinity was obtained by mixing hypo-(0M) and hyper-saline (4.8M) solutions (**Table S14**). Before being introduced in the evolutionary treatments, the strains were maintained under favourable laboratory conditions of 2M salinity and 24°C. The carrying capacity of each strain was measured (approximately 470k cells/mL for S1, 940k for S3, 830k for S11 and 770k for S13) and adjusted to start populations at similar densities (approximately 100k cells/mL). The measurements of cell concentration during all experimental phases were obtained with the Guava easyCyte HT flow cytometer and the corresponding software guavasoft 3.4 by Luminex. An established pipeline (22) allowed us to discriminate between live and dead *D. salina* cells, as well as other particles on the basis of forward scatter (FSC), side scatter (SSC) and Red and yellow Fluorescence.

For each strain, we established 4 evolutionary treatments in 3 replicates (**Table S15**) by manipulating temperature and salinity. In the control non-stressful treatment, we used 24°C and 2M salinity, in line with previous work in the group (*e.g.*, (22)). For the 3 stressful treatments (salinity stress, temperature stress, and the combination of the two), pilot experiments indicated that a temperature of 36°C and salinities of 3.5M or 4M NaCl would cause fitness reduction with little risk of population extinctions.

For each strain, the experimental flasks were set up at different salt concentrations by mixing 2mL of the strain with 13mL of medium. The flasks were then placed into two incubators at their respective treatment temperatures under 12/12 dark/light cycle (200 μmol m^-2^ s^-1^ light intensity). Note that the 3.5M salt concentration only occurred at 36°C (Table S4.3) where our preliminary tests suggested that 4M would be too stressful. This gave us a total of 75 evolutionary replicates: 5 strains x 2 salinities (2M, 4M) x 3 replicates at 24° C, and 5 strains x 3 salinities (2M, 3.5M, 4M) x 3 replicates at 36°C. In addition, five media flasks with no cells (for the different conditions) were added to calibrate spectrometer measurements. Air composition was not controlled in the incubator, but carbon dioxide necessary to photosynthesis was provided in dissolved form. At 36°C, we additionally placed a tray filled with water to prevent excessive evaporation. At the first transfer, at the start of the experimental evolution phase, we tracked population growth with daily measurements of population density over eleven days for all replicates of the methods described above.

##### Rearing and transferring protocol

Each of the 75 evolutionary lines was transferred into new flasks weekly. The standard transfer (for strains S1, S13 and Mix) consisted of diluting 2mL of culture into 13mL of fresh medium at the same salinity (i.e., a dilution rate of 2/15). However, dilution rates had to be adjusted for some strains and transfers to prevent out-dilution, as detailed in **Table S16**. Before each transfer, cell concentration was measured using a flow cytometer (see below) to keep track of densities and reduce the risk of out-diluting the cultures. Nonetheless, all replicates from strain S11 failed to grow and were lost at transfer 5 (**Table S16**).

##### Final phenotypic tests and data collection

At the last transfer, we tracked population growth with daily measurements of population density over eleven days. Each of the populations evolved in a stressor environment was measured in their own stressor environment and in the control environment. All measurements of cell concentration for every evolutionary line (each strain and replicate) were obtained with the Guava easyCyte HT flow cytometer and the corresponding software guavasoft 3.4 by Luminex.

### Euglena gracilis, and Tetrahymena thermophila (strain T1)

#### Authors

João Gabriel Colares Silveira, Justina Givens, Claire Gougat-Barbera, Marie-Ange Devillez, Giacomo Zilio, Emanuel A. Fronhofer

##### Study organism

*Tetrahymena thermophila* and *Euglena gracilis* were chosen due to being two low-maintenance unicellular eukaryote species with very distinct life-history traits and feeding habits. *T. thermophila* and *E. gracilis* have respectively generation times of 2-4 hours (23–25) and 10-30 hours (26, 27), and present heterotrophs (28) and mixotrophy (29) life style. *T. thermophila* and *E. gracilis* have been extensively used as model unicellular eukaryotes in a variety of scientific works (15).

##### Experimental setup and treatments

This study was designed around a total of 72 clonal stock cultures or “ancestor populations”, of *T. thermophila* and 24 of *E. gracilis* lines. Populations of the two species were subjected to eight treatment conditions, each replicated six times. These treatments involved the presence of one or more environmental variables: moderately elevated temperature and the presence of copper (CuSO_4_) and/or a salinity (NaCl) concentration. All treatments were also conducted under ambient temperatures with the presence of a control.

The specific concentrations of CuSO_4_ and NaCl, as well as the temperature regimes for the evolutionary phase, were chosen through preliminary experiments with the goal of making initial fitness effects homogenous (*e.g.,* same fitness loss).

##### Rearing and transferring protocol

Each line was kept in batch cultures fed with filtered bacterized lettuce medium (BLM), which was comprised of mineral water (Volvic®), shredded dry lettuce (1 g for 1.5 L of water, or 0.6 g/L) and a previously prepared liquid culture of the bacterium *Serratia marcescens* (10% of the total volume of the bacterial culture at equilibrium density). The lettuce serves as food resource for the bacterium, which is itself preyed upon by both protist species. Each lineage was transferred to a 100 mL flask as batch cultures for a period of 5 months with 10% bottlenecks. Transfers were done once a week.

##### Final phenotypic tests and data collection

After the experimental evolution phase, we conducted a common garden experiment to eliminate possible transgenerational plastic effects. An aliquot of 5 mL from each evolved stock culture was washed with 10 mL of sterilized shredded lettuce solution medium without *S. marcescens*. This was then centrifuged for 3 minutes under 1100 rpm in a Heraeus Multifuge 3S Centrifuge (Thermo Fisher Scientific Inc., Waltham, Massachusetts, United States) and the supernatant was discarded. The process was then repeated and the same 5 mL aliquots were transferred to 80 Sarstedt tubes of 25 mL with 15 mL of BLM and no stressors. The tubes were kept in a 25 °C incubator for five days, which was expected to be enough to eliminate the plastic responses.

We then prepared an assay with 16-12-well Cellvis glass bottom black plates (Cellvis, Mountain View, California, United States), with each well receiving 3 mL of BLM. One aliquot of 50 µL for *T. thermophila* or 150 µL for *E. gracilis* (due to its longer generation time) of each of the Sarstedt tubes was subsequently added to each well of eight plates that would be submitted to experimental conditions (with stressors), as well as to each well of eight more plates, that would be kept under control conditions (no stressors). The well plates were kept under monoxenic conditions (with *S. marcescens*), with BreatheEasy® sealing membranes (Diversified Biotech, Dedham, Massachusetts, United States) preventing contamination. Additionally, the plates were kept in incubators with constant temperature (36 °C for some lineages of *T. thermophila*, 33 °C for some of *E. gracilis* and 25 °C for the rest; according to the scheme shown in **Table S17**) and a 12:12 hour cycle of light and darkness. The experimental condition plates were submitted to the same treatments used in the first part of the experiment on the ancestor populations, in a way that all possible combinations of stressors were present, including one treatment without any stressors.

The Agilent BioTek Cytation 5 (Agilent Technologies, Santa Clara, California, United States) cell imaging multimode reader was used to record a time-series of cell density in each well for four weeks through 15 second-videos. Moreover, the samples were kept under semicontinuous batch regime, with periodical medium replacements 3 times a week in alternated days for the supplementing of food (10% volume replacement). Sterile conditions were maintained with UV exposure to the plates for 20mins and sterile plastic film to seal wells. Through the videos, the density of cells in each population were estimated using BEMOVI, a package that uses R (version 4.2.3) and ImageJ software (17).

Each population was submitted to the same treatment under which it evolved previously, and under control conditions, so that we could also test if a possible adaptation to the stressors brought fitness cost under no stress. We then ended up with 5 replicates for each of the 8 treatments and species (5 x 8 x 2 = 80). This was replicated for the control condition, adding other 80 lines.

#### Ostreococcus tauri, and Picochlorum costavermella

##### Authors

Edith Vidal, Marc Krasovec and Gwenael Piganeau

##### Study organism

*Ostreococcus tauri* (Mamiellophyceae, Chlorophyta) is a photosynthetic marine picoeukaryote (cell diameter smaller than 2 µm). It was first discovered in the Mediterranean Thau Lagoon (30). *Ostreococcus* species can be dominant during phytoplanktonic blooms in coastal seas, and are the smallest photosynthetic eukaryotic with haploid and very compact genomes of 13 Mb (31, 32). While meiosis has never been observed in the lab, population genomic analyses provided evidence for meiosis and recombination in natural population in *O. tauri* (33).

*Picochlorum costavermella* (Trebouxiophyceae, Chlorophyta) is a marine unicellular green alga (cell diameter 1.4µm). This unicellular coccoid microalga has a broader halotolerance and thermotolerance range as compared to *Ostreococcus*, as *P. costavermella* cultures can stand freshwater to hypersaline media changes (*113*). *P. costavermella* RCC4223 has been isolated in 2011 from the estuary of the coastal river “La Massane” (NW Mediterranean Sea, France). *P. costavermella* has a compact haploid 13.3Mb genome (34).

##### Experimental setup and treatments

In preliminary tests, we performed a literature survey about growth rates as a function of temperature and salinity compared to our standard laboratory control conditions (**Table S18**). This survey suggested a temperature range up to 34 and 35°C and a maximum salinity range up to 50 and 80 g.L^-1^, for *O. tauri* and *P. costavermella*, respectively. We performed an initial test on 32°C and 35°C as well as an increased salinity (**Table S19**). These conditions decreased the growth rate of the strains without imposing high mortality (**Table S20**). One incubator per temperature was used, with a lateral light source. The initial fitness measurements were performed for each species from a clonal culture maintained at the Culture Collection of the GENOPHY team. Fitness was estimated by the growth rate, *g*, the number of divisions per day, *t*, during the exponential growth phase of the culture. *N*(*t*) is the number of cells at day *t*.

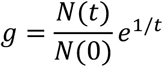

The cell concentration was followed over 20 consecutive days to assess the range of cell concentrations corresponding to the exponential phase in each condition (for details on cell counting protocol see below). The growth rates measured under the control and stress conditions allowed the exponential growth rate to be estimated from *N*(0)= 10^5^ cells to *N*(7) that is 7 days after inoculation. The initial stress response was then estimated by the average growth rate over the seven first days of the experiment (**Table S20**), each well was inoculated with 10^5^ cells at the start of each cycle.

##### Rearing and transferring protocol

We started from a culture that originated from one single cell sorted by dilution. The ancestor culture is thus isogenic, with the exception of variants due to spontaneous mutations. The spontaneous point mutation rate as well as the spontaneous aneuploidy rate is known for these two species (33, 35, 36). The long-term protocol was performed in 24 well plaques (2mL per well). Each well was inoculated with 10^5^ cells every seven days in a total volume of 1mL. Cells count was performed using a Beckman-Coulter CytoFLEX flow cytometer (Beckman Coulter, USA) equipped with three lasers emitting light in spatially separated pathways at 405 nm (80 mW), 488 nm (50 mW), and 561 nm (30 mW). Subsequent estimations of cell concentrations were calculated using Becton-Dickinson TrucountTM beads. We used the software CytExpert (v. 2.3.1.22), generating listmode (FCS 3.0) files. Phytoplankton cells were analysed unstained and enumerated based on chlorophyll a fluorescence (692 nm maximum emission) and side scatter (SSC). The concentration of cells was used to estimate the volume required to transfer 10^5^ cells from each cell line into fresh media at each transfer. At the start of the experiment, the cultures were left 7 days under each treatment prior to the first measurement. The number of cells was then estimated every week (corresponding to one cycle) in their own evolutionary treatment for the complete duration of the experiment.

In total we established 6 replicates per species and condition, corresponding to a single line in a 24 wells plate. For *P. costavermella* the experiment was restarted (**Table S20-21**, Exp1-Exp2) because of too harsh combined stress conditions, so that we have: 2 species (*P. costavermella* and *O. tauri*) x 4 treatments x 6 biological replicates for 17 cycles (119 days), and 1 species (*P. costavermella*) x 4 treatments x 6 biological replicates for 12 cycles (84 days).

##### Final phenotypic tests and data collection

In the last cycle, cells were transferred once more in their own evolutionary treatment and counted by flow cytometry as described, but they were not assayed in control conditions in the absence of stressors.

#### Paramecium caudatum

##### Authors

Oliver Kaltz, Marie-Ange Devillez

##### Study organism

The freshwater ciliate *Paramecium caudatum* represents a classic model system for genetic, developmental and physiological study (37). It has also been used for ecological studies (38, 39) and experimental evolution (40, 41). Previous work has shown that *P. caudatum* is highly sensitive to change in temperature and osmotic stress (37, 41–44). For the present study, we used the strain KS3, collected in Germany (45) and kept in the OK lab since 2019. Under standard conditions, we keep *Paramecium* mass cultures at 23°C, in a lettuce medium, made from organic lettuce, Volvic^TM^ mineral water, and supplemented with the bacterium *Serratia marcescens* as food resource (46).

##### Experimental setup and treatments

From a series of pilot experiment, testing the KS3 strain over a range of temperatures and salt concentrations, we determined the conditions for the long-term experiment, consisting of a control treatment (23°C), a single-temperature treatment (30°C), a single-salt treatment (2g/L of NaCl), and a combined treatment (28°C and 2g/L of NaCl).

From a KS3 stationary-phase mass culture at 23°C, we established replicate cultures of 20mL (in 50mL tubes), with a starting density of 100 individuals and exposed to the four above treatments (control, single stress, combined stress). For the temperature treatments, tubes were placed in incubators set to the appropriate temperature. Over the course of 9 days, population density was tracked in 1-to 2-day intervals, by removing small samples of up to 300µL and counting the number of individuals under a dissecting microscope. After 4 days, we replenished the cultures, by adding 1mL of medium. For each treatment, we established four replicates (16 replicates).

##### Rearing and transferring protocol

From a KS3 stationary-phase mass culture at 23°C, we established replicate cultures of 150mL (in 250mL glass bottles) with initially c. 1000 individuals. These were assigned to the four treatments, with 5 long-term replicates per treatment (= 20 lines in total). For the temperature treatments, tubes were placed in incubators set to the appropriate temperature. The long-term culture protocol consisted of weekly transfers of 30mL of culture to a new bottle, to which we added 120mL of freshly prepared medium, adjusted for the appropriate salt concentration. Over the course of 1 year, 48 transfers were accomplished, corresponding to ≥96 population doublings (i.e., asexual generations).

##### Final phenotypic tests and data collection

At the end of the long-term experiment, each evolved line was tested in its own long-term environment and in the control environment. These assays were run in 50mL tubes, and initiated by placing 200 individuals of a given long-term line in 20mL of lettuce medium, adjusted to the appropriate salt concentration. The tubes were then transferred to the appropriate temperature condition. Densities were tracked over 13 days, as described above.

#### Pseudomonas aeruginosa

##### Authors

Jeanne Hamet, Stéphanie Bedhomme

##### Study organism

*Pseudomonas aeruginosa* (PA) is a gram-negative bacterium and opportunistic pathogen, responsible for a growing number of nosocomial infections due to its frequent resistance to antibiotics (47). It is a ubiquitous organism, with a preference for environments close to human activity (48). PA can be observed in planktonic form, but most of the time, it is observed in biofilms (49). This habitat enables it to protect itself from biotic and abiotic environmental stresses (50). In standardized laboratory conditions, at 37°C and classical culture broth (LB Broth), they asexually divide with a generation time of 30 minutes (51). Strain PAO1 (Taxonomy ID: 208964) was chosen here because its genome has been sequenced and it is a model laboratory strain (52). This strain was isolated from a wound (53).

##### Experimental setup and treatments

We designed four evolutionary treatments, a control, copper stressor, salt stressor and salt plus copper stressor. To decide on the concentrations of salt and copper in the three stress treatments, growth curves were realised over a range of salt and copper stress concentrations in a crossed design allowing us to define a 2-dimension niche. A pre-culture was set up from the glycerol stock of the ancestral strain (PAO1) and cultivated in 4mL of LB Broth at 37°C with 220 rpm orbital shaking overnight. 2µL of the pre-culture was then transferred to each well of a microtiter plate containing 200µL of LB broth and various concentrations of salt (0, 0.1, 0.2, 0.3, 0.4, 0.5, 0.6, 0.7, 0.8, 0.9 and 1M Sodium chloride) and copper (0, 0.001, 0.002, 0.003, 0.004, 0.005, 0.006 and 0.007M copper(II) sulfate pentahydrate) in triplicate. Sodium chloride and copper(II) sulfate pentahydrate were ordered from Sigma Aldrich. Growth curves (density over time) were obtained monitoring the optical density at 550nm (OD550) during 24h with a Tecan plate reader following the program indicated in **Table S22**. For each simple and double stress treatment, we chose the experimental evolution stressor concentrations causing a reduction in absolute fitness close to 50% (details in **Table S23**).

Prior to the experiment, a pre-culture was set up from the glycerol stock of the ancestral strain (PAO1) and cultivated in 4mL of LB Broth at 37°C with 220 rpm orbital shaking for 24 hours. All experimental evolution lines are thus derived from a clonal ancestor. To begin the experimental evolution, we added 12µL of bacterial culture to 1188µL of medium in a Deepwell plate. 12 replicate populations were set up for each of the four evolutionary treatments. From these initial mixes, 200µL were transferred to a microtiter plate. Treatments were distributed in a checkerboard pattern with the same positions on each plate, and evolution medium in alternating wells to identify medium contamination or high levels of cross-contamination during transfer. The Deepwell plate was incubated for 24 hours at 37°C, with 220 rpm orbital shaking. The microtiter plate was incubated at 37°C, with shaking and OD550 was recorded following the cycle described in Table S8.1.

##### Rearing and transferring protocol

On following days during experimental evolution, 10µL of each population was transferred to a new plate containing 990µL of fresh medium (10^−2^ dilution, giving approximately 6.6 generations/day). When a population had low turbidity, we transferred 20µL of that population (**Table S24**). Each transfer started a new cycle.

Populations were maintained for 26 cycles, resulting in approximately 173 generations of experimental evolution. At cycle 10 and 26, one line in the salt and copper treatment (S3) became extinct despite transferring 20µL of the evolved population, leaving 46 lines for the final phenotypic assays. Frozen archives of the evolving populations were stored at cycle 1, 4, 10, 16, 22 and 26 in the form of glycerol stocks. Each week, we mixed 280µL of homogenate from each evolved population with 350µL of 70% glycerol saline solution, and stored at -70°C.

##### Final phenotypic tests and data collection

At cycle 26, we closely followed the growth of the evolutionary lines in their own environment. We added 2µL of the populations to 198µL of their evolutionary medium in a microtiter plate, and growth curves were obtained by monitoring the OD550 as previously described. We also tested the lines in control condition. We defrosted the glycerol stock from cycle 26, and 10µL of each evolutionary line was transferred to 990µL of their evolutionary medium and incubated at 37°C, with 220 rpm orbital shaking for 24 hours. Afterwards, we added 2µL of each population to 198µL of control medium in a microtiter plate. The populations that did not grow during the final tests are listed in **Table S25**.

#### Ralstonia pseudosolanacearum

##### Authors

Delphine Capela, Alice Guidot, Philippe Remigi

##### Study organism

*Ralstonia solanacearum* is a soil bacterium belonging to the beta subclass of Proteobacteria. It is a species complex responsible for the bacterial wilt disease on more than 250 plant species in tropical and subtropical countries (54). The strain GMI1000 isolated from Tomato in French Guyana (55) was used as a model strain in many different studies and its genome was the first *R. solanacearum* genome completely sequenced (56).

##### Experimental setup and treatments

Liquid cultures of GMI1000 strain were carried out in MP synthetic medium which composition is as follows (g liter^-1^): FeSO_4_ 7H_2_O, 1.25 10^-4^; (NH_4_)_2_SO_4_, 0.5; MgSO_4_ 7H_2_O, 0.05; KH_2_PO_4_, 3.4. The pH was adjusted to 6.5 with KOH. The MP synthetic liquid medium was supplemented with L-glutamine(10 mM) and oligo-elements (ZnSO4 7 H2O 55 µg.L^-1^, H_3_BO_3_ 28.5 µg.L^-1^, MnCl2 4·H2O 12.6 µg.L^-1^, CoCl2 6·H2O 4 µg.L^-1^, CuSO4 5·H2O 3.9 µg.L^-1^, (NH4)6Mo7O24 4·H2O 2.8 µg.L^-1^) (57).

NaCl was added at final concentrations of 75 or 150 mM as needed (see below). The standard rearing temperature of *R. solanacearum* growth in laboratory conditions is 28°C. In preliminary tests conducted before the start of the long-term experiments, we assayed growth at higher temperatures. We observed that growth rate was strongly reduced at 37°C (about 5 folds reduction) but was completely inhibited at 38°C. We therefore decided to keep 37°C as the high temperature stress condition. The impact of NaCl addition on GMI1000 growth rate was more linear. The addition of NaCl at a concentration of 150 mM in the bacterial culture reduced the growth rate (about 5 folds reduction). We therefore decided to use 75 and 150 mM NaCl as salt stress conditions at 37 and 28°C, respectively. The ancestral strain GMI1000 was revived from glycerol stock on plates containing rich agar medium (bacto peptone 10 g.L^-1^, casamino acids 1 g.L^-1^, yeast extract 1 g.L^-1^) for 2 days at 28°C. One individual colony was used to inoculate 100 mL Erlenmeyer flasks containing 20 mL MP-glutamine liquid medium and was incubated overnight at 28°C. After 24 h, optical density at 600 nm (OD_600_) was measured, and the pre-culture was used to inoculate 100 mL Erlenmeyer flasks containing 20 mL MP-glutamine (at starting OD_600_ = 0.001) in four different evolutionary treatments: control (28°C, 0 mM NaCl), saline stress (28°C, 150 mM NaCl), temperature stress (37°C, 0 mM NaCl), and the combination of two stresses (37°C, 75 mM NaCl).

##### Rearing and transferring protocol

Five independent cultures were inoculated for each treatment. After 24 hours of growth in these conditions, OD_600_ was measured, the cultures were used to re-inoculate new cultures in the same conditions (starting at OD_600_ = 0.001), and aliquots of the cultures were stored at -80°C in the presence of 20% glycerol. The same protocol was applied for 15 cycles.

##### Final phenotypic tests and data collection

We measured the growth rate of all evolved lines in control and in their own evolutionary condition. The initial growth of the ancestor was tested as well in all four conditions (three stress and control conditions). To standardize these assays, the ancestor strain and the evolved populations were first restarted from frozen glycerol stocks in their evolution medium and incubated for 24 hours at 28°C. These precultures were used to generate, for each population, multiple 100 µl aliquots of bacterial cultures containing 20% glycerol that were stored at -80°C. To initiate a phenotypic test, one entire frozen aliquot per population was thawed in 1ml of its evolution medium and 50 µl or 100 µl of these suspensions were used to inoculate an overnight pre-culture in the evolution conditions (ancestral strain GMI1000 was grown in control). On the next day, the pre-cultures were used to inoculate fresh 20 mL cultures either in control or in evolution medium (ancestral strain GMI1000 was grown in control as well as in all stress conditions). The starting OD_600_ of these cultures were adjusted between 0.000003 and 0.01, so that all populations reached an OD_600_∼0.05-0.1 after ∼16 hours (the next morning), from which point OD_600_ was monitored every ∼2 hours for ∼8-10 hours. After 24 hours, evolved populations grown in control condition were used to inoculate new 20 mL cultures in their evolution treatment and their growth was followed as during the first round of culture, in order to estimate the effect of pre-culture condition on the growth in the stressful environment. The whole experiment was performed at least three times independently for each of the lines.

#### Saccharomyces cerevisiae (all strains)

##### Authors

Apolline Vedrenne, Diego Segond, Delphine Sicard and Thibault Nidelet

##### Study organism

*Saccharomyces cerevisiae* is the most known of yeasts also called baker yeast (58). We worked with three strains: EC1118 (a commercial strain that is used as reference in oenological research) and M1OA, a strain from our collection and collected from vineyards around Montpellier. Under our experimental conditions, yeast divides asexually by budding with a generation time of around 2h (59).

##### Experimental setup and treatments

Two different strains of Saccharomyces cerevisiae (EC1118, and M1OA) were inoculated into deep-well microplates with a starting population of 10^6^ cells/mL. Each strain was inoculated in 4 different media: an unstressed medium with a temperature of 28°C and no copper, a high-temperature stress medium (33°C) with no copper, a medium with a temperature of 28°C but stressed by a high copper concentration (1g/L) a doubly stressed environment with a high temperature of 33°C and a high copper concentration (1g/L). The based medium was a synthetic must that we use for all our experiments (Bely *et al.*, 1990). It contains mainly 200g of sugars and 200mg of assimilable nitrogen sources, as well as phytosterols and other essential nutrients. These conditions have been chosen after preliminary experiments.

##### Rearing and transferring protocol

For each strain and each medium, three independent lineages were started. Twice a week, 80% of the medium was scattered and replaced by fresh medium. The experiment was carried out for 26 cycles (93 days). Once a week, yeast population growth was determined by cytometry, using an Attune NxT (Life Technologies) with 200 μL/min flux. At the end of the evolutionary phase, we froze all our evolved lines (populations).

##### Final phenotypic tests and data collection

We thawed all the evolved lines, including the ancestral populations, and grew them for 24 days in yeast extract peptone dextrose (YPD), a neutral environment. We then inoculated in microplates (at 10^6^ cells/ml) each of the lines, including the ancestors, in triplicate in their own evolutionary conditions (no stress, high temperature, high copper concentration and double stress), and in control conditions in the absence of stressors. Ancestors were tested in the four evolutionary conditions. As previously illustrated, we used cytometry and measured population density (OD) during 24h.

#### Sinorhizobium meliloti

##### Authors

Laurent Sauviac and Claude Bruand

##### Study organism

*Sinorhizobium meliloti* is a soil bacterium belonging to the alpha subclass of Proteobacteria. It is the natural nitrogen fixing microsymbiont of legume plants of the *Medicago* and *Melilotus* genera, including the forage crop alfalfa (*Medicago sativa*) and the model legume *Medicago truncatula*. The strain *Sinorhizobium meliloti* CBT2801 is a derivative of the strain Rm2011 (60).

##### Experimental setup and treatments

Liquid cultures of *S. meliloti* were carried out in Vincent minimal medium: VMM; 7.35 mM KH_2_PO_4_, 5.74 mM K_2_HPO_4_, 1 mM MgSO_4_, 456 µM CaCl_2_, 35 µM FeCl_3_, 4 µM biotin, 48.5 µM H_3_BO_3_, 10 µM MnSO_4_, 1 µM ZnSO_4_, 0.5 µM CuSO_4_, 0.27 µM CoCl_2_, 0.5 µM NaMoO_4_, 10 mM sodium succinate, 18.7mM NH_4_Cl pH 7, always supplemented with 100 µg.ml^-1^ streptomycin (Sm). NaCl was added at final concentrations of 250 or 400 mM as needed (see below). The standard rearing temperature of *S. meliloti* growth in laboratory conditions is 28°C. In preliminary tests conducted before the start of the long-term experiments, we assayed growth at higher temperatures. Bacteria grew better at 37°C than at 28°C, their growth rate was strongly reduced at 40°C and above, as was the maximal culture density. Moreover, addition of even low salt concentrations at 40°C led to a complete cessation of growth. We observed that at 39.5°C, the strain grew more slowly than at 28°C (∼2-fold reduction), while reaching a normal maximal culture density, and supporting the addition of salt (see below). We therefore decided to keep 39.5°C as the high temperature stress condition. NaCl addition at concentrations of 200-250 mM had little effect on growth at 28°C, which was significantly reduced (∼2-fold) at concentrations of 350 mM and above. At 39.5°C, addition of 250 mM NaCl further slowed down bacterial growth (∼60% reduction) without affecting maximum cell density. We therefore decided to use 250 and 400 mM NaCl as salt stress conditions at 39.5 and 28°C, respectively.

Five independent clones isolated on LB rich medium supplemented with 2.5 mM MgSO_4_, 2.5 mM CaCl_2_ and 100 µg.ml^-1^ Sm were used to inoculate tubes containing 5 mL VMM and incubated overnight at 28°C under agitation. After 24h, optical density at 600 nm (OD_600_) was measured, and cultures were used to inoculate 100 mL Erlenmeyer flasks containing 20 mL VMM (at starting OD_600_ = 0.001) in four different evolutionary treatments: control (28°C, 0 mM NaCl), saline stress (28°C, 400 mM NaCl), temperature stress (39.5°C, 0 mM NaCl), and the combination of two stresses (39.5°C, 250 mM NaCl).

##### Rearing and transferring protocol

After 24 hours of growth in the four evolutionary conditions, OD_600_ was measured, the cultures were used to re-inoculate new cultures in the same conditions (starting at OD_600_ = 0.001), and aliquots of the cultures were stored at -80°C in the presence of 20% glycerol. The same protocol was applied for 20 cycles.

##### Final phenotypic tests and data collection

We measured the growth rate of all evolved lines in control and in their own evolutionary condition. The initial growth of the ancestor was tested as well in all four conditions (three stress and control conditions). The ancestor strain as well as populations evolved for 20 cycles were first restarted from frozen glycerol stocks in 5 ml tubes of VMM (start OD_600_= 0.001) incubated for 24 hours at 28°C. These precultures were used to inoculate 100 ml flasks containing 20 ml VMM at 28°C. After ∼16 hours (overnight) OD_600_ was monitored every ∼2 hours for ∼8-10 hours, to generate the first dataset (fitness of each line in control conditions). 24 hours post-inoculation, 20 mL cultures were again inoculated from each of these cultures (start OD_600_= 0.001), but then incubated in the evolutionary condition of the tested line, except for the ancestor which was grown in the four evolutionary conditions. After this first cycle of 24 hours, this step was repeated, and after 16 hours of growth, OD_600_ was monitored every ∼2 hours for ∼8-24 hours, to generate the second dataset (fitness of each line in their own evolutionary treatment). The whole experiment was performed at least three times independently for each of the lines.

#### *Tetrahymena thermophila* (strain D16, D19, D21, D3)

##### Authors

Delphine Legrand, Staffan Jacob, Michèle Huet, Hervé Philippe

##### Study organism

*Tetrahymena thermophila* is a free-living ciliate living from northern America found in freshwater ecosystems. It is a main actor of aquatic trophic chains, as a grazer of bacteria and prey of mesoplanktonic species (61). It is a historical model species in cellular and molecular biology, which has led to major discoveries like identifications of dynein or telomere structure (28, 62), and has more recently turned into an experimental model in ecological and evolutionary studies, for instance in the fields of dispersal, adaptation and phenotypic plasticity (*e.g.*, (63–65)). The species alternates phases of sexual and asexual reproduction in nature, and can be long-maintained as clonal populations in laboratory. Under standard conditions (24-well plates, 23°C, aqueous 0.3X PPYE medium, 0.6% of Proteose Peptone and 0.06% Yeast Extract), the asexual generation time is about 4-6 hours.

Here, we used six clonally reproducing *T. thermophila* strains, namely D3, D8, D10, D16, D19 and D21, initially collected by Pr Paul Doerder and maintained in our laboratory for ∼20 years. These strains were recently sequenced (66), which revealed close genomic proximity between D8, D19 and D21, a probable recent cross between a D8/D19/D21-like strain and a divergent strain leading to D16, and diverging genomes for D3 and D10, the only two strains whose origin seems to match the initial sampling of Pr P. Doerder.

##### Experimental setup and treatments

To obtain the six ancestors’ cultures, we filled six 1L Erlenmeyers with 100mL of PPYE 0.3X in which we inoculated 100µL of 7-old stock cultures. Erlens were placed in an incubator at 23°C and gently shaken during seven days to obtain sufficient initial densities to start the experiment.

We then established four evolutionary treatments (Control, Salt, the antibiotic Chloramphenicol, Salt + Chloramphenicol) by manipulating salinity and chloramphenicol concentrations (**Table S26**), two pollutants that incur serious threats to aquatic ecosystems (67, 68). Pollutants’ concentrations were chosen based on pilot experiments in which we exposed the strains to a gradient of salt concentrations (from 0 to 20g/L) and a gradient of chloramphenicol concentrations (from 0 to 500mg/L). Treatment concentrations (5g/L for NaCl and 40mg/L for chloramphenicol) were fixed so that they reduced but not stopped the growth over all strains. As strains have their own growth capacities, the initial growth reduction was strain-specific (see details on growth measurements below).

To initiate experimental evolution, we filled three 24-well plates with 2mL of each of the four evolutionary treatments, for a total of 12 managed plates. Each strain was exposed to the four evolutionary treatments in six replicates, leading to 144 evolved populations (6 strains x 4 treatments x 6 replicates = 144). Each plate contained two replicates of the six strains, meaning that 12 wells were left empty in a checkerboard configuration to limit cross-contaminations. We randomized the position of each strain between the three plates containing the same evolutionary treatments to control for the effect of the well position. To initiate the experiment, we inoculated 100µL of ancestral populations. To estimate initial growth (fitness), we used growth rate as the fitness proxy. At the time of evolved populations inoculation in 24-well plates, we parallelly inoculated 10µL (∼100 to 1,000 cells depending on the strain) of ancestral cultures in five technical replicates into 96-well plates pre-filled with 250 µL of each of the four treatments (with randomized position of both strains and treatments). Population growth was quantified by measuring Optical Density (OD) at 450nm every four hours during 20 days (sufficient delay to reach the stationary phase for each strain x treatment condition) using a microplate reader (Tecan Infinite spectrophotometer). OD at 450nm is positively correlated with *T. thermophila* cell density in our laboratory conditions (*e.g.*,(64)).

##### Rearing and transfer protocol

Evolved populations were maintained in the 24-well plates during five months and a half, with plate transfer using 5% of populations (100µL over the 2mL of each well) every seven days during two months (10 cycles), and then every 14 days until the end of the experiment (7 cycles). We indeed observed that some evolved populations had critical densities requiring longer growth periods to ensure their survival. In total, we thus performed 17 cycles. Under standard conditions, the duration of the experiment corresponds to ∼800 asexual generations.

##### Final phenotypic tests and data collection

During experimental evolution, two types of data were collected:

i) growth rates by spectrophotometry following the protocol described above for the ancestral populations at cycles 17 (final time, 168 days). We placed 10µL of the 144 evolved populations in the four treatments immediately after pipetting (144 evolved lines x 4 treatments x 5 replicates = 2,880 growth assays). We further assayed the obtained 144 evolved lines in control conditions.
ii) cell densities seven days after inoculation in fresh plates by video analysis at cycles 1 (ancestors) and 17 (final time, 168 days). To do so, we pipetted 10μl of each evolved population into multichambered counting slides (Kima precision cell) and immediately took 15s videos from each chamber under darkfield macroscopy to count the number of cells using the BEMOVI R-package (17). These videos were used to estimate cell densities in the treatment of evolution, so as to check for population extinctions.

From videos and complementary observation of whole wells under a macroscope, we observed that all populations survived. However, some had very low densities, zero to three cells in the 10µL sampled for video analyses, which corresponds to no more than ∼600 cells at the population scale. At such low densities after seven days of growth, we are unable to properly assess growth rates using spectrometry (cell number below the detection threshold). This was the case of all D10 replicates for the combined stressors. We thus excluded this strain from the final analyses. Besides, ancestors of strain D8 did not grow in the control treatment, meaning that we could not assess its initial fitness. We suspect a pipetting omission when performing the growth assays of D8 ancestor. We thus excluded D8 from final analyses, despite having evolved all populations correctly. In the end, we only kept strains D3, D16, D19 and D21 for the final data analyses.

#### Vibrio aestuarianus

##### Authors

Fiona Elmaleh, Gaelle Courtay, Guillaume M. Charrière

##### Study organism

*Vibrio aestuarianus* is a gram-negative halophile γ-proteobacteria. It is an actively studied oyster pathogen that has colonized Europe since the second half of the 20^th^ century (69). The pathogenic subspecies *Vibrio aestuarianus francensis,* like the 12/016 strain, is a primary pathogen for adult oysters and was shown recently to carry a wide array of copper-resistance genes when compared to the non-pathogenic environmental stain, like *Vibrio aestuarianus* U29 (69).

##### Experimental setup and treatments

To start our experiment, we used 2 strains (12/016 and U29) isolated from diseased oysters or within the environment respectively. *V. aestuarianus* strains were cultivated in LB-0.3M NaCl at 25°C under agitation for all the experiments. Four evolutionary treatments, a control, copper stress, salt stress and salt plus copper stress were chosen. To decide on the concentrations of salt and copper in the three stress treatments, growth curves were performed over a range of salt and copper stress concentrations in a crossed design allowing to define a 2-dimension niche. A pre-culture was set up from the glycerol stock of the ancestral strains (12/016 or U29) and cultivated in 4mL of LB-NaCl Broth at 25°C with 150 rpm orbital shaking overnight. After adjusting OD600 at 1, 2µL of the pre-culture was then transferred to each well of a microtiter plate containing 200µL of LB broth and various concentrations of sodium chloride and/or copper (copper(II) sulfate pentahydrate) in triplicate. Sodium chloride and copper(II) sulfate pentahydrate were ordered from Sigma Aldrich. The optical density at 600nm (OD600) was monitored during 24h with a plate reader with 5 min double orbital agitation just before reading, which were acquired every 20 min.

These experiments allowed us to establish 4 evolutionary treatments in 3 replicates (**Table S27**) by testing different concentrations of CuSO_4_ and NaCl. In the control non-stressful treatment, we used LB-0.3M NaCl. For the 3 stressful treatments (copper, salt, and the combination of the two), pilot experiments indicated that for 12/016 strain, CuSO_4_ at 2.5M, NaCl at 0.84M, and 1.25M CuSO_4_ + 1.125M NaCl allowed to obtain about 50% initial reduction of growth. For U29 strain, CuSO_4_ at 1.25M, NaCl at 0.84M, and 0.625M CuSO_4_ + 0.84M NaCl allowed to obtain about 30% initial reduction of growth rate.

For each strain, the experimental microplates were set up with triplicate for each strain and for each stress condition. Microplate wells were filled with 180µl of media and inoculated with 20µl of culture at OD600 equal to 1 to obtain a final OD600 at 0.01 and incubated at 25°C under double orbital agitation in a microplate reader. This gave us a total of 24 evolutionary replicates: 2 strains x 4 stressor conditions x 3 replicates (**Table S27**).

##### Rearing and transferring protocol

To start each experiment, a pre-culture was set up from a colony and cultivated in 4mL of LB-0.3M NaCl at 25°C with 150 rpm orbital shaking for 24 hours. All experimental evolution lines were derived from the same clonal ancestor. For 12/016 strain, evolution lines were propagated every 48h (to achieve stationary phase) by 1/100 dilution in a new microplate. For U29, evolution lines were propagated every 12h (to achieve stationary phase) by 1/100 dilution in a new microplate. Readings of OD600 were acquired every 20min for each well. After propagation, each microplate was cryopreserved at -80°C after adding 15% of glycerol per well. For 12/016 strain, the experiment lasted for 46 days (22 cycles), corresponding to ∼ 180 generations. For U29 strain, the experiment lasted 8 days (16 cycles), corresponding to ∼ 128 generations.

##### Final phenotypic tests and data collection

As described above, initial and final growth rate was obtained in its own evolutionary treatment via spectrophotometry at cycles 1 and 22 for strain 12/016, and at cycle 1 and 16 for strain U29. In addition, after defrosting the glycerol stock from the last cycle, 20µL of each evolutionary line was transferred to 180µL and assayed in the control media (LB-0.3M NaCl, incubated at 25°C), with double orbital shaking for 24 hours and OD600 recorded every 20min.

### Organismal complexity

We investigated with linear multilevel models whether direct evolutionary responses could be explained by species complexity, as approximated by the number of protein-coding genes (after GenBank annotation), or the number of protein families (after Pfam annotation, (*51*)). For *Colpidium striatum*, no data were available for protein-coding genes, and it was therefore excluded for this analysis (x13 species and x22 strains). Data for *Colpidium striatum*, *Dunaliella salina*, *Euglena gracilis*, and *Picochlorum costavermella* were not available for the number of protein families, and they were removed from this analysis (10 species). For *Paramecium caudatum* and *Vibrio aestuarianus* data were also not available, and we used the 2 closely related species, *Paramecium tetraurelia* and *Vibrio cholerae* respectively.

Species with more protein-coding genes or protein families (as proxies for the number of functions, and therefore for organismal complexity (*51*)) might be less constrained in their evolutionary response to combined stressors. However, we found no signal for an association between direct response to selection and the number of protein-coding genes (after GenBank annotation) in a species, neither for single nor for combined stressors (LMM, Simple: -0.039 [-0.194, 0.118], Complex: 0.007 [-0.146, 0.165]; **Fig. S13A**). We also found no relationship (**Fig. S13B**) when using protein families as explanatory variable (across the 8 species for which this information was available; information from the close relatives *Paramecium tetraurelia* and *Vibrio cholerae* was used for the 2 species *P. caudatum* and *V. aestuarianus*): posterior slopes for simple and complex were both not different from 0 (LMM, Simple median and 95% Compatibility Intervals: 0.042 [-0.224, 0.310]; Complex: 0.022 [-0.224, 0.289]).

## Figures

**Fig. S1.**
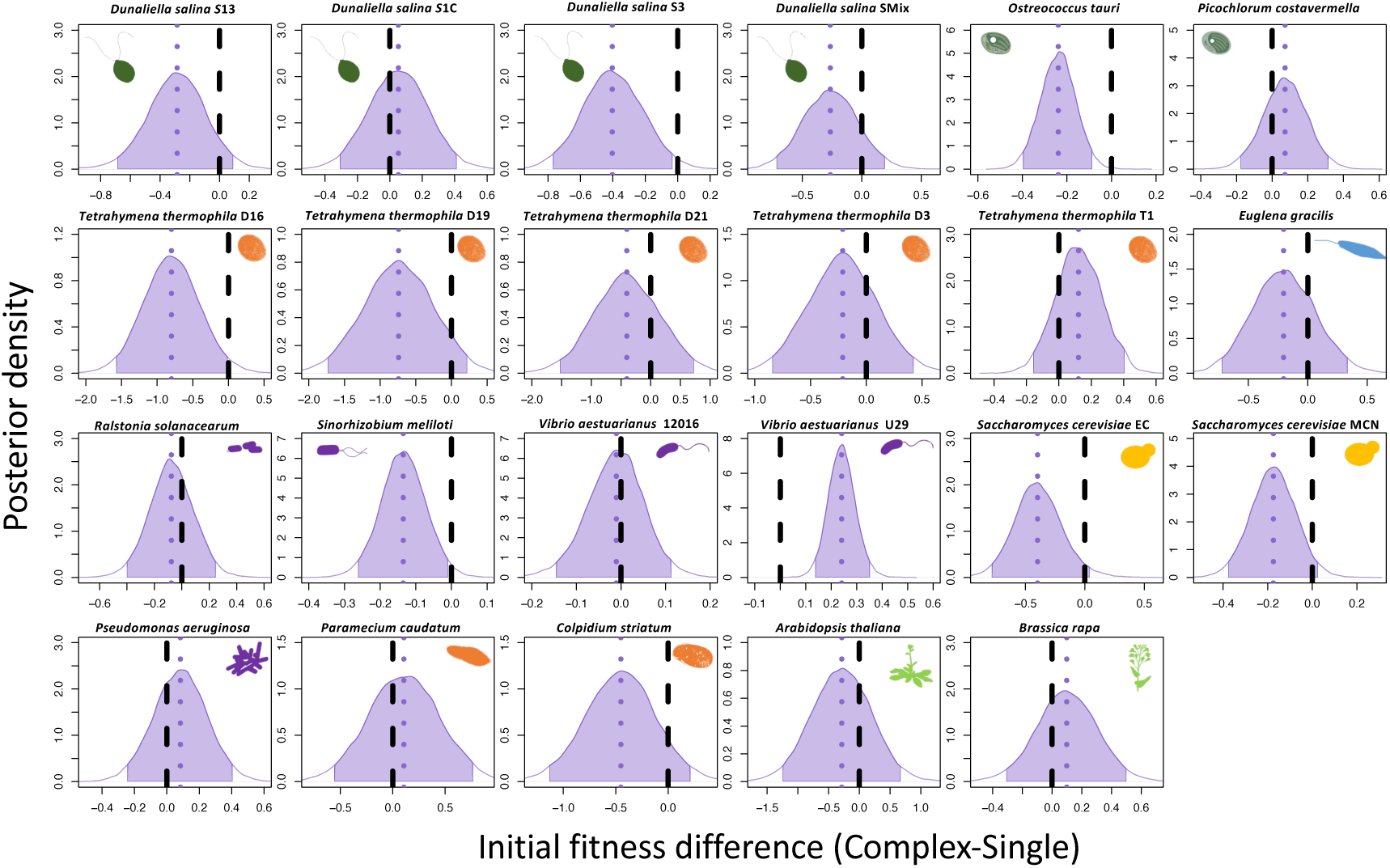
Posterior distributions, for each of the 14 species, of the difference between combined minus single stressors of the initial fitness reduction. The dashed purple lines visualize the median posterior values, and the shaded area the 95% compatibility intervals of the posterior distributions. Different strains of the same species are indicated next to their scientific name, with their silhouette repeated. Overlap of the distributions with the reference (black dashed line at zero values) indicates comparable initial reduction in fitness for simple vs. complex stress treatments.

**Fig. S2.**
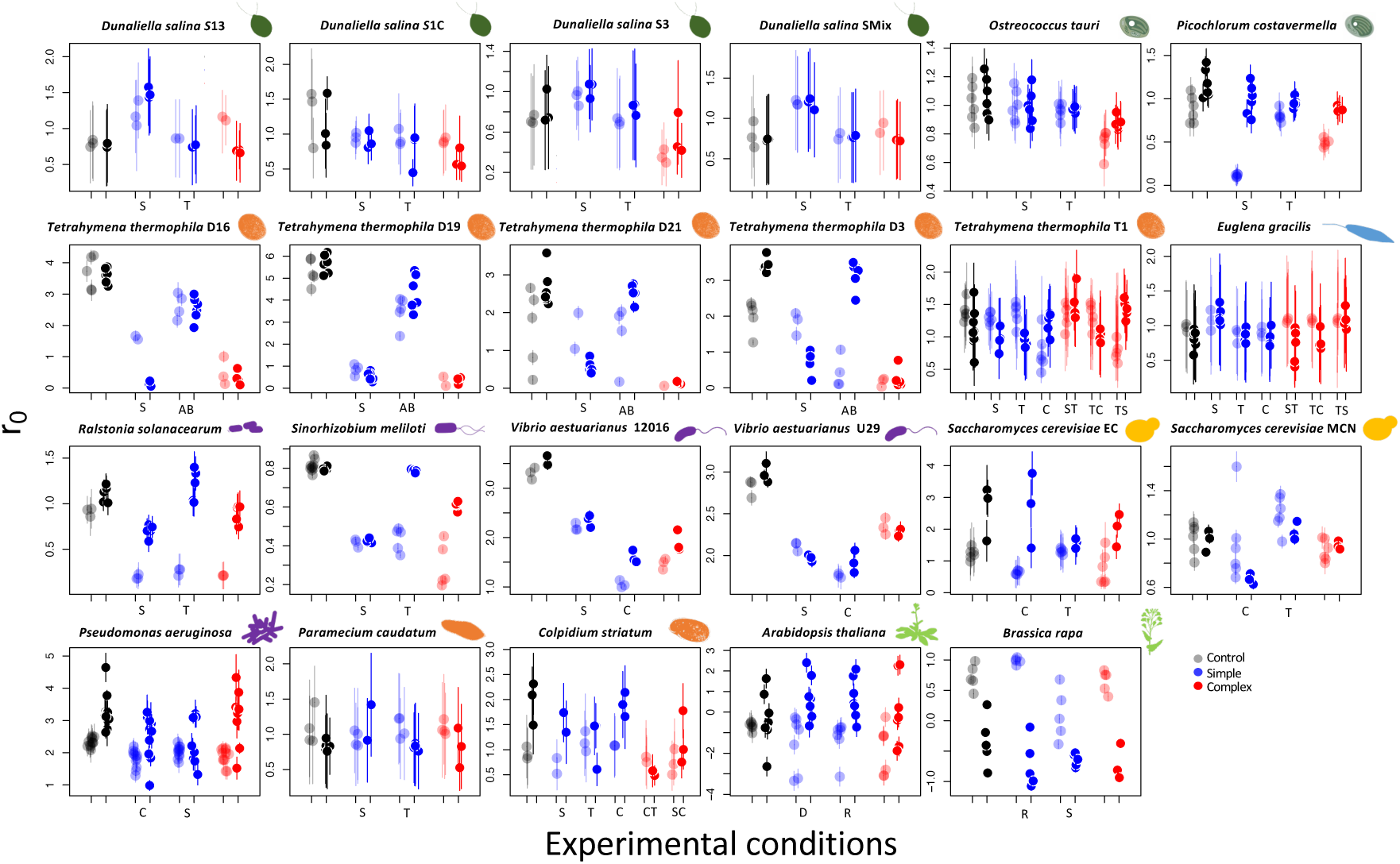
Fitness estimates (*r_0_*) for each of the 14 species (silhouette) assayed in their own evolutionary conditions: control (grey), single stressors (blue) or combined stressors (red). Shaded points are fitness estimates after the initial transfer to the experimental conditions, while thick points are the final fitness estimates at the end of the evolutionary experiments. The corresponding initial and final fitness estimates are therefore coupled by experimental treatments. Points and bars represent the fitness estimate and standard deviation for a biological replicate. Different strains of the same species are indicated next to their scientific name, with their silhouette repeated. Capital letters indicate the identity of the single stressors, S (Salt), T (Temperature), C (Copper), AB (Antibiotics), D (Disturbance), R (Resources), or of the combined stressors when more than one combination was tested (i.e., ST, TC, TS).

**Fig. S3.**
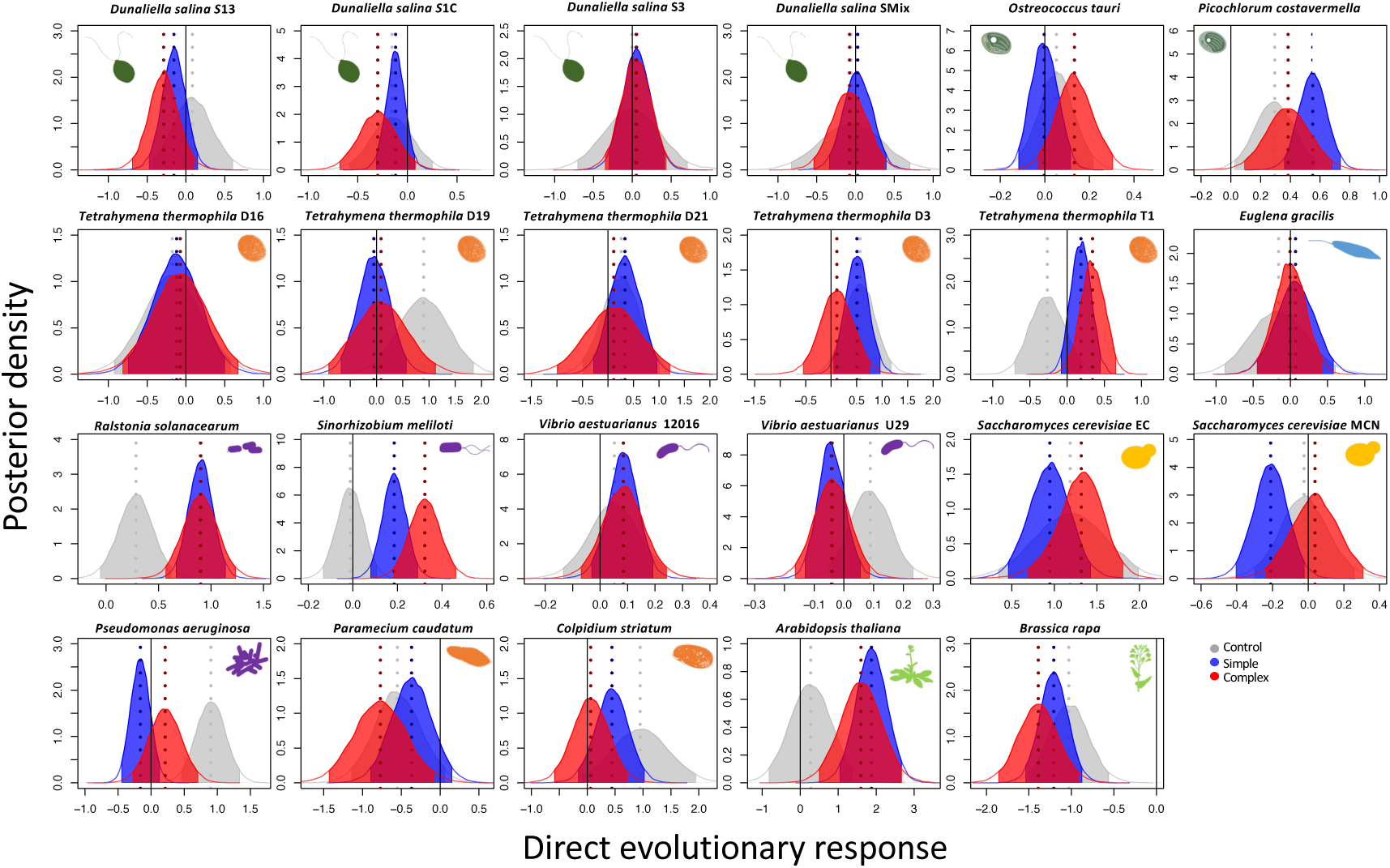
Posterior distributions for direct evolutionary responses of each strain. Positive values represent a gain in fitness over the evolutionary experiment. In grey is the control in the absence of stressor, in blue and red the single and combined stressors respectively. The dashed lines visualize the median posterior values, and the shaded area the 95% compatibility intervals. The posteriors were obtained by calculating the difference between final and initial fitness for the control, single and combined stressors evolutionary treatments respectively (14 species and 23 strains, complete dataset).

**Fig. S4.**
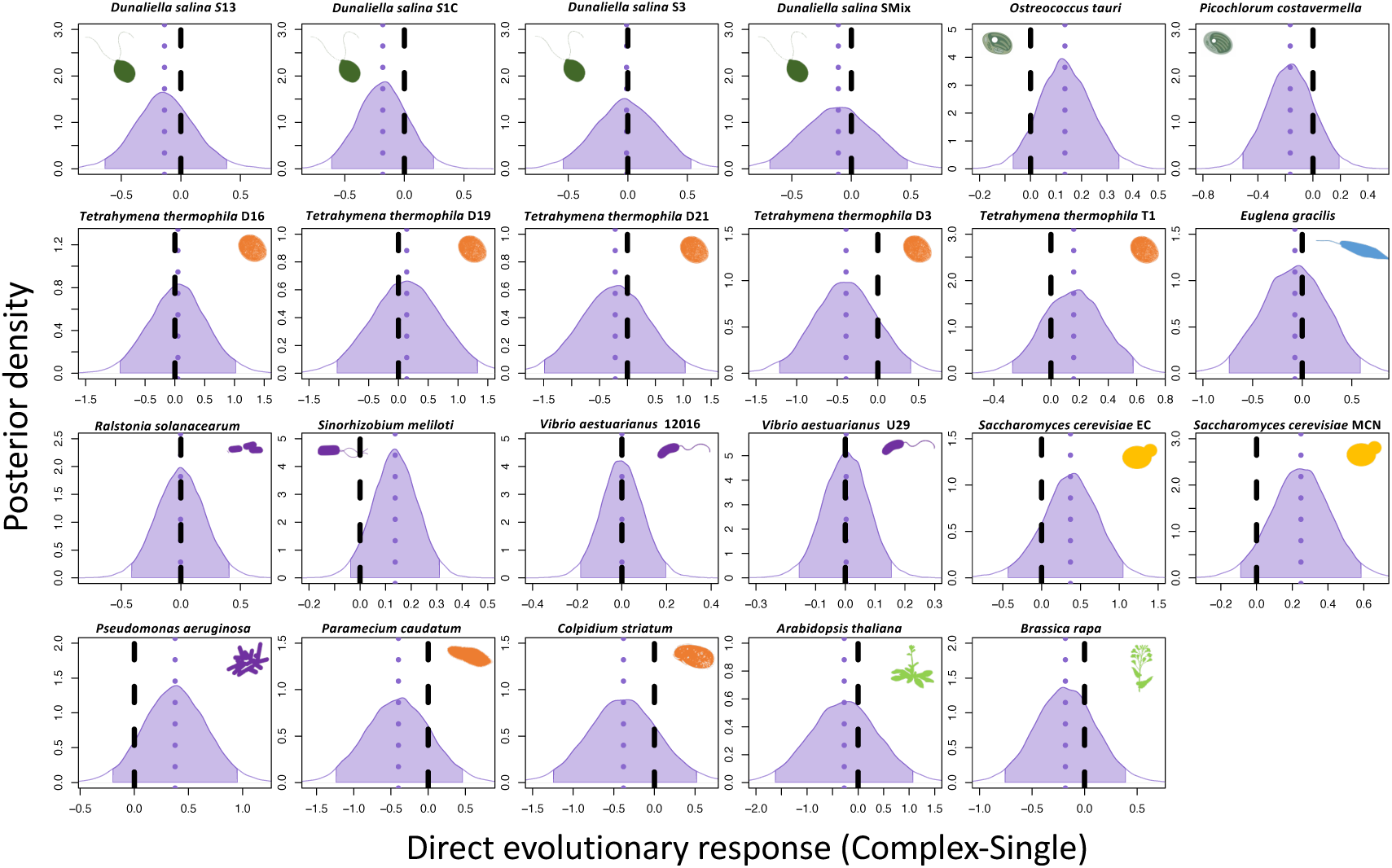
Posterior distributions, for each of the 14 species, of the difference between combined minus single stressors (contrast) of the direct response. The dashed purple lines visualize the median posterior values, and the shaded area the 95% compatibility intervals of the posterior distributions. Different strains of the same species are indicated next to their scientific name, with their silhouette repeated. Overlap of the distributions with the reference (black dashed line at zero values) indicates comparable direct response for simple vs. complex stress treatments.

**Fig. S5.**
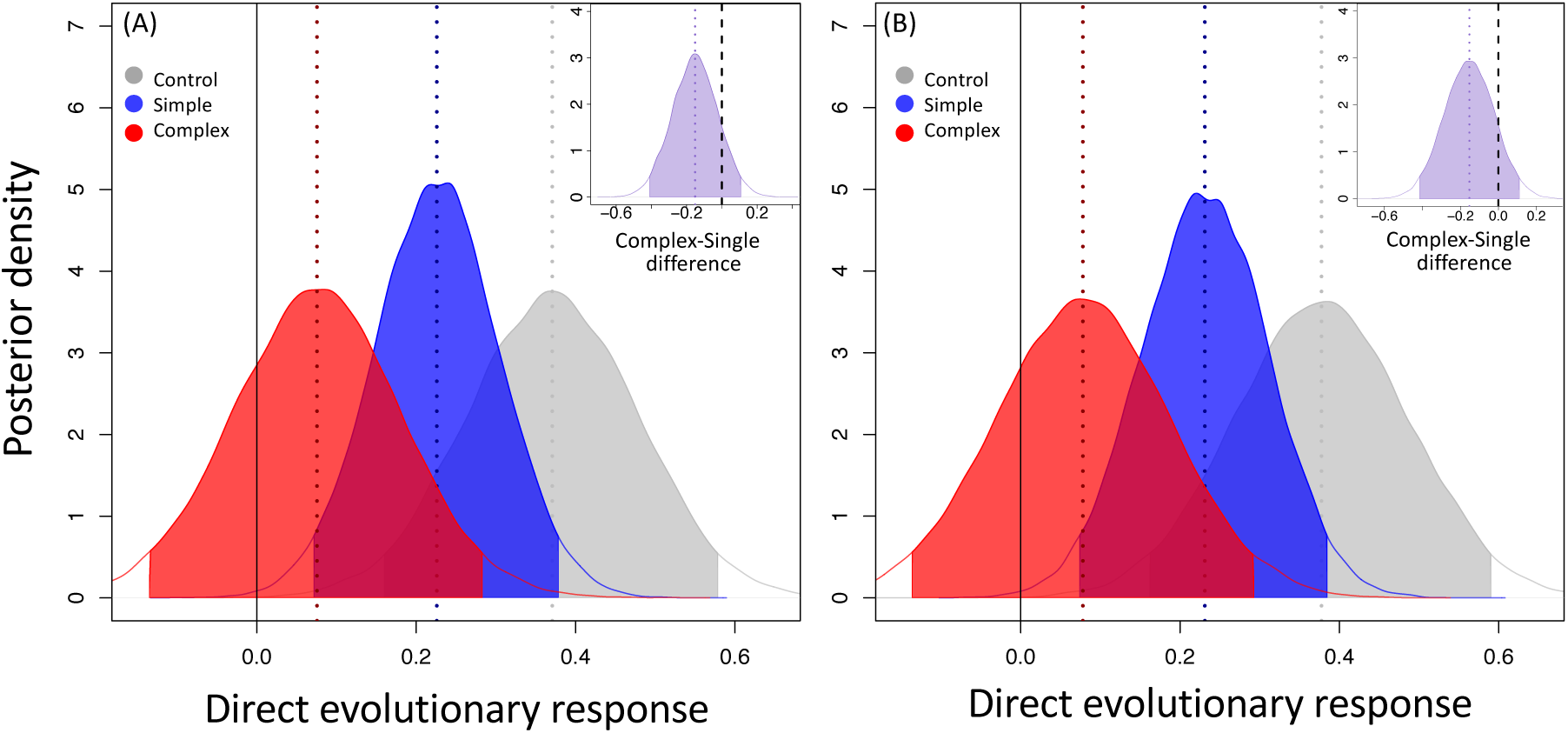
Posterior distributions for direct evolutionary responses without the two plant species (A) and without the two plant species and a *D. salina* strain (B), which started the experimental evolution from some standing genetic variation. Fitness estimates were obtained from assay in own evolutionary conditions. The posteriors were obtained by calculating the difference between final and initial fitness for the control, single and combined stressors evolutionary treatments respectively (A: 12 species and 21 strains, B: 11 species and 20 strains). The inset illustrates the posterior difference combined and single stressor.

**Fig. S6.**
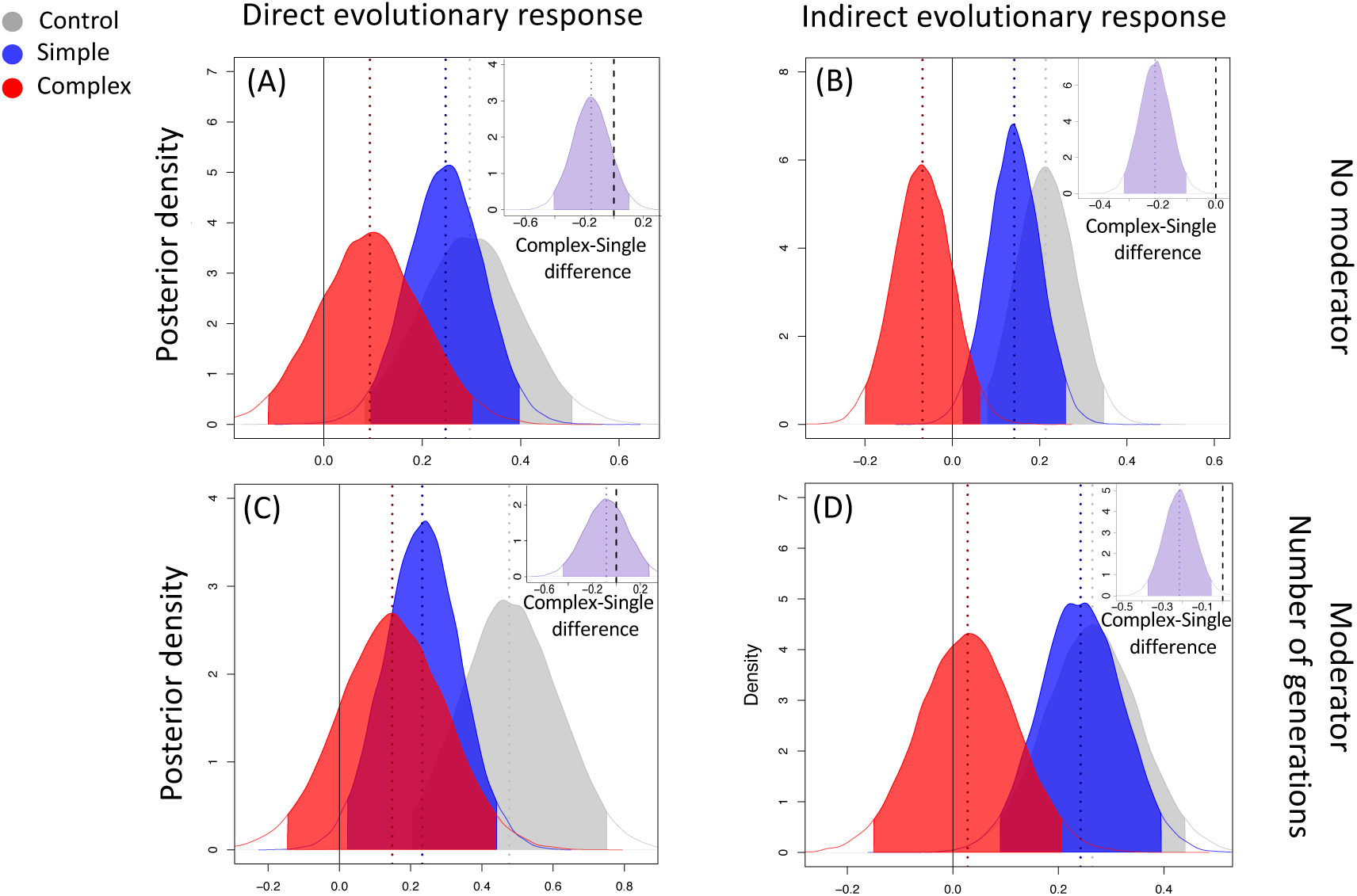
Posterior distributions for direct (14 species and 23 strains) and indirect (11 species and 20 strains) evolutionary responses with (A-B) or without (C-D) number of generations as moderator. The inset illustrates the posterior difference combined and single stressor. Panels (A-B) correspond to the one reported in the main text.

**Fig. S7.**
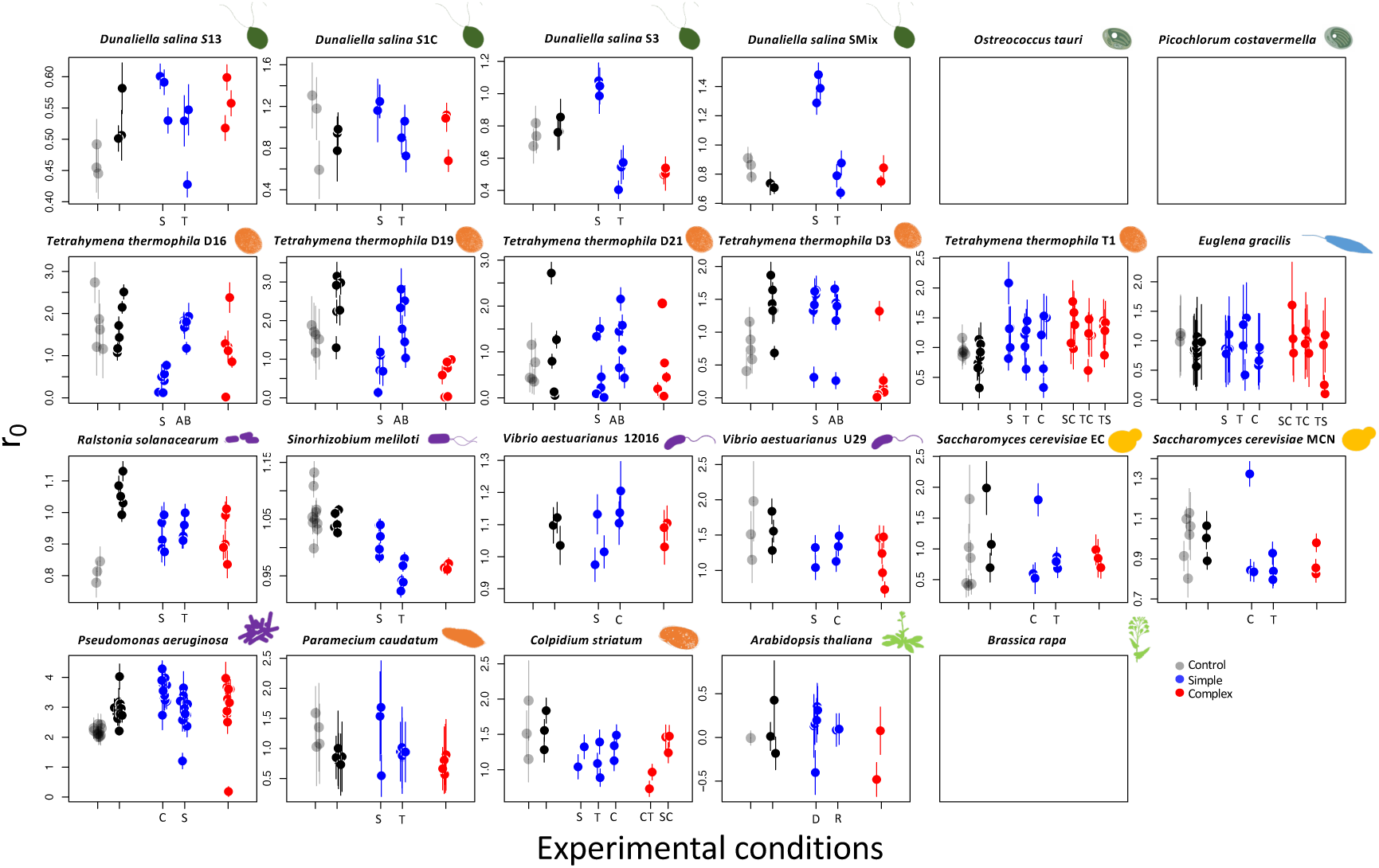
Fitness estimates (*r_0_*) of evolved lines assayed in control conditions for 11/14 species (silhouette), the 3 empty plots correspond to species for which such back-to-control conditions were not tested. The evolutionary treatment of origin is indicated by different colours; grey is control, blue is single stressors, and red is combined stressors. Shaded grey points are the initial fitness estimates in control conditions, while thick points are the final fitness estimates at the end of the evolutionary experiments. Different strains of the same species are indicated next to their scientific name, with their silhouette repeated. Points and bars represent the fitness estimate and standard deviation for a biological replicate. Capital letters indicate the identity of the single stressors experienced during evolution, S (Salt), T (Temperature), C (Copper), AB (Antibiotics), D (Disturbance), R (Resources), or of the combined stressors when more than one combination was used (i.e., ST, TC, TS).

**Fig. S8.**
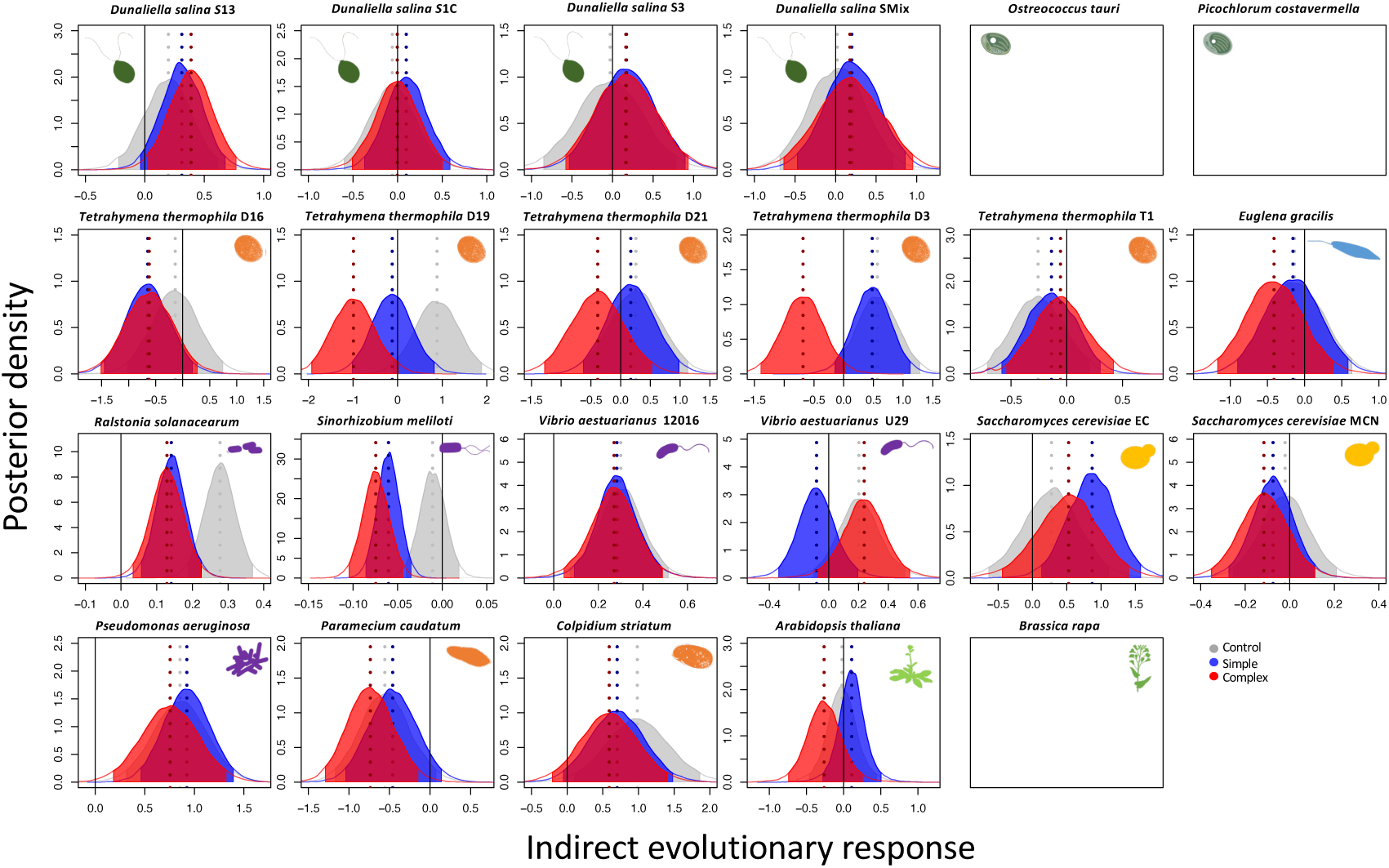
Posterior distributions for indirect evolutionary responses of each strain. Positive values represent a gain in fitness over the evolutionary experiment. In grey is the control in the absence of stressor, in blue and red the single and combined stressors respectively. The dashed lines visualize the median posterior values, and the shaded area the 95% compatibility intervals. The posteriors were obtained by calculating the difference between the final fitness of the 3 evolutionary treatments assayed in control conditions and the initial fitness in control conditions (11 species and 20 strains, reduced dataset).

**Fig. S9.**
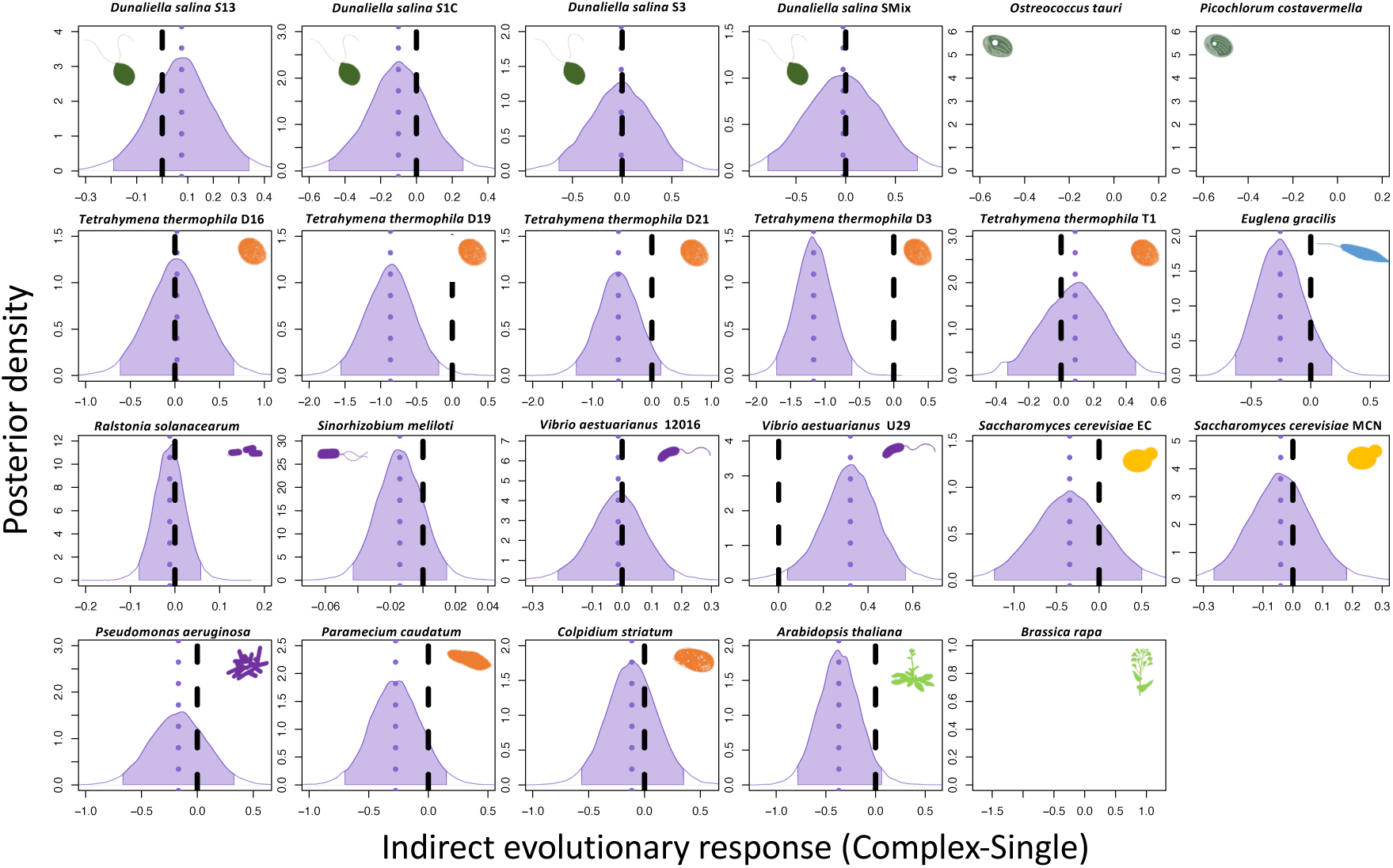
Posterior distributions, for 11 species and 20 strains (reduced dataset), of the difference between combined minus single stressors (contrast) of the indirect response. The dashed purple lines visualize the median posterior values, and the shaded area the 95% compatibility intervals of the posterior distributions. Different strains of the same species are indicated next to their scientific name, with their silhouette repeated. Overlap of the distributions with the reference (black dashed line at zero values) indicates comparable indirect response for simple vs. complex stress treatments.

**Fig. S10.**
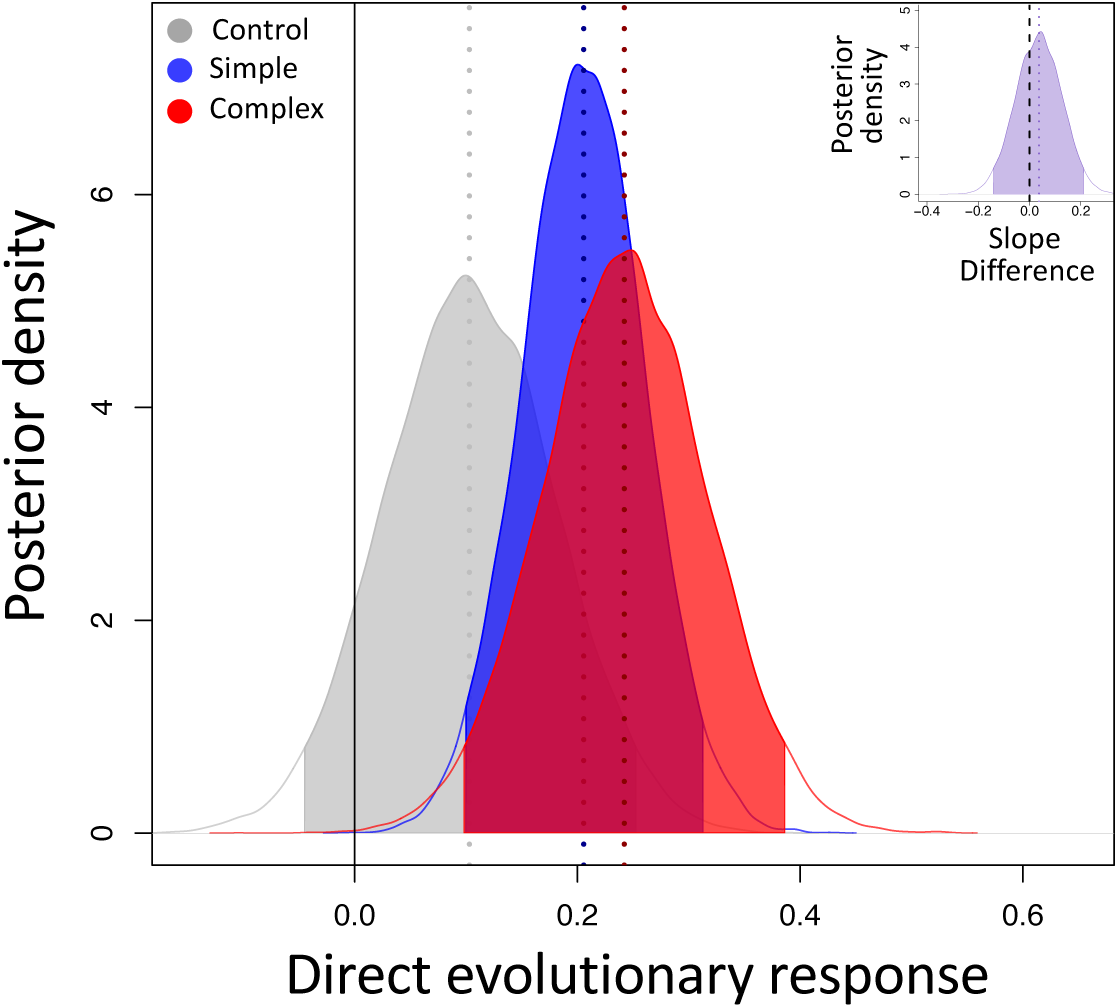
Posterior distributions for direct evolutionary responses without the four *Tetrahymena spp.* strains that evolved under antibiotics. Fitness estimates were obtained from assay in own evolutionary conditions. The posteriors were obtained by calculating the difference between final and initial fitness for the control, single and combined stressors evolutionary treatments respectively. The inset illustrates the posterior difference combined and single stressor.

**Fig. S11.**
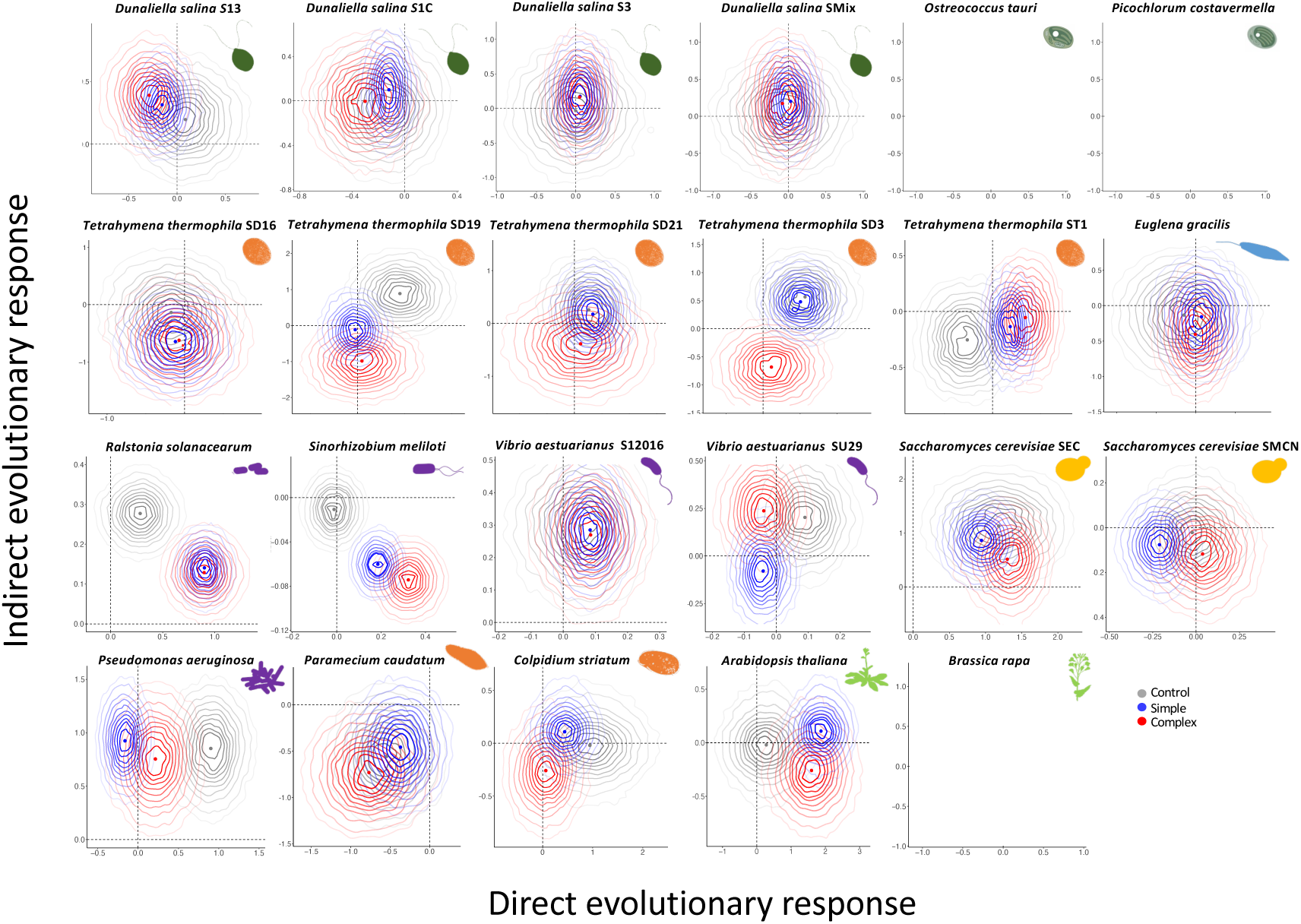
Density distributions of the direct response to selection (assay in own evolutionary conditions) against the indirect evolutionary response (assay in control conditions) for the control, single and combined stressors (11 species and 20 strains). Closed lines are the probability regions spanning from 10% to 95% from darker to lighter. The points and dashed lines are the median values of the posterior distributions. The direct and indirect responses correspond to the difference between final and initial fitness, with positive values representing a gain in fitness over the evolutionary experiment.

**Fig. S12.**
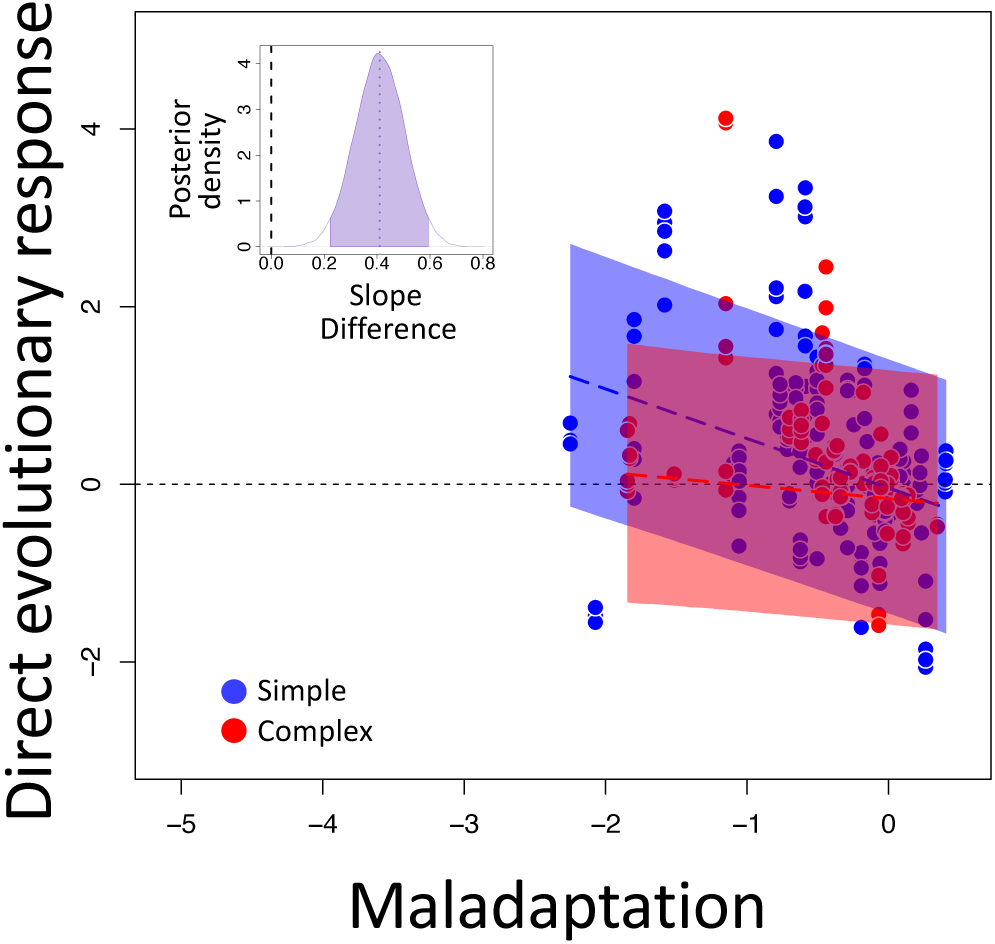
Declining adaptability in simple vs. complex environments. Relationship between direct response to selection and initial maladaptation (fitness in treatment minus fitness in control), for single (blue) and combined (red) stressors. The extreme values of maladaptation (<-3) were removed. Each point represents an independent evolutionary replicate, standardized as the difference to its initial average control (14 species and 23 strains, further details in the Material and Methods). The dashed lines are the median, and the shaded areas the 95% CI of the model predictions. The inset represents the posterior distribution of the difference complex minus simple, for the slope of the relationship between direct response to selection and initial maladaptation.

**Fig. S13.**
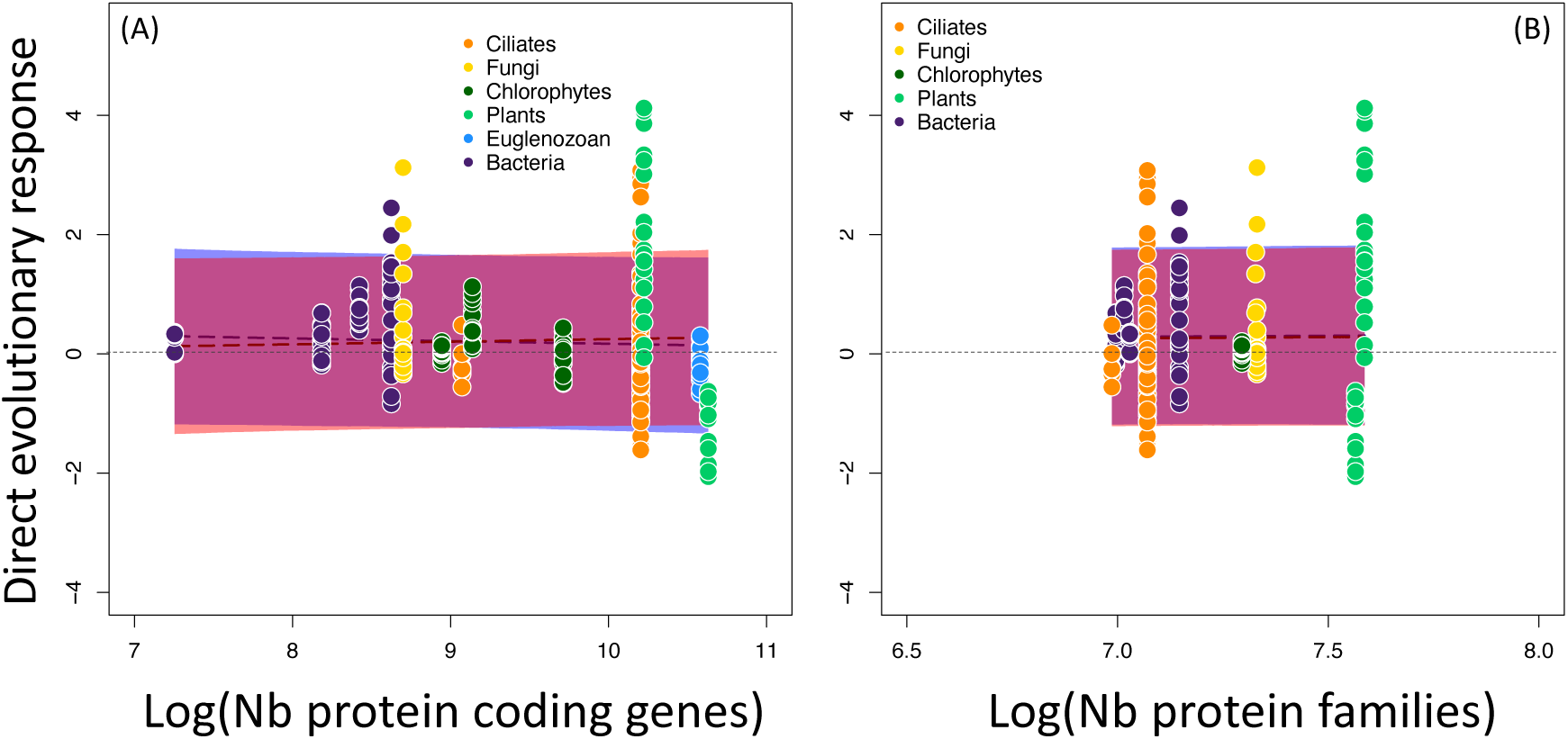
Direct evolutionary response in fitness as a function of proxies of organismal complexity. (A) Change in fitness of the taxon in function of their complexity defined as the logarithm of the number of protein coding genes. Information for Colpidium striatum was not available, and it was removed from this analysis (13 species and 22 strains). (B) Change in fitness of the taxon in function of their complexity (logarithm of the number of protein families). The points are the mean fitness value of an independent evolutionary replicate relative to its average control, and each colour identifies a different taxon. Dashed lines are the median, and the shaded areas the 95% CI of the model predictions for single (blue) and combined (red) stressors.

**Fig. S14.**
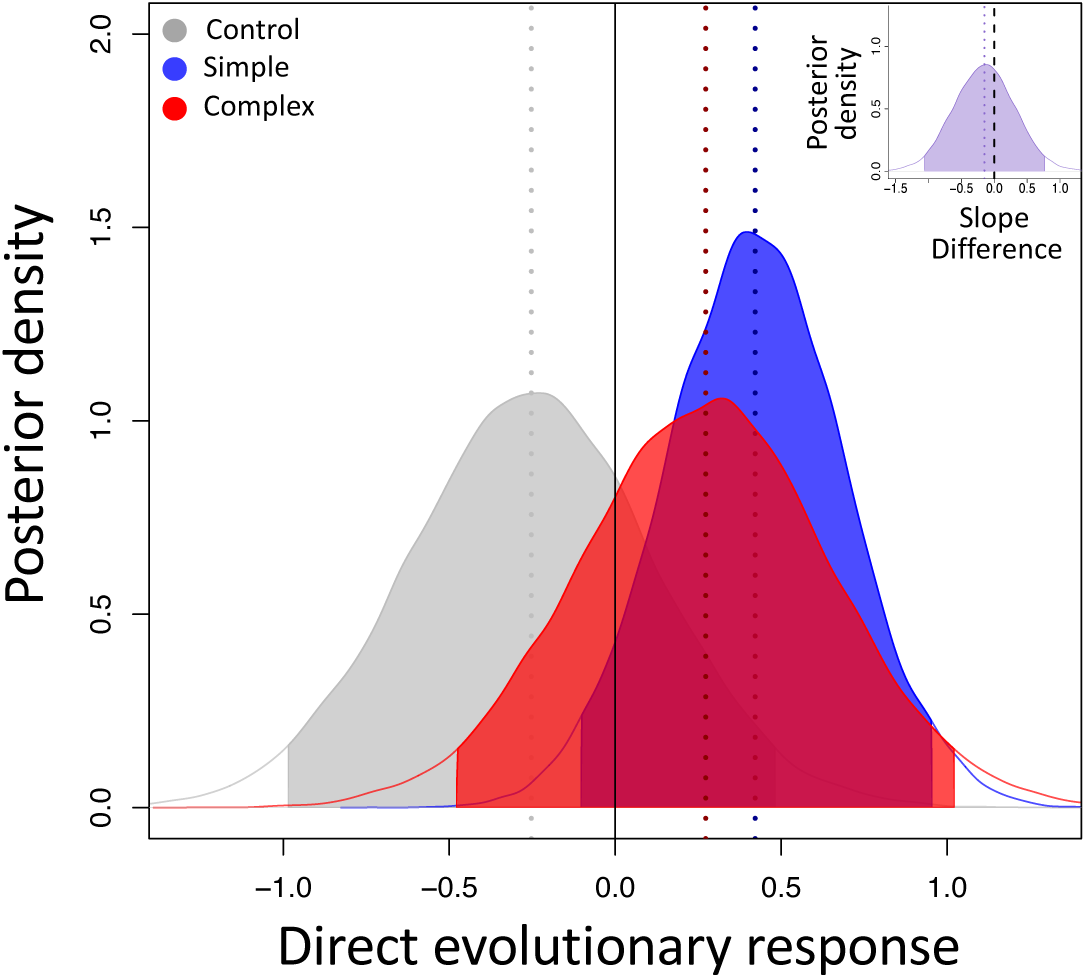
Posterior distributions for direct evolutionary responses for two plant species and one experimental group of the chlorophyte *D. salina* (standing genetic variation). Fitness estimates were obtained from assay in own evolutionary conditions. The posteriors were obtained by calculating the difference between final and initial fitness for the control, single and combined stressors evolutionary treatments respectively. The inset illustrates the posterior difference combined and single stressor.

**Fig. S15.**
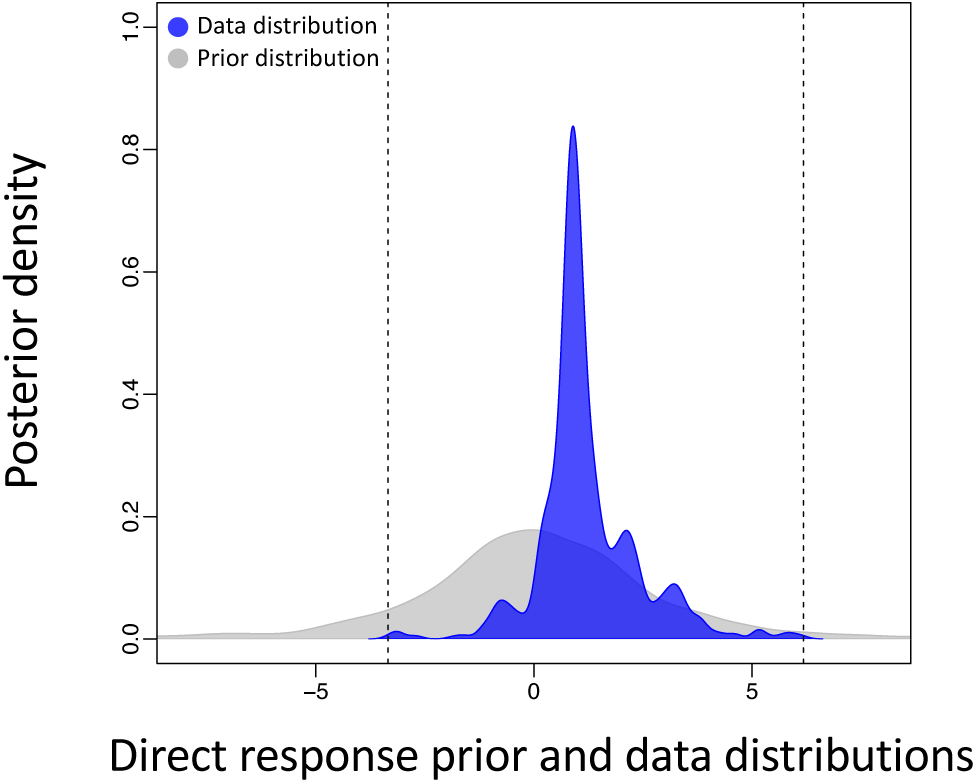
Prior predictive check for the direct response model. In grey the prior distribution and in blue the distribution of the fitness data. The dashed lines are the range or the data.

**Fig. S16.**
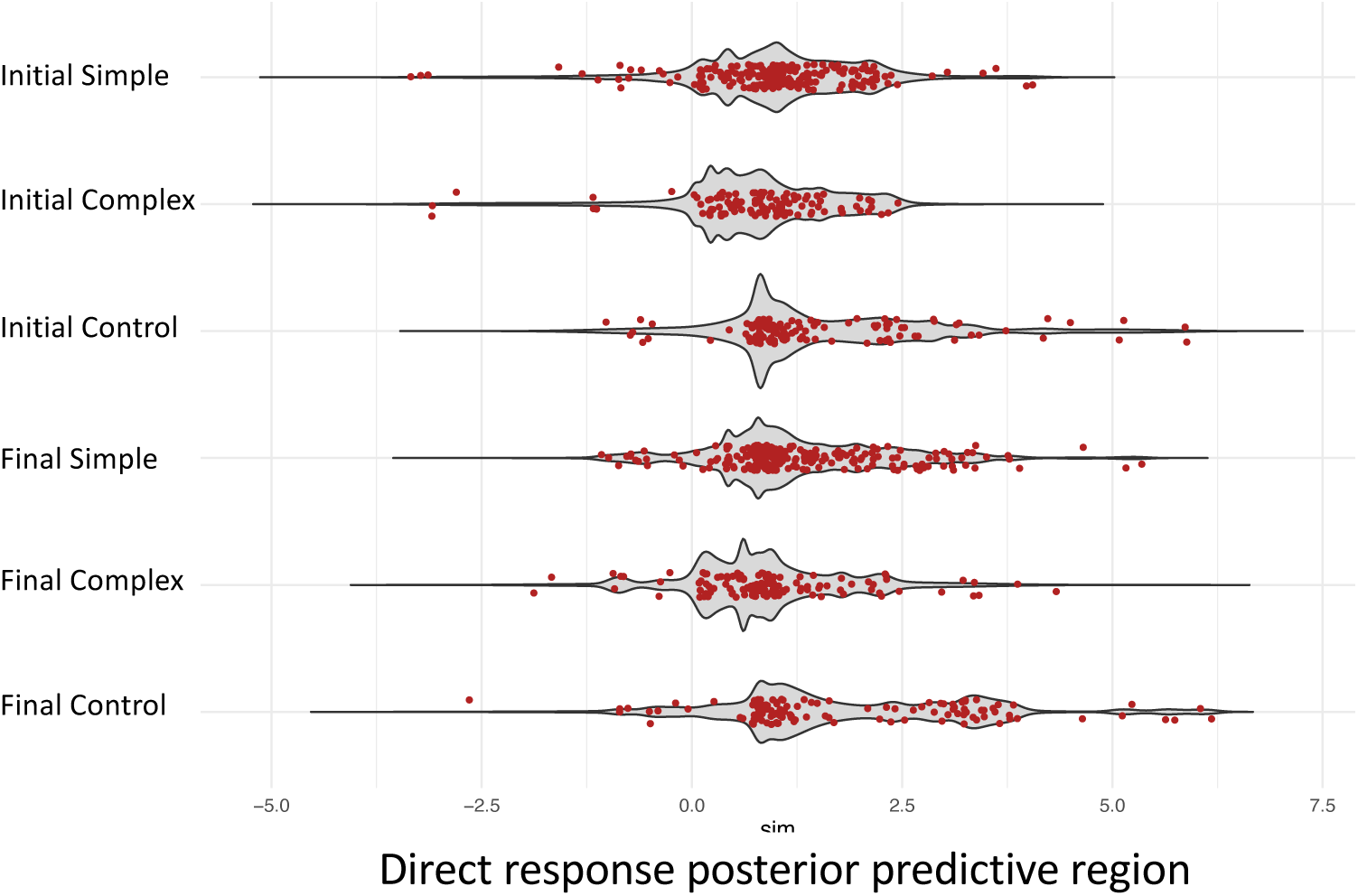
Posterior predictive distribution in grey for the direct response model including the error propagation, each red dot is an observed fitness value. The slightly composite (rather than completely smooth) predictive region in grey reflects the pooling of specific measurement-error distributions.

## Tables

**Table S1.** Priors and model details used to analyse the direct response of each strain independently (individual analysis). Each row is a different model.

| **Species** | **Intercept** | **# Iterations of sampling** | **Warm-up period** | **Acceptance probability (adapt_delta)** |
|---|---|---|---|---|
| *Arabidopsis thaliana* | normal(0,1) | 30000 | 1000 | 0.999 |
| *Brassica rapa* | normal(0,1) | 30000 | 10000 | 0.99999999 |
| *Colpidium striatum* | normal(0,1) | 30000 | 1000 | 0.999 |
| *Dunaliella salina*<br>(all strains) | normal(0,1) | 30000 | 10000 | 0.99999999 |
| *Euglena gracilis* | normal(0,1) | 40000 | 10000 | 0.999999999999 |
| *Ostreococcus tauri* | normal(0,1) | 30000 | 10000 | 0.99999999 |
| *Paramecium caudatum* | normal(0,1) | 30000 | 1000 | 0.999 |
| *Picochlorum costavermella* | normal(0,1) | 30000 | 10000 | 0.99999999 |
| *Pseudomonas aeruginosa* | normal(0,1) | 30000 | 1000 | 0.999 |
| *Ralstonia solanacearum* | normal(0,1) | 40000 | 10000 | 0.999999999999 |
| *Saccharomyces cerevisiae*<br>(all strains) | normal(0,1) | 30000 | 1000 | 0.999 |
| *Sinorhizobium meliloti* | normal(0,1) | 40000 | 10000 | 0.999999999999 |
| *Tetrahymena thermophila*<br>(D16, D19, D21, D3)<br>(T1) | normal(0,1)<br>normal(0,1) | 30000<br>40000 | 10000<br>10000 | 0.99999999<br>0.999999999999 |
| *Vibrio aestuarianus* | normal(0,1) | 30000 | 10000 | 0.999 |

**Table S2.**
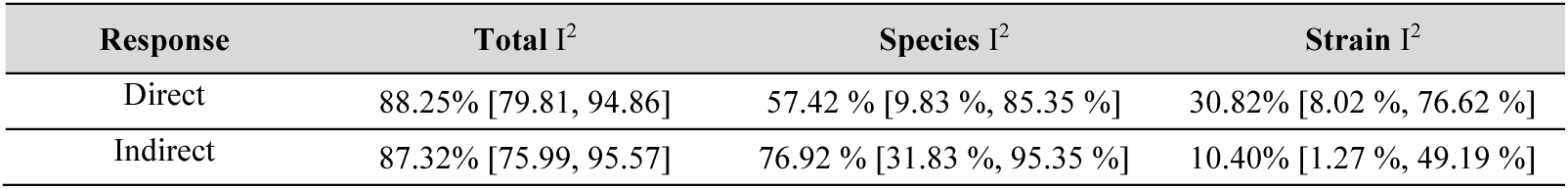
Heterogeneity (I^2^) and 95% CI for direct and indirect response.

| **Response** | **Total** I<sup>2</sup> | **Species** I<sup>2</sup> | **Strain** I<sup>2</sup> |
|---|---|---|---|
| Direct | 88.25% [79.81, 94.86] | 57.42 % [9.83 %, 85.35 %] | 30.82% [8.02 %, 76.62 %] |
| Indirect | 87.32% [75.99, 95.57] | 76.92 % [31.83 %, 95.35 %] | 10.40% [1.27 %, 49.19 %] |

**Table S3.** Priors and model details used to analyse the indirect response of each strain independently (individual analysis). Each row is a different model.

| **Species** | **Intercept** | **# Iterations of sampling** | **Warm-up period** | **Acceptance probability (adapt_delta)** |
|---|---|---|---|---|
| *Arabidopsis thaliana* | normal(0,1) | 30000 | 1000 | 0.999 |
| *Colpidium striatum* | normal(0,1) | 30000 | 1000 | 0.999 |
| *Dunaliella salina*<br>(all strains) | normal(0,1) | 30000 | 10000 | 0.99999999 |
| *Euglena gracilis* | normal(0,1) | 40000 | 10000 | 0.999999999999 |
| *Paramecium caudatum* | normal(0,1) | 30000 | 1000 | 0.999 |
| *Pseudomonas aeruginosa* | normal(0,1) | 30000 | 1000 | 0.999 |
| *Ralstonia solanacearum* | normal(0,1) | 40000 | 10000 | 0.999999999999 |
| *Saccharomyces cerevisiae*<br>(all strains) | normal(0,1) | 30000 | 1000 | 0.999 |
| *Sinorhizobium meliloti* | normal(0,1) | 40000 | 10000 | 0.999999999999 |
| *Tetrahymena thermophila* |  |  |  |  |
| (strain D16, D19, D21, D3) | normal(0,1) | 30000 | 10000 | 0.99999999 |
| (strain T1) | normal(0,1) | 40000 | 10000 | 0.999999999999 |
| *Vibrio aestuarianus* | normal(0,1) | 30000 | 1000 | 0.999 |

**Table S4.** Posterior distributions of the direct and indirect response for the overall stressors or each stressor.

| **Stressors** | **Direct response<br>(mean ± 95% CI)** |  | **Indirect response<br>(mean ± 95% CI)** |  |
|---|---|---|---|---|
|  | **Simple** | **Complex** | **Simple** | **Complex** |
| Overall | 0.247<br>[0.095, 0.397] | 0.093<br>[-0.113, 0.301] | 0.141<br>[0.023,0.258] | -0.068<br>[-0.200, 0.062] |
| Antibiotics | 3.427<br>[2.855, 4.002] | 1.632<br>[1.138, 2.119] | 0.380<br>[-0.010, 0.768] | -0.541<br>[-0.919, -0.160] |
| Temperature | 0.152<br>[-0.003, 0.305] | 0.168<br>[0.031, 0.302] | -0.051<br>[-0.102, -0.0001] | -0.033<br>[-0.082, 0.018] |
| Salt | -0.005<br>[-0.141, 0.127] | 0.081<br>[-0.048, 0.209] | 0.039<br>[-0.137, 0.217] | -0.107<br>[-0.278, 0.063] |
| Copper | 0.080<br>[-0.090, 0.259] | 0.171<br>[0.012, 0.333] | 0.210<br>[0.048, 0.383] | 0.154<br>[-0.004, 0.328] |
| Herbivory | 1.798<br>[0.579, 2.949] | 1.559<br>[0.343, 2.729] | 0.113<br>[-0.290, 0.505] | -0.257<br>[-0.742, 0.266] |
| Resources | 1.609<br>[0.464, 2.725] | 1.599<br>[0.456, 2.690] | 0.113<br>[-0.290, 0.505] | -0.257<br>[-0.742, 0.266] |

**Table S5.** Model comparison for a linear or quadratic relationship between direct response and initial maladaptation. The best model is highlighted in bold.

| **Model** | **# Iterations of sampling** | **Warm-up period** | **Acceptance probability (adapt_delta)** | **WAIC** | **Δ WAIC** | **SE** | **WAIC weight** |
|---|---|---|---|---|---|---|---|
| **Linear** | **35000** | **10000** | **0.99999999999999** | **23715843600** |  | **7032920346** | **1** |
| Quadratic | 35000 | 10000 | 0.99999999999999 | 24487085616 | 771242017 | 7264572800 | 0 |

**Table S6.** Priors and model details used to obtain the fitness estimates of direct response (14 species and 23 strains). Each row is a different model, and (-) indicates that parameters were not estimated for that specific model.

| **Species** | **Intercept** | **Slope** | **Technical replicates** | **# Iterations of sampling** | **Warm-up period** | **Acceptance probability (adapt_delta)** |
|---|---|---|---|---|---|---|
| *Arabidopsis thaliana* | normal(0,1) | - | - | 30000 | 15000 | 0.999 |
| *Brassica rapa* | normal(0,1) | - | normal(0,1) | 30000 | 15000 | 0.999 |
| *Colpidium striatum* | normal(0,1) | halfnormal(0,1) | normal(0,1) | 30000 | 15000 | 0.999 |
| *Dunaliella salina*<br>(all strains) | normal(0,1) | halfnormal(0,1) | - | 30000 | 15000 | 0.999 |
| *Euglena gracilis* | normal(0,1) | halfnormal(0,1) | - | 30000 | 15000 | 0.999 |
| *Ostreococcus tauri* | normal(0,1) | halfnormal(0,1) | - | 30000 | 15000 | 0.999 |
| *Paramecium caudatum* | normal(0,1) | halfnormal(0,1) | - | 30000 | 15000 | 0.999 |
| *Picochlorum costavermella* | normal(0,1) | halfnormal(0,1) | - | 30000 | 15000 | 0.999 |
| *Pseudomonas aeruginosa* | normal(0,1) | halfnormal(0,1) | - | 30000 | 15000 | 0.999 |
| *Ralstonia solanacearum* | normal(0,1) | halfnormal(0,1) | normal(0,1) | 30000 | 15000 | 0.9999999 |
| *Saccharomyces cerevisiae*<br>(all strains) | normal(0,1) | halfnormal(0,1) | normal(0,1) | 30000 | 15000 | 0.9999999 |
| *Sinorhizobium meliloti* | normal(0,1) | halfnormal(0,1) | normal(0,1) | 30000 | 15000 | 0.9999999 |
| *Tetrahymena thermophila*<br>(all strains) | normal(0,1) | halfnormal(0,1) | normal(0,1) | 30000 | 15000 | 0.999 |
| *Vibrio aestuarianus* | normal(0,1) | halfnormal(0,1) | - | 30000 | 15000 | 0.999 |

**Table S7.** Priors and model details used to obtain the fitness estimates of indirect response (11 species and 20 strains). Each row is a different model, and (-) indicates that parameters were not estimated for that specific model.

| **Species** | **Intercept** | **Slope** | **Technical replicates** | **# Iterations of sampling** | **Warm-up period** | **Acceptance probability (adapt_delta)** |
|---|---|---|---|---|---|---|
| *Arabidopsis thaliana* | normal(0,1) | - | - | 30000 | 15000 | 0.999999999 |
| *Colpidium striatum* | normal(0,1) | halfnormal(0,1) | normal(0,1) | 30000 | 15000 | 0.999 |
| *Dunaliella salina*<br>(all strains) | normal(0,1) | halfnormal(0,1) | - | 30000 | 15000 | 0.999 |
| *Euglena gracilis* | normal(0,1) | halfnormal(0,1) | - | 30000 | 15000 | 0.999 |
| *Paramecium caudatum* | normal(0,1) | halfnormal(0,1) | - | 30000 | 15000 | 0.999 |
| *Pseudomonas aeruginosa* | normal(0,1) | halfnormal(0,1) | - | 30000 | 15000 | 0.999 |
| *Ralstonia solanacearum* | normal(0,1) | halfnormal(0,1) | normal(0,1) | 30000 | 15000 | 0.9999999 |
| *Saccharomyces cerevisiae*<br>(all strains) | normal(0,1) | halfnormal(0,1) | normal(0,1) | 30000 | 15000 | 0.9999999 |
| *Sinorhizobium meliloti* | normal(0,1) | halfnormal(0,1) | normal(0,1) | 30000 | 15000 | 0.9999999 |
| *Tetrahymena thermophila*<br>(all strains) | normal(0,1) | halfnormal(0,1) | normal(0,1) | 30000 | 15000 | 0.999 |
| *Vibrio aestuarianus* | normal(0,1) | halfnormal(0,1) | - | 30000 | 15000 | 0.999 |

**Table S8.** Model comparison for including or not the phylogeny in the main analysis, here reported for the direct response. The best model is highlighted in bold.

| **Model** | **df** | **logLik** | **AICc** | **Δ AICc** | **LRT** | **p** |
|---|---|---|---|---|---|---|
| **No phylogeny** | **8** | **-2566.44** | **5149.05** | **0.40** | **1.64** | **0.201** |
| Phylogeny | 9 | -2565.62 | 5149.45 | - | - | - |

**Table S9.** Priors and model details use to analyse direct and indirect response for the overall stressors and each stressor. Each row is a different model, and (-) indicates that parameters were not estimated for that specific model.

| **Main analysis** | **Stressors** | **# Species** | **# Strains** | **Intercept** | **Slope** | **Strain** | **Species** | **# Iterations of sampling** | **Warm-up period** | **Acceptance probability (adapt_delta)** |
|---|---|---|---|---|---|---|---|---|---|---|
| Direct response | Overall | 14 | 23 | normal(0,1) | - | normal(0,1) | normal(0,1) | 30000 | 1000 | 0.999 |
| Direct response (no plants) | Overall | 12 | 21 | normal(0,1) | - | normal(0,1) | normal(0,1) | 30000 | 1000 | 0.999 |
| Direct response (no plants and *D. salina*) | Overall | 11 | 21 | normal(0,1) | - | normal(0,1) | normal(0,1) | 30000 | 1000 | 0.999 |
| Direct response (#generations) | Overall | 14 | 21 | normal(0,1) | - | normal(0,1) | normal(0,1) | 30000 | 1000 | 0.999 |
| Direct response (no *T. thermophila*) | Overall (no antibiotics) | 14 | 19 | normal(0,1) | - | normal(0,1) | normal(0,1) | 30000 | 1000 | 0.999 |
| Direct response (merged data) | Overall | 11 | 20 | normal(0,1) | - | normal(0,1) | normal(0,1) | 30000 | 1000 | 0.999 |
| Indirect response | Overall | 11 | 20 | normal(0,1) | - | normal(0,1) | normal(0,1) | 30000 | 1000 | 0.999 |
| Indirect response (#generations) | Overall | 11 | 20 | normal(0,1) | - | normal(0,1) | normal(0,1) | 30000 | 1000 | 0.999 |
| Indirect response (merged data) | Overall | 11 | 20 | normal(0,1) | - | normal(0,1) | normal(0,1) | 30000 | 1000 | 0.999 |
| Direct response (maladaptation) | Overall | 14 | 23 | normal(0,1) | normal(0,1) | normal(0,1) | normal(0,1) | 10000 | 2000 | 0.99999999999999 |
| Direct response (#protein-coding genes) | Overall | 13 | 22 | normal(0,1) | normal(0,1) | normal(0,1) | normal(0,1) | 35000 | 10000 | 0.99999999999999 |
| Direct response (#protein families) | Overall | 13 | 22 | normal(0,1) | normal(0,1) | normal(0,1) | normal(0,1) | 35000 | 10000 | 0.99999999999999 |
| Direct response | Antibiotics | 1 | 4 | normal(0,1) | - | normal(0,1) | normal(0,1) | 30000 | 1000 | 0.999 |
| Indirect response | Antibiotics | 1 | 4 | normal(0,1) | - | normal(0,1) | normal(0,1) | 30000 | 1000 | 0.999 |
| Direct response | Temperature | 8 | 12 | normal(0,1) | - | normal(0,1) | normal(0,1) | 30000 | 1000 | 0.999 |
| Indirect response | Temperature | 8 | 12 | normal(0,1) | - | normal(0,1) | normal(0,1) | 30000 | 1000 | 0.999 |
| Direct response | Salt | 9 | 17 | normal(0,1) | - | normal(0,1) | normal(0,1) | 30000 | 1000 | 0.999 |
| Indirect response | Salt | 9 | 17 | normal(0,1) | - | normal(0,1) | normal(0,1) | 30000 | 1000 | 0.999 |
| Direct response | Copper | 6 | 8 | normal(0,1) | - | normal(0,1) | normal(0,1) | 30000 | 1000 | 0.999 |
| Indirect response | Copper | 6 | 8 | normal(0,1) | - | normal(0,1) | normal(0,1) | 30000 | 1000 | 0.999 |
| Direct response | Herbivory | 1 | 1 | normal(0,1) | - | normal(0,1) | normal(0,1) | 30000 | 1000 | 0.999 |
| Indirect response | Herbivory | 1 | 1 | normal(0,1) | - | normal(0,1) | normal(0,1) | 30000 | 1000 | 0.999 |
| Direct response | Resources | 1 | 1 | normal(0,1) | - | normal(0,1) | normal(0,1) | 30000 | 1000 | 0.999 |
| Indirect response | Resources | 1 | 1 | normal(0,1) | - | normal(0,1) | normal(0,1) | 30000 | 1000 | 0.999 |

**Table S10.** Evolutionary treatments and number of replicates at the start and end of the experiment for *Colpidium caudatum*.

| **Strain ID** | **Salinity (g/L)** | **Copper (µg/L)** | **Temperature (°C)** | **Stress** | **Replicates start/end** |
|---|---|---|---|---|---|
| *C. striatum* | 0 | 0 | 20 | Control | 3/3 |
| *C. striatum* | 0 | 0 | 24 | Single Temperature | 3/3 |
| *C. striatum* | 3 | 0 | 20 | Single Salinity | 3/2 |
| *C. striatum* | 0 | 200 | 20 | Single Copper | 3/3 |
| *C. striatum* | 3 | 0 | 24 | Combined Salinity x Temperature | 3/0 |
| *C. striatum* | 3 | 200 | 20 | Combined Salinity x Copper | 3/2 |
| *C. striatum* | 0 | 200 | 24 | Combined Copper x Temperature | 3/3 |

**Table S11.** Overview of selection cultures and common garden environment condition used in the common garden experiment for *Colpidium caudatum*. S, C and T stand for salinity, copper, and temperature, respectively.

| **Selection environment** | **Assay environment (common garden environment)** |  |  |  |  |  |
|---|---|---|---|---|---|---|
|  | Control | Single salinity | Single copper | Single temperature | Combined S x C | Combined T x C |
| Control | x | x | x | x | x | x |
| Single salinity | x | x |  |  |  |  |
| Single copper | x |  | x |  |  |  |
| Single temperature | x |  |  | x |  |  |
| Combined S x C | x |  |  |  | x |  |
| Combined T x C | x |  |  |  |  | x |

**Table S12.** Overview of salt and copper concentrations for the common garden experiment of *Colpidium caudatum*. S, C and T stand for salinity, copper, and temperature, respectively.

| **Selection environment** | **Common garden environment** | **Total [L]** | **Culture [L]** | **Medium [L]** | **S culture [g/L]** | **S medium [g/L]** | **C culture [µg/L]** | **C medium [µg/L]** | **S end [g/L]** | **C end [µg/L]** |
|---|---|---|---|---|---|---|---|---|---|---|
| Control | Control / Single temperature | 0.1 | 0.003 | 0.097 | 0 | 0.093 | 0 | 6.186 | 0.093 | 6.186 |
|  | Single salinity | 0.1 | 0.003 | 0.097 | 0 | 3.093 | 0 | 6.186 | 3 | 6.186 |
|  | Single copper / Combined T x C | 0.1 | 0.003 | 0.097 | 0 | 0.093 | 0 | 206.186 | 0.093 | 200 |
|  | Combined S x C | 0.1 | 0.003 | 0.097 | 0 | 3.093 | 0 | 206.186 | 3 | 200 |
| Single salinity | Control | 0.1 | 0.003 | 0.097 | 3 | 0 | 0 | 6.186 | 0.093 | 6.186 |
|  | Single Salinity | 0.1 | 0.003 | 0.097 | 3 | 3 | 0 | 6.186 | 3 | 6.186 |
| Single copper | Control | 0.1 | 0.003 | 0.097 | 0 | 0.093 | 200 | 0 | 0.093 | 6.186 |
|  | Single copper | 0.1 | 0.003 | 0.097 | 0 | 0.093 | 200 | 200 | 0.093 | 200 |
| Single temperature | Control | 0.1 | 0.003 | 0.097 | 0 | 0.093 | 0 | 6.186 | 0.093 | 6.186 |
|  | Single temperature | 0.1 | 0.003 | 0.097 | 0 | 0.093 | 0 | 6.186 | 0.093 | 6.186 |
| Combined S x C | Control | 0.1 | 0.003 | 0.097 | 3 | 0 | 200 | 0 | 0.093 | 6.186 |
|  | Combined S x C | 0.1 | 0.003 | 0.097 | 3 | 3 | 200 | 200 | 3 | 200 |
| Combined T x C | Control | 0.1 | 0.003 | 0.097 | 0 | 0.093 | 200 | 0 | 0.093 | 6.186 |
|  | Combined T x C | 0.1 | 0.003 | 0.097 | 0 | 0.093 | 200 | 200 | 0.093 | 200 |

**Table S13.** Coordinates and additional information on the sampling sites for *Dunaliella salina*.

| **Site** | **Strain ID** | **Saltern** | **Site name** | **Latitude** | **Longitude** | **Salinity (in g/l)** | **Salinity (in M)** | **Mean salinity (in g/l)** | **Mean salinity (in M)** |
|---|---|---|---|---|---|---|---|---|---|
| S1 | S1 | Gruissan | Tables Gruissan | 43.096087 | 3.086734 | 280 | 4.8 | 280.00 | 4.8 |
| S3 | S3 | Gruissan | Lac | 43.086220 | 3.091718 | 120 | 2.1 | 80.00 | 1.4 |
| S11 | S11 | La Palme | Nord-Ouest | 42.973766 | 3.028587 | 130 | 2.2 | 133.33 | 2.3 |
| S13 | S13 | La Palme | Reprise | 42.976025 | 3.032964 | 200 | 3.4 | 176.67 | 3.0 |

**Table S14.** Artificial seawater composition for *Dunaliella salina* (total of 1l of water). The two solutions SS1 and SS2 were prepared separately to avoid unwanted chemical reactions during autoclaving. 20 ml of Guillard′s (F/2) Marine Water Enrichment Solution were added before use.

| Solution | Compound | Added mass [g] of the compound |
|---|---|---|
| SS1 (0.9l) | NaCl (only for the hypersaline solution) | 280.512 |
|  | Na<sub>2</sub>SO<sub>4</sub> | 3.55 |
|  | KCl | 0.599 |
|  | NaHCO<sub>3</sub> | 0.174 |
|  | KBr | 0.0863 |
|  | H<sub>3</sub>BO<sub>3</sub> | 0.023 |
|  | NaF | 0.0028 |
| SS2 (0.1l) | MgCl, 6 H2O | 9.592 |
|  | CaCl<sub>2</sub>, 2 H2O | 1.344 |
|  | SrCl<sub>2</sub>, 6 H2O | 0.0218 |

**Table S15.** Evolutionary treatments and replicates at the start and end of the experiment for *Dunaliella salina*.

| **Strain ID** | **Salinity** | **Temperature** | **Stress** | **Replicates** |
|---|---|---|---|---|
| S1 | 2M | 24°C | Control | 3 |
| S1 | 4M | 24°C | Single Salinity | 3 |
| S1 | 2M | 36°C | Single Temperature | 3 |
| S1 | 3.5M | 36°C | Combined | 3 |
| S1 | 4M | 36°C | Combined | 3 |
| S3 | 2M | 24°C | Control | 3 |
| S3 | 4M | 24°C | Single Salinity | 3 |
| S3 | 2M | 36°C | Single Temperature | 3 |
| S3 | 3.5M | 36°C | Combined | 3 |
| S3 | 4M | 36°C | Combined | 3 |
| S11 | 2M | 24°C | Control | 3 |
| S11 | 4M | 24°C | Single Salinity | 3 |
| S11 | 2M | 36°C | Single Temperature | 3 |
| S11 | 3.5M | 36°C | Combined | 3 |
| S11 | 4M | 36°C | Combined | 3 |
| S13 | 2M | 24°C | Control | 3 |
| S13 | 4M | 24°C | Single Salinity | 3 |
| S13 | 2M | 36°C | Single Temperature | 3 |
| S13 | 3.5M | 36°C | Combined | 3 |
| S13 | 4M | 36°C | Combined | 3 |
| Mix | 2M | 24°C | Control | 3 |
| Mix | 4M | 24°C | Single Salinity | 3 |
| Mix | 2M | 36°C | Single Temperature | 3 |
| Mix | 3.5M | 36°C | Combined | 3 |
| Mix | 4M | 36°C | Combined | 3 |

**Table S16.** Dilution rates for the different strains of *Dunaliella salina* during the weekly transfers. Variations from the 2/15 dilution are highlighted in bold, and X indicates the strain that failed to grow and was lost.

| **Transfer week** | **S1** |  | **S3** |  | **S11** |  | **S13** |  | **Mix** |  |
|---|---|---|---|---|---|---|---|---|---|---|
|  | 24°C | 36°C | 24°C | 36°C | 24°C | 36°C | 24°C | 36°C | 24°C | 36°C |
| 1 | 2/15 | 2/15 | 2/15 | 2/15 | 2/15 | 2/15 | 2/15 | 2/15 | 2/15 | 2/15 |
| 2 | 2/15 | 2/15 | 2/15 | **5/15** | 2/15 | **5/15** | 2/15 | 2/15 | 2/15 | 2/15 |
| 3 | 2/15 | 2/15 | 2/15 | **4/15** | 2/15 | **4/15** | 2/15 | 2/15 | 2/15 | 2/15 |
| 4 | 2/15 | 2/15 | 2/15 | **4/15** | 2/15 | **4/15** | 2/15 | 2/15 | 2/15 | 2/15 |
| 5 | 2/15 | 2/15 | **4/15** | **4/15** | **X** | **X** | 2/15 | 2/15 | 2/15 | 2/15 |
| 6 | 2/15 | 2/15 | **4/15** | **4/15** | **X** | **X** | 2/15 | 2/15 | 2/15 | 2/15 |
| 7 | 2/15 | 2/15 | 2/15 | **4/15** | **X** | **X** | 2/15 | 2/15 | 2/15 | 2/15 |

**Table S17.** Experiment design of the assays. Stressful conditions are indicated with “+”, while the control conditions are represented by “-”. Tet = *Tetrahymena thermophila* strain T1, Eug = *Euglena gracilis*, ns = no stressor, cu = copper, na = salt, tp = temperature, cs = copper + salt, ct = copper + temperature, st = salt + temperature, cst = copper+salt+temperature, T = temperature, C = copper level, S = salinity.

| **Species** | **Treatment** | **Variables** |  |  | **Replicates** | **Stressors** |  |  |
|---|---|---|---|---|---|---|---|---|
|  |  | Temperature (°C) | CuSO<sub>4</sub> (mM) | NaCl (g/L) |  | T | C | S |
| Tet | ns | 25 | - | - | 5 | - | - | - |
| Tet | cu | 25 | 0.005 | - | 5 | - | + | - |
| Tet | na | 25 | - | 7.5 | 5 | - | - | + |
| Tet | cs | 25 | 0.005 | 3 | 5 | - | + | + |
| Tet | tp | 36 | - | - | 5 | + | - | - |
| Tet | ct | 36 | 0.005 | - | 5 | + | + | - |
| Tet | st | 36 | - | 7.5 | 5 | + | - | + |
| Tet | cst | 36 | 0.005 | 3 | 5 | + | + | + |
| Eug | ns | 25 | - | - | 5 | - | - | - |
| Eug | cu | 25 | 0.005 | - | 5 | - | + | - |
| Eug | na | 25 | - | 7.5 | 5 | - | - | + |
| Eug | cs | 25 | 0.005 | 4.5 | 5 | - | + | + |
| Eug | tp | 33 | - | - | 5 | + | - | - |
| Eug | ct | 33 | 0.005 | - | 5 | + | + | - |
| Eug | st | 33 | - | 7.5 | 5 | + | - | + |
| Eug | cst | 33 | 0.005 | 4.5 | 5 | + | + | + |

**Table S18.** Photoperiod, media and light intensity of the control conditions of *Ostreococcus tauri* and *Picochlorum costavermella*.

| **Species** | *Ostreococcus tauri* and *Picochlorum costavermella* |
|---|---|
| **Temperature** | 26°C |
| **Media** | L1 (NCMA, Bigelow Lab for Ocean Science, USA) |
| **Hours Day:Night** | 8:16 |
| **Light Intensity** | 50 µmol.m<sup>-2</sup>.s<sup>-1</sup> |
| **Incubator** | Memmert IPP110 ECO PLUS |

**Table S19.** Conditions of the experimental evolution experiment for *Ostreococcus tauri* and *Picochlorum costavermella*.

| Species | Control |  | Temperature |  | Salt |  | Temperature + Salt |  | Experiment |
|---|---|---|---|---|---|---|---|---|---|
| *O. tauri* | 26°C | 0.60M | 32°C | 0.60M | 26°C | 0.86M | 32°C | 0.86M | Exp1 |
| *P. costavermella* | 26°C | 0.60M | 35°C | 0.60M | 26°C | 1.37M | 35°C | 1.37M | Exp1 |
| *P. costavermella* | 26°C | 0.60M | 35°C | 0.60M | 26°C | 1.03M | 35°C | 1.03M | Exp2 |

**Table S20.** Average growth rates and average growth rate decrease (in percent) in control *vs.* stressful experiments over 6 replicates for *Ostreococcus tauri* and *Picochlorum costavermella*.

| **Species** | **Control** | **Temperature (g \| %)** |  | **Salt (g \| %)** |  | **Temperature + Salt (g \| %)** |  | **Experiment** |
|---|---|---|---|---|---|---|---|---|
| *O. tauri* | 1.90 | 1.87 | -1.5% | 1.83 | -55% | 1.60 | -63% | Exp1 |
| *P. costavermella* | 1.83 | 1.54 | -16% | 1.74 | -57% | 1.05 | -95% | Exp1 |
| *P. costavermella* | 1.52 | 1.00 | -34% | 1.46 | -3.9% | 1.26 | -17% | Exp2 |

**Table S21.** Number of cycles per experiment for *Ostreococcus tauri* and *Picochlorum costavermella*.

| **Species** | **Control** | **Temperature** | **Salt** | **Temperature + Salt** | **Experiment** |
|---|---|---|---|---|---|
| *O. tauri* | 17 | 17 | 17 | 17 | Exp1 |
| *P. costavermella* | 17 | 17 | 17 | 0 | Exp1 |
| *P. costavermella* | 12 | 12 | 12 | 12 | Exp2 |

**Table S22.** Description of the 30-minute cycle used to follow the optical density of growing *Pseudomonas aeruginosa* populations.

| Length | Action | Description |
|---|---|---|
| 5s | Absorbance observation | Number of points measured: 5<br>OD 550nm |
| 22min | Waiting time | - |
| 5min | Shaking time | Shaking (Double Orbital) amplitude: 2, mm<br>Shaking frequency: 108rpm |
| 2min | Waiting time | - |

**Table S23.** Description of the 4 evolutionary treatments for *Pseudomonas aeruginosa*.

| Name | Treatment | Concentration |
|---|---|---|
| S0 | Control |  |
| S1 | Salt | 0.7M NaCl |
| S2 | Copper | 0,00325M CuSO<sub>4</sub><sup>-</sup> |
| S3 | Salt + Copper | 0.5M NaCl - 0,002M CuSO<sub>4-</sub> |

**Table S24.** List of the 20µL transfers during the experimental evolution of *Pseudomonas aeruginosa*.

| Cycle | Stress | Population |
|---|---|---|
| 5 | Copper | C11 |
| 10 | Copper | CN2 |
| 22 | Copper | C2 |
| 22 | Copper | C3 |
| 22 | Copper | C6 |
| 22 | Salt + Copper | CN10 |
| 26 | Salt + Copper | CN10 |

**Table S25.**
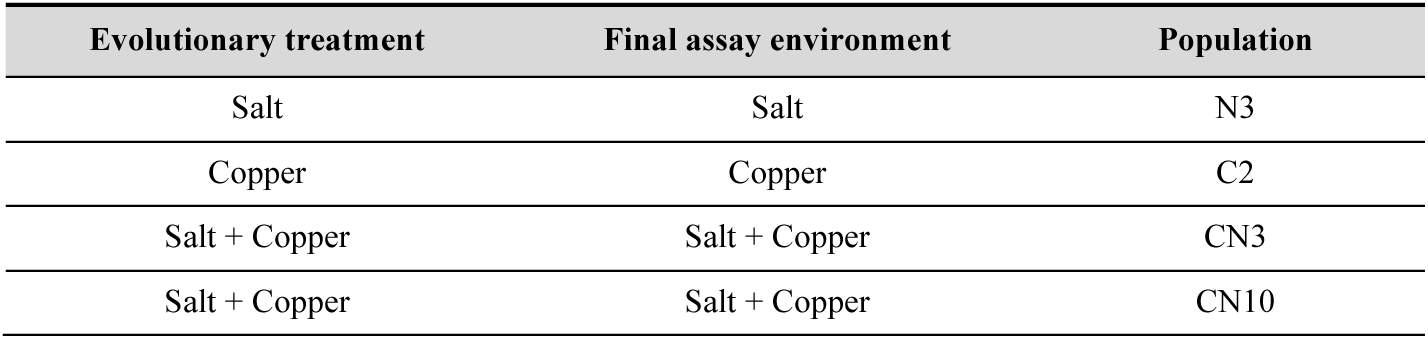
Populations of *Pseudomonas aeruginosa* that did not grow in the final phenotypic tests.

| **Evolutionary treatment** | **Final assay environment** | **Population** |
|---|---|---|
| Salt | Salt | N3 |
| Copper | Copper | C2 |
| Salt + Copper | Salt + Copper | CN3 |
| Salt + Copper | Salt + Copper | CN10 |

**Table S26.**
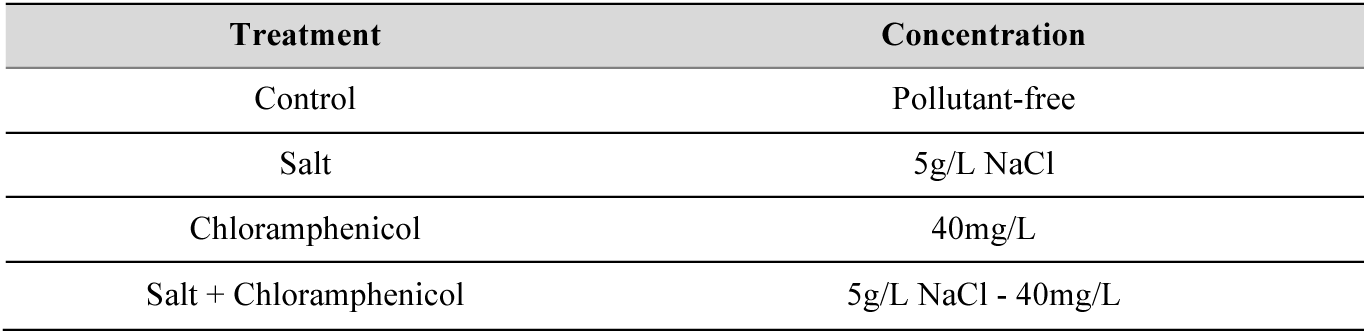
Description of the four evolutionary treatments for *Tetrahymena thermophila* (strains D16, D19, D21, D3). These four treatments were conducted in 0.3X PPYE medium.

| **Treatment** | **Concentration** |
|---|---|
| Control | Pollutant-free |
| Salt | 5g/L NaCl |
| Chloramphenicol | 40mg/L |
| Salt + Chloramphenicol | 5g/L NaCl - 40mg/L |

**Table S27.** Evolutionary treatments and number of replicates at the start and end of the experiment for *Vibrio aestuarianus*.

| Strain ID | Salinity | Copper | Stress | Replicates |
|---|---|---|---|---|
| 12/016 | 0.3M | 0 | Control | 3 |
| 12/016 | 0 | 1.25M | Copper | 3 |
| 12/016 | 0.84M | 0 | Salt | 3 |
| 12/016 | 0.84M | 0.625M | Copper + Salt | 3 |
| U29 | 0.3M | 0 | Control | 3 |
| U29 | 0 | 2.5M | Copper | 3 |
| U29 | 0.84M | 0 | Salt | 3 |
| U29 | 1.125M | 1.25M | Copper + Salt | 3 |

